# *Ormyrus labotus* Walker (Hymenoptera: Ormyridae): another generalist that should not be a generalist is not a generalist

**DOI:** 10.1101/2021.10.26.465982

**Authors:** Sofia I. Sheikh, Anna K.G. Ward, Y. Miles Zhang, Charles K. Davis, Linyi Zhang, Scott P. Egan, Andrew A. Forbes

## Abstract

Several recent reappraisals of supposed generalist parasite species have revealed hidden complexes of species, each with considerably narrower host ranges. Parasitic wasps that attack gall-forming insects on plants have life history strategies that are thought to promote specialization, and though many species are indeed highly specialized, others have been described as generalist parasites. *Ormyrus labotus* Walker (Hymenoptera: Ormyridae) is one such apparent generalist, with rearing records spanning more than 65 host galls associated with a diverse set of oak tree species and plant tissues. We pair a molecular approach with morphology, host ecology, and phenological data from across a wide geographic sample to test the hypothesis that this supposed generalist is actually a complex of several more specialized species. We find 16–18 putative species within the morphological species *O. labotus*, each reared from only 1–6 host gall types, though we identify no single unifying axis of specialization. We also find cryptic habitat specialists within two other named *Ormyrus* species. Our study suggests that caution should be applied when considering host ranges of parasitic insects described solely by morphological traits, particularly given their importance as biocontrol organisms and their role in biodiversity and evolutionary studies.

## Introduction

Parasitism is the most common life history strategy among multicellular organisms (Price 1980, Windsor 1998, Poulin and Morand 2014, Weinstein and Kuris 2016). Species richness of parasitic clades is largely a function of their host ranges: even as many host species (animals, plants, fungi, even other parasites) are parasitized by numerous parasitic species (Price 1977, 2002), many of those parasites are specialized on just one or a few hosts. One hypothesis for why increased specialization tends to evolve among parasites is because new adaptations that increase performance on some host species can consequently reduce the same parasite’s ability to successfully attack other hosts – i.e., fitness tradeoffs favor specialized life histories (Fox and Morrow 1981, Futuyma and Moreno 1988, Jaenike 1990, Agrawal et al. 2010). In parallel, specialization itself has many benefits, including occupation of enemy-free spaces (Bernays and Graham 1988), escape from interspecific / congeneric resource competition (Denno et al. 1995), and reduced time searching for and selecting a host (Bernays and Funk 1999). Because many of the hypotheses for host-specificity are not mutually exclusive, and host environments often vary along multiple relevant axes (including both biotic and abiotic features), multifarious selection may exist for a combination of traits that help maximize fitness within the context of one host/environment (Futuyma and Moreno 1988, Nosil and Harmon 2009).

The biology of parasitic insects and the ways in which they interact with their hosts and surrounding environments may be particularly conducive to the development of specialized host-associations. For example, for many phytophagous insects, the volatile chemical compounds emitted by plants serve as chemosensory cues for locating both preferred habitats and avoiding less favorable ones (Feeny et al. 1989, Berenbaum and Feeny 2008). At the same time, neuronal constraints on the number of host/nonhost signals that can be processed by an insect might often limit the evolution of broad host preferences (Bernays and Minkenberg 1997, Janz and Nylin 1997, Bernays 2001, Egan and Funk 2006). Additionally, while polyphagous feeding strategies can expand food availability in a complex landscape, they also increase the variation and complexity of resources used, which can both increase the likelihood of selecting a lower quality host (Janz and Nylin 1997, Nylin et al. 2000) and weaken the relationship between oviposition preference and offspring performance (Singer 1972, Craig et al. 1989, Itami and Craig 2008). Evidence regarding the magnitude of correlation between preference and performance is equivocal, although, overall, host-specialist females show stronger preference for more suitable, higher quality hosts (Gripenberg et al. 2010).

In addition to navigating a plethora of sensory cues in heterogeneous environments, parasitic insects – especially those with short adult lifecycles – are under strong selection to efficiently find mates (Bush 1975). Because mating often occurs on or near the host for many parasitic insects, as the number of potential hosts increases, the probability of encountering conspecifics may decrease, implying counterbalancing selection for restricted host ranges (Rohde 1979). For insects with short adult life spans, finding acceptable mates on a host plant may also require a degree of synchrony between the phenology of the insect and host. This synchrony plays a crucial role in parasite fitness (e.g., Yukawa 2000, van Asch and Visser 2007) and has been shown to be important in the evolution of new host-associated populations thought to be the progenitors of new specialist species (Komatsu and Akimoto 1995, Forbes et al. 2009). Additionally, behavior (Forister et al. 2012), competition (Futuyma and Moreno 1988), drift (Gompert et al. 2014, Hardy et al. 2016), and standing genetic variation for traits involved in novel host use (Futuyma et al. 1995, Forister et al. 2007) have all been proposed to explain the tendency toward specialization in parasitic insects.

Despite the advantages of specialization, some insect parasites nevertheless appear to act as generalist species (e.g., Thompson 1998). Further, sometime insect species within the same genera are described as apparent specialists while others are described as using many hosts, despite few other obvious differences in their general biology, life histories, the type of hosts they attack, or other dimensions of their niche (e.g., Mitter et al. 1993, Menken 1996). This could portend a real and meaningful difference among congeners (and certainly there may be some circumstances under which specialist clades beget generalist species or vice-versa; Janz et al. 2001, Nosil 2002, Stireman 2005) but might instead signal that a generalist species is not a generalist at all, but rather a conglomeration of several specialist species. After all, most named insect species were originally described solely based on morphological traits, such that the taxonomist who placed too little weight on variation in a particular character or had chosen a genus where morphology was often unhelpful, might not have captured differences relevant to actual reproductive isolating barriers. And indeed, with the advent of molecular ecological studies of parasitic insect species carefully reared from known hosts, there have been several dramatic examples of putative generalists revealed to instead consist of multiple previously obscure specialists (Table 1). We submit that many or most apparent generalist parasitic insects remain unexamined in this respect, and that one may only need to look closer - and with the right tools - at these taxa to reveal hidden specialist clades.

**Table 1.**
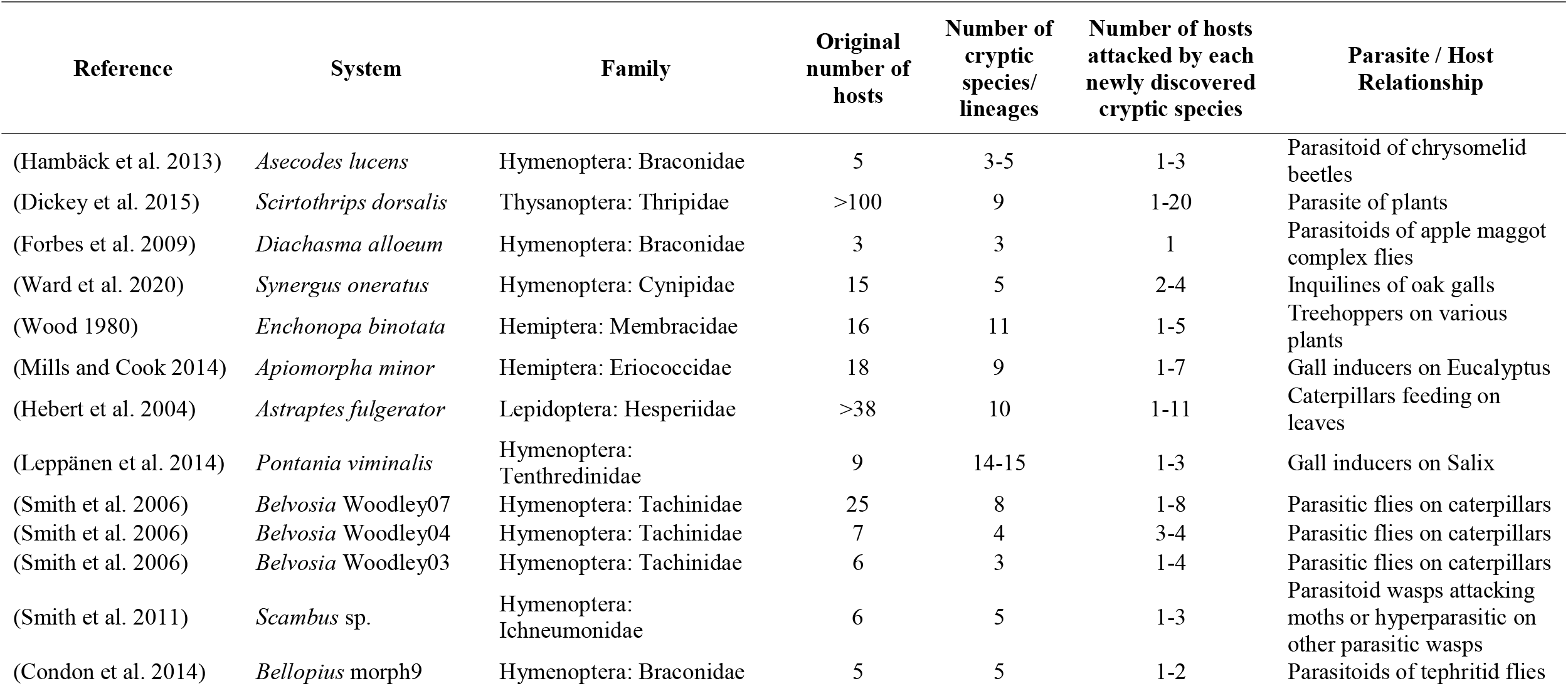
Summary of some previous studies that used an integrative approach to investigate putative generalists and which discovered the presence of several specialist lineages, each with smaller host ranges relative to the original “generalist” species.

One example of a system where one might expect to find – and indeed one often does find – abundant specialization is among the oak gall wasps and their associated communities of hymenopteran parasitoids and inquilines. Oak gall wasps (Hymenoptera: Cynipidae: Cynipini) are a diverse tribe of herbivorous wasps that induce highly structured growths (galls) on oak trees, inside which their progeny feed on the plant tissue. Tight associations between gall wasps and specific oak species indicate that plant volatiles may be involved in tree-host recognition at multiple trophic levels (Germinara et al. 2011), or that oak chemistry influences either female preference for oviposition or offspring performance, or both (Abrahamson et al. 1998, 2003). Among the ∼700 described species of cynipid gall wasps on oaks in the Nearctic (Melika and Abrahamson 2002, Melika and Nicholls 2021), the vast majority are specialized gallers of just one or a few closely related tree species, such that the tree host identity, along with the appearance and location on the tree of the gall itself, is often sufficient to identify the species of gall wasp responsible for the gall (Weld 1957, 1959, 1960, Stone and Schönrogge 2003, Csóka et al. 2005)

Though galls offer protection from predators (Stone et al. 2002, Stone and Schönrogge 2003, Bailey et al. 2009, Ronquist et al. 2015), a taxonomically diverse community of parasitoid and inquiline wasps are nevertheless commonly associated with most galls. These natural enemies often have life histories closely linked with the gall wasp, and/or morphological adaptations that appear essential to overcoming certain gall defensive traits, and/or rearing records that apparently closely track the oak tree species on which the host gall is induced (Ronquist and Liljeblad 2001, Stone et al. 2002, Zhang et al. 2019, Ward et al. 2019, 2020). For many of the reasons explained above, such high levels of specialization should come as no surprise. Galls of greatly varying internal and external morphologies, occurring on specific tissues of specific tree-host species at discrete times during the growing season compose distinct spatiotemporal niches. Galls growing on different tissues are known to release different chemical volatiles (Hayward and Stone 2005), and the high interspecific morphological variation among oak galls may play a role in pattern searching by parasitoid enemies, as well as their respective abilities to parasitize the gall (Bailey et al. 2009). These gall features range from external traits (for example, size, toughness, or nectar secretion) to internal structures (such as the number of chambers and presence of airspace or radiating fibers).

Despite the myriad apparent hurdles to a parasitoid successfully acting as a broad generalist on oak galls, some species have indeed been described as such. For example, *Sycophila biguttata* Swederus (Hymenoptera: Chalcidoidea: Eurytomidae) is described as having 80 host galls associations (Askew et al. 2013). Another parasitoid, *Torymus auratus* Mu□ller, is reported to emerge from 41 gall species (Askew et al. 2013). The expansive host ranges of these species are unexpected, and indeed other gall-associated species in these same genera are considerably more specialized (e.g., *Torymus longiscapus* Grissell, one gall wasp host; (Grissell 1976)). However, both of these enigmatic ultra-generalists, and many others like them, have not been interrogated using a combination of molecular and ecological tools, such that their description as host generalists relies on a shared morphology and little else. Defining the true host ranges of these animals is of interest whatever the outcome. If these really are generalists, we might learn what quirks of biology allow for their cosmopolitan nature among so many closely related specialist congeners. On the other hand, if supposed generalist parasitoids are more often complexes of specialists, among other things, this may have important implications for biocontrol efforts and for understanding patterns underlying the evolution of new diversity.

One such supposed generalist belongs to the genus *Ormyrus* (Hymenoptera: Chalcidoidea, Ormyridae), which are solitary idiobiont ectoparasitoids of various gall-forming insects such as Hymenoptera (Cynipidae, Eurytomidae, Pteromalidae, Agaonidae), Diptera (Cecidomyiidae, Tephritidae), and Coleoptera (Curculionidae) (Gómez et al. 2017). The Nearctic species *Ormyrus labotus* has been recorded from more than 65 named oak gall hosts (Hanson 1987; Supplemental Table 1) from a broad range of tree habitats, gall morphologies, and seasons. In this study, we hypothesize that this apparent generalist parasitoid is actually a complex of several species, each with a much smaller host range. The presence of multiple niche dimensions, each highly variable on its own, which are apparently navigated by this parasitoid makes it ideal for addressing the hypothesis that species like *Ormyrus labotus* are often complexes of several species. Most other species within the genus *Ormyrus* show considerably smaller host ranges than *O. labotus*. For example, *Ormyrus unifasciatrpennis* Girault has been reared from just three gall species; *Ormyrus acylus* Hanson has four host records, three of which are galls in the genus *Callirhytis* Förster (Hanson 1992); and *Ormyrus hegeli* Girault has six recorded host-associations, five of which largely share the same general gall morphology associated with stem tissues. Here, we employ mitochondrial sequence data in conjunction with morphological and ecological data to test whether *O. labotus* is truly an exceptional - and unusual - generalist, or if instead the species consists of several, much more specialized lineages.

## Methods

### Collections, rearing, and morphological identification

Between August 2015 and September 2019, we collected cynipid galls from various oak species across the continental United States (Figure 1). We recorded the date of collection, the geographical location, and host plant species from which galls were collected. The species of gall was determined based on tree host, plant tissue, and gall morphology (Weld 1957, 1959, 1960). Where the gall species could not be immediately determined, we documented a description of the morphology and specific plant tissue upon which the gall was found. We assigned a unique number to represent a collection (representing date, location, tree host, and species of gall), and stored the gall(s) from that collection in an individual container kept in an incubator (SANYO Electric Co. Ltd, Osaka, Japan). The incubator mimicked the external environment in terms of temperature, humidity, and light/dark cycles. We checked the incubator daily and removed any emergent insects for storage in 95% ethanol. We also recorded the collection number and date of emergence. Finally, we used taxonomic keys to identify each non-galler insect to the genus level (Goulet and Huber 1993, Gibson et al. 1997). Out of the ∼150 species of oak galls collected, 51 species reared *Ormyrus*. For wasps in the genus *Ormyrus*, we chose a set of 65 specimens reared from a diverse set of gall hosts and locations that all keyed to *O. labotus*, as well as 29 other *Ormyrus* wasps reared from our collections that did not fit the morphological description of *O. labotus* (Supplemental table 2). We photographed a single forewing and a profile of the body of most wasps used in genetic work and provide them for future reference (Supp Figs. S1–S39). In addition to these, we collected six previously published COI sequences of *Ormyrus labotus*, one of *Ormyrus thymus* Girault (Weinersmith et al. 2020), and 25 of *Ormyrus rosae* (Zhang et al. 2014). *Ormyrus* specimens were keyed to the species level using Hanson (1992) based on pinned specimens, photographs of extracted samples, or both.

**Figure 1:**
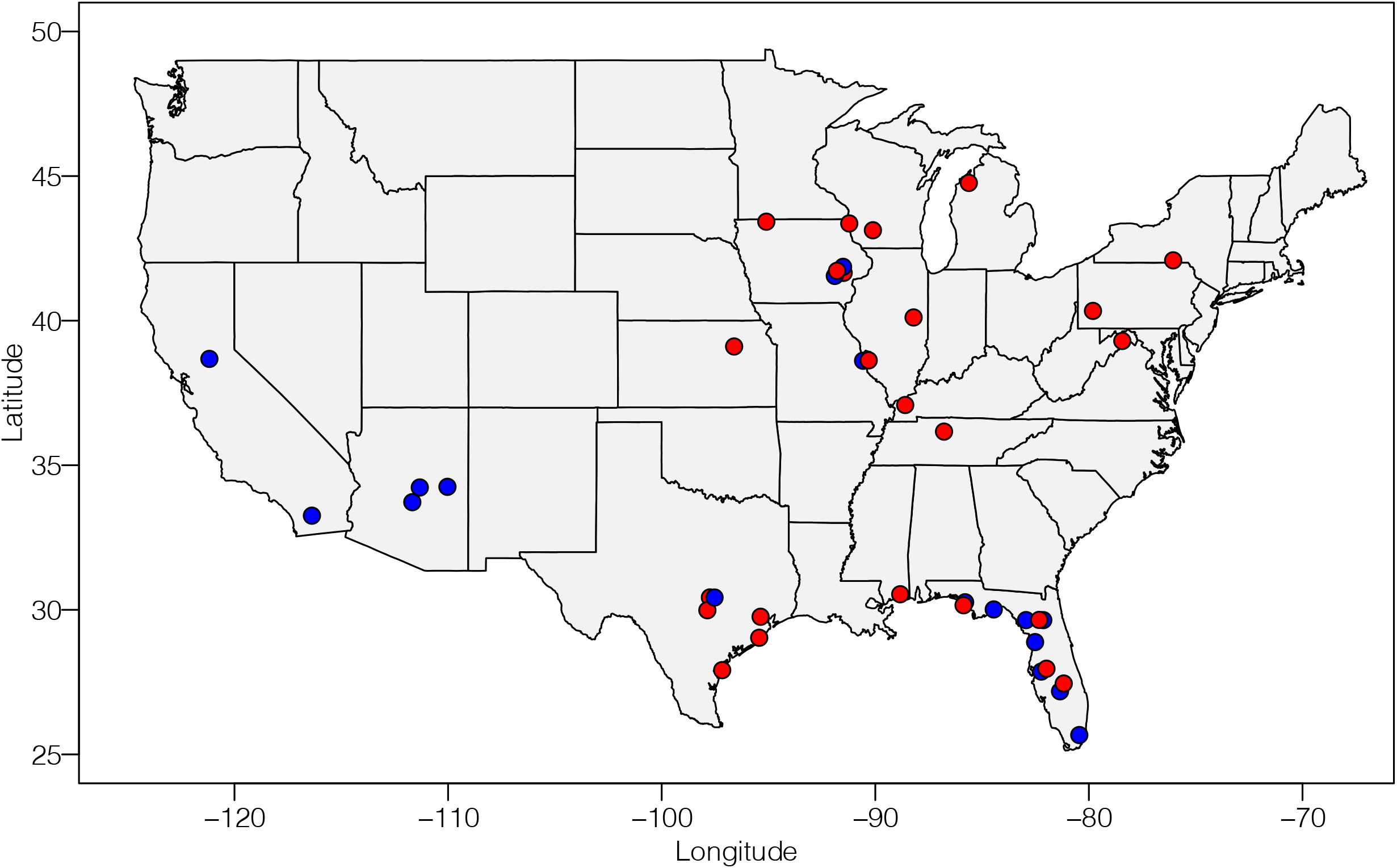
Map of all unique collection regions represented in the COI study. Each dot represents one locale from which at least one collection yielded an *Ormyrus* used in this study. Red dots indicate unique regions from which a gall collection resulted in a wasp that keyed to *Ormyrus labotus*. Blue dots indicate unique geographic regions from which a gall collection reared an *Ormyrus* species other than *O. labotus* (including unidentified individuals and those that did not match any existing descriptions). See Supplemental table 2 for full collection information.

### Sequencing and phylogenetic reconstruction

We extracted DNA using a DNeasy Blood and Tissue kit (Qiagen) from twenty-four of the *Ormyrus* used in this study. For the remaining specimens, we used a CTAB/PCI approach following the methods developed by Chen *et al*. (2010). For all extracted DNA samples, we amplified a ∼650bp region of the mitochondrial COI gene; we amplified ten samples using either lep primers (LepR15’ TAAACTTCTGGATGTCCAAAAAATCA 3’ and LepF1 5’ ATTCAACCAATACATAAAGATATTGG 3’) (Smith et al. 2008), or universal COI primers (COI_pF2: 5′ ACC WGT AAT RAT AGG DGG DTT TGG DAA 3′ and COI_2437d: 5′ GCT ART CAT CTA AAW AYT TTA ATW CCW G 3′) (Kaartinen et al. 2010). For the remaining samples, we used an in-house forward primer Orm_2: 5’ TRG GDG CTC CDG ATA TRG CW 3’ paired with the COI_2437d reverse primer from Kaartinen *et al*.(2010). We designed this forward primer to be more specific to *Ormyrus* COI, while amplifying an overlapping region with the universal COI primers (primer pairs differed by a 26bp region). Sanger sequencing was done in both forward and reverse directions on an ABM 3720 DNA Analyzer (Applied Biosystems, Foster City, CA) in the University of Iowa’s Roy J. Carver Center for Genomics. We then used Geneious v9.1.8 (Biomatters, Inc., San Diego, CA) to prepare consensus sequences, and Geneious Alignment (a built-in aligning program) to generate and manually edit a multiple sequence alignment. jModelTest2 (Darriba et al. 2012) was used to test for the best fitting substitution model for our dataset, and GTR+I+G was selected. For phylogenetic reconstruction, we used two approaches: MrBayes v3.2.7 (Ronquist et al. 2012), which ran two independent analyses using four MCMC chains (one cold, three hot) for 3,500,000 generations, and RaxML v8.2.12 (Stamatakis 2014) with 1000 bootstrap pseudoreplicates.

### Formulation of species hypotheses

To develop working hypotheses about the number of species in our samples, we used a combination of morphological, molecular, and ecological data. After organizing collections by morphological IDs (see above), we then used COI sequence data to group wasps based on sequence similarity using two computational approaches. We first used ASAP, which takes a multiple sequence alignment as input to search for a gap between inter- and intraspecific divergence, and then uses that to sort sequences into putative species groups (Puillandre et al. 2021). The second method was bPTP, a coalescence-based approach that uses a phylogenetic tree as input and estimates the probability of descendant branches being members of the same or different species at each node present in the tree by using branch lengths as a proxy as for substitutions (Zhang et al. 2013). We used the multiple sequence alignment generated in Geneious as input for ASAP, and the Bayesian tree as input for bPTP. For both programs, we used the default settings for all parameters, except we used a Kimura-2-parameter substitution model in ASAP. In cases where there were disagreements between the two computational approaches, we used the more conservative estimate of fewer species. Finally, we used these ASAP and bPTP species results alongside collection information (gall species, gall morphology, tree host, plant tissue, and geography) and rearing data (dates of collection and emergence for each wasp) to generate final estimates of the number of putative species. We calculated the average percent difference between clades using the final putative species assignment for each wasp and the pairwise distance matrix generated for the MSA in Geneious. Where examples were available, representatives of putative species were deposited in the University of Iowa Museum of Natural History (SUI:INS# 39302-39341; Supplemental table 3).

## Results

The Bayesian (Figure S40) and ML (Figure S41) approaches produced similar tree topologies, although relationships among some of the youngest clades differed between inferred trees. In some older nodes, the RaxML tree (Figure S41) produced bifurcating events but with very low bootstrap values. Because our goal was to detect putative species and not definitively resolve the evolutionary histories of those species, we generally did not focus on older nodes. Instead, we relied on well-supported terminal groups to formulate molecular species hypotheses. Both phylogeny approaches produced identical terminal groups.

Figure 2 shows a synthesis of the molecular results, morphological identification, and ecological data used to determine the number of putative species in our sampling (represented by the corresponding clade numbers to the right of the tree). In most cases, bPTP and ASAP results were congruent but there were four exceptions. In three cases where bPTP and ASAP disagreed (clades 15, 26 and 29; Figure 2), we used the more conservative estimate provided by ASAP, as bPTP can sometimes overestimate species number due to geographic structure, or population-level differentiation (Blair and Bryson Jr 2017, Luo et al. 2018). In the fourth case, delimitation methods split clade 31 into two species, but we retained them as a single putative species because they shared several ecological characters (see below).

**Figure 2:**
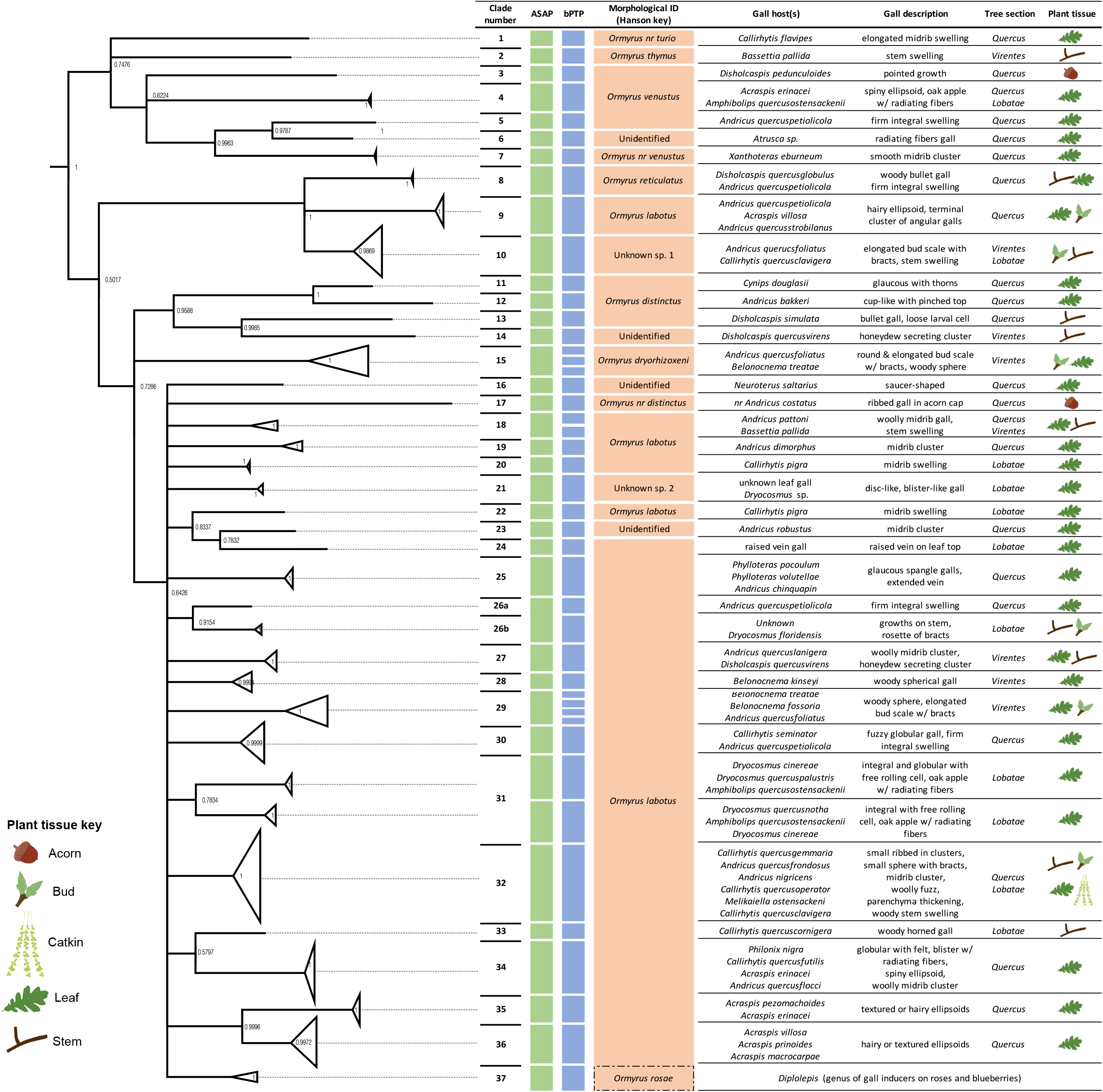
Combination of molecular (COI), morphological, and host data used alongside emergence data (Figure 3) to develop species hypotheses for *Ormyrus* specimens included in the phylogeny. The clade numbers represent putative species based on all the evidence combined. To the furthest left is a cartoon of the mtCOI Bayesian tree (Fig. S40), with nodes collapsed based on our species hypotheses (two exceptions to this are clades 26a/b and 31, which are collapsed based on molecular species delimitation results; see clade descriptions in Results). The two columns to right of the clade number indicate species assignments based on the ASAP and bPTP approaches, respectively. The next column shows the morphological identification of wasps within the corresponding clade based on existing species descriptions. Columns to the right of the morphological ID summarize ecological data for each clade, including gall host, gall morphology, oak tree section, and location of gall tissue.

Emergence data (Figure 3) includes all wasps sequenced in this study, as well as wasps from the same or similar collections that we could definitively link to the same clade (usually through shared morphology). In cases where molecular data showed that wasps from different clades emerged from the same host galls, and where those wasps shared the *O. labotus* morphology, only sequenced wasps that emerged from those were included to avoid confusing emergence timing of wasps from different clades.

**Figure 3:**
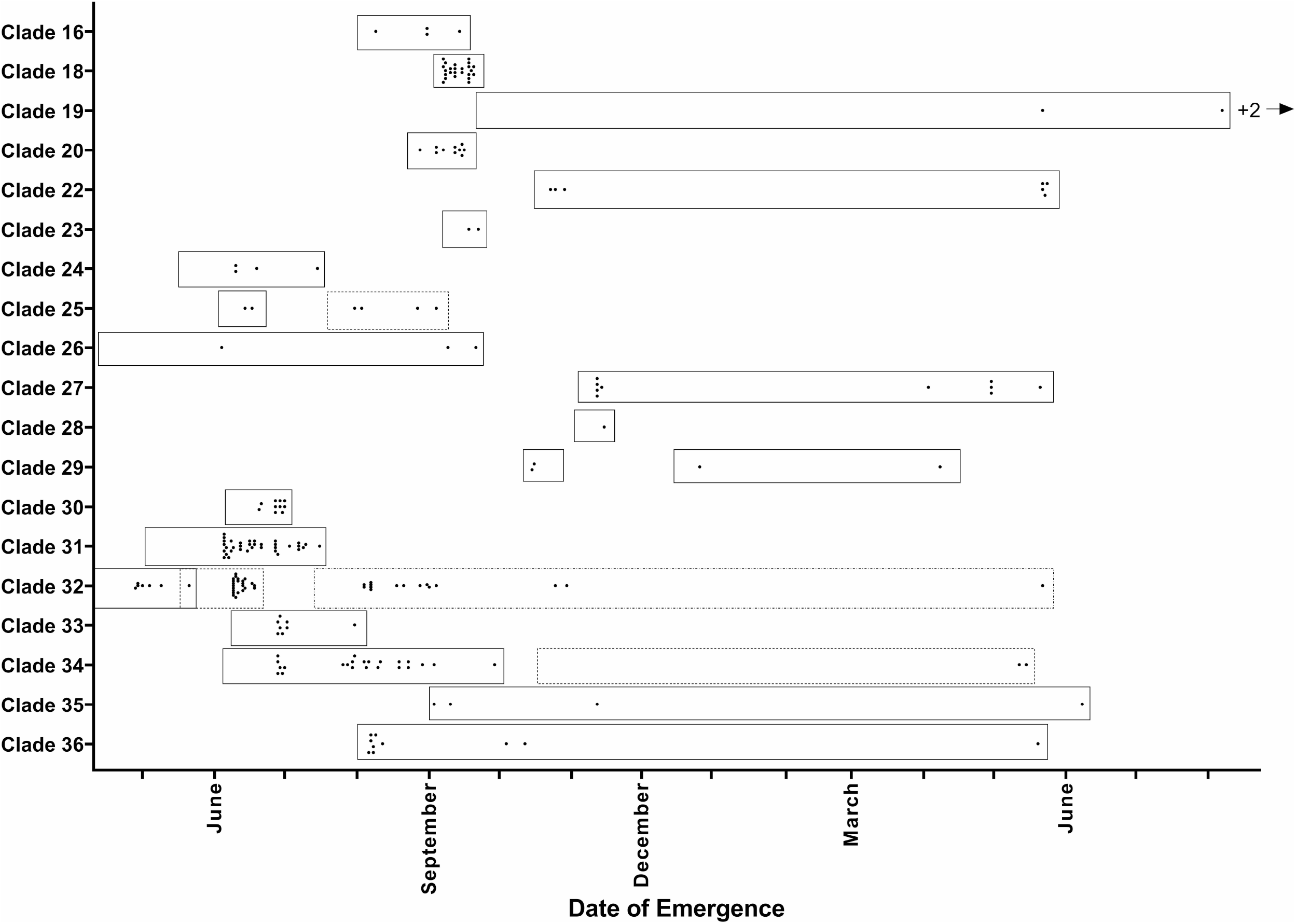
Collection and emergence dates for 19 clades (putative species) of wasps in the “labotus” clade (Figure 2). Excluded are clades 17 (morphologically *O. distinctus*) and 21 (no emergence data available). Dots indicate individual *Ormyrus* wasp emergences; left-most margins of boxes demarcate the earliest date of gall collection from this clade that produced *Ormyrus*. In three cases (Clades 24, 32, 34), boxes with different outline patterns are used to indicate collection events from galls that occur at different times during the year, and which may indicate the use of temporally distributed gall hosts and/or multiple generations. Though galls were collected across several years, each collection was standardized to the year in which the gall first formed. Dates span >1 yr because some insects did not emerge from galls until more than a year after galls were collected. See main text for additional details.

Below we provide a discussion of wasps assigned to each putative species, their species names according to Hanson (1992), host galls, collection locations, dates of collection and emergence (Figure 3), and morphological variation, some of which was only readily apparent after individuals had been sorted based on molecular species hypotheses. In some cases, we refer to percent COI sequence divergence, but for a full table of these percentages for *O. labotus* morphology wasps refer to Table 2. Clade numbers refer to clades in Figure 2. Note, we are not attempting to formally delimit species, but rather to present a synthetic foundation for species hypotheses within this group of wasps.

**Table 2:** Average percentage mtCOI sequence divergence among clades of wasps fitting the morphological description of *Ormyrus labotus* (see Figure 2 for clade assignments and additional information)

|  | Clade 9 | Clade 18 | Clade 19 | Clade 20 | Clade 22 | Clade 24 | Clade 25 | Clade 26 | Clade 27 | Clade 28 | Clade 29 | Clade 30 | Clade 31 | Clade 32 | Clade 33 | Clade 34 | Clade 35 |
| --- | --- | --- | --- | --- | --- | --- | --- | --- | --- | --- | --- | --- | --- | --- | --- | --- | --- |
| Clade 18 | 13.34 |  |  |  |  |  |  |  |  |  |  |  |  |  |  |  |  |
| Clade 19 | 14.71 | 10.02 |  |  |  |  |  |  |  |  |  |  |  |  |  |  |  |
| Clade 20 | 13.53 | 7.35 | 9.63 |  |  |  |  |  |  |  |  |  |  |  |  |  |  |
| Clade 22 | 13.91 | 9.23 | 8.55 | 9.38 |  |  |  |  |  |  |  |  |  |  |  |  |  |
| Clade 24 | 14.60 | 9.88 | 11.8 | 9.41 | 8.97 |  |  |  |  |  |  |  |  |  |  |  |  |
| Clade 25 | 13.95 | 8.45 | 9.63 | 8.27 | 8.22 | 9.84 |  |  |  |  |  |  |  |  |  |  |  |
| Clade 26 | 13.48 | 8.08 | 8.39 | 6.45 | 9.08 | 10.02 | 8.55 |  |  |  |  |  |  |  |  |  |  |
| Clade 27 | 13.84 | 9.20 | 11.23 | 7.84 | 7.67 | 8.22 | 9.37 | 9.14 |  |  |  |  |  |  |  |  |  |
| Clade 28 | 14.81 | 8.54 | 11.81 | 7.15 | 10.27 | 9.80 | 9.22 | 6.59 | 8.75 |  |  |  |  |  |  |  |  |
| Clade 29 | 15.78 | 10.52 | 12.77 | 10.69 | 11.49 | 11.63 | 10.05 | 9.94 | 11.19 | 10.27 |  |  |  |  |  |  |  |
| Clade 30 | 14.61 | 8.56 | 8.59 | 8.19 | 7.84 | 10.86 | 8.56 | 8.04 | 9.04 | 7.58 | 10.74 |  |  |  |  |  |  |
| Clade 31 | 14.53 | 8.93 | 8.73 | 8.17 | 8.98 | 10.42 | 8.65 | 8.27 | 8.69 | 7.94 | 11.55 | 8.00 |  |  |  |  |  |
| Clade 32 | 14.38 | 7.76 | 9.54 | 6.65 | 9.14 | 10.84 | 9.61 | 7.15 | 8.33 | 7.22 | 10.39 | 8.03 | 8.44 |  |  |  |  |
| Clade 33 | 13.72 | 8.34 | 8.00 | 7.11 | 7.41 | 8.80 | 7.87 | 7.00 | 8.62 | 8.47 | 10.38 | 7.68 | 7.54 | 6.73 |  |  |  |
| Clade 34 | 14.92 | 11.30 | 9.87 | 8.41 | 8.13 | 10.30 | 10.32 | 8.79 | 9.77 | 8.57 | 13.01 | 8.17 | 8.51 | 9.27 | 6.78 |  |  |
| Clade 35 | 15.61 | 9.54 | 10.92 | 10.28 | 12.00 | 12.39 | 9.88 | 10.13 | 11.53 | 10.82 | 12.59 | 10.48 | 10.54 | 10.11 | 10.66 | 12.05 |  |
| Clade 36 | 15.46 | 9.91 | 11.59 | 10.33 | 9.73 | 10.67 | 10.24 | 9.82 | 10.26 | 10.36 | 12.75 | 9.07 | 10.90 | 9.87 | 10.29 | 11.23 | 8.39 |

### Ormyrus distinctus

Fullaway (clades 11, 12, 13, 17) is a complex of three to four putative species. The first (clade 11, Fig. S12) was reared from the spiny turban leaf gall, *Cynips douglasii*, collected on valley oak [*Quercus lobata*] in Folsom, CA. The second putative species (clade 12, Fig. S13) was reared from *Andricus bakkeri*, a cup-like leaf gall on scrub oak [*Quercus dumosa*] collected in Borrego Springs, CA. The first two putative species differ in their host ecologies (different gall species and tree hosts), although their different locales and overlapping gall phenology leave open the possibility that they constitute one species with divergent COI haplotypes. The third putative species (clade 13, Fig. S14) was reared from *Disholcaspis simulata*, a round bullet stem gall on scrub oak from Borrego Springs, CA. Wasps from the two clades collected in Borrego Springs had COI sequences diverging by 12.3% and differences in the distribution of setae in the basal cell and speculum. These differences, alongside their having been collected at the same site, support the hypothesis that they are reproductively isolated. Lastly, clade 17 (nr *distinctus*, Fig. S17) was part of the larger unresolved “*labotus*” clade and thus not apparently even sister to these other three putative species. Clade 17 wasps were reared from an acorn gall that matches the description of *Andricus costatus*, a small, ribbed acorn gall on Sonoran scrub oak [*Quercus turbienlla*] from Payson, AZ. All tree hosts noted here are in the white oak section [*Quercus*].

### Ormyrus dryohizoxeni

Ashmead (clade 15, Fig. S16) is represented by four individuals from two gall species: *Belonocnema treatae*, a woody leaf gall on southern live oak [*Quercus virginiana*] from NC, and *Andricus quercusfoliatus*, a cell in elongated bud scales on sand live oak [*Quercus geminata*] from FL. This species is distinctive in its strong blue body, though Hanson (1992) describes collections from the Southwest that are more bronze/green in color. Our collections were from FL and NC and were all the typical blue color.

### Ormyrus reticulatus

Hanson (clade 8, Fig. S9) was reared from two gall species, both on white oak [*Quercus alba*] in Iowa City, IA. One individual represented a collection of *Ormyrus* that overwintered in the round bullet stem gall *Disholcaspis quercusglobulus*; these gall collections were made in April of one year and *Ormyrus* emerged during June and July of the following year. A second individual in this clade was reared from a midrib or petiole swelling gall, *Andricus quercuspetiolicola*. This wasp emerged in July from galls collected in June of the same year.

### Ormyrus rosae

Ashmead (clade 37; no image) is a species previously described by Hanson (1992) and not reared in this study, but sequences from Zhang *et al*.(2014) were used in our analyses. This species does not attack galls on oaks, having been reared exclusively from cynipid galls on roses [*Rosa*] and pteromalid galls on blueberries [*Vaccinium*]. Hanson (1992) recommends host associations as the most reliable character for distinguishing these wasps from *Ormyrus labotus*.

### Ormyus thymus

Girault (clade 2, Fig. S2) is represented by one individual reared from a crypt stem gall *Bassettia pallida* on sand live oak [*Q. geminata*] in Inlet Beach, FL. In addition, a single, enigmatic, host is recorded for this species – seeds of *Bucida cucides* (Combretaceae) in Belize, but adult wasps have previously been collected from Florida, California, and Georgia (Hanson 1992). This collection was previous reported in Weinersmith *et al*. (2020).

### *Ormyrus* nr. *turio*

Hanson (clade 1, Fig. S1) was reared in Oxford, IA from *Callirhytis flavipes*, a multi-cell midrib swelling on the leaves of bur oak [*Quercus macrocarpa*]. These wasps were morphologically closest to *O. turio*, a species previously recorded from only one gall host, *Bassettia ligni* in California (Hanson 1992). If this is the same species it would appear to have a particularly large geographic range, but we caution that some characters do not exactly match the Hanson (1992) description of *O. turio* (e.g., head not obviously subquadrate, antennae not strongly clavate). Though they are closest to *O. turio*, their next best morphological match is to *O. labotus* - though the setae of their cubital veins do not continue across the base of their speculum (i.e., they have an “open” speculum).

### Ormyrus venustus

Hanson (clades 3, 4, 5, 7) is a complex composed of between two and four putative species. The first species (clade 3, Fig. S3) is represented by one individual that emerged in October from *Discholcaspis pedunculoides*, an acorn gall on Sonoran scrub oak [*Q. turbienlla*] collected in Rio Verde, AZ. The second species (clade 4, Fig. S4) was reared from two gall hosts in Iowa City, IA: *Acraspis erinacei*, a leaf gall with spines on white oak [*Q. alba*], and *Amphibolips quercusostensackenii*, a round integral leaf gall with radiating fibers, on post oak [*Quercus stellata*]. The third species (clade 5, Fig. S5) is also represented by one individual reared from *A. quercuspetiolicola* on post oak [*Q. stellata*] in Austin, TX. The fourth species (clade 7– *nr venustus*, Fig. S7-8) consists of two individuals, both reared from the same collection of *Xanthoteras eburneum*, a leaf gall on gambel oak [*Quercus gambelii*] in Show Low, AZ. Clade 7 wasps differed from the Hanson (1992) description of *O. venustus* in having their scutella diagonally strigate. Because these four putative species were collected from different geographic locations, it remains possible that they are a single widely distributed species with highly variable COI (ranging from an average of 10.3 to 14% between clades). However, apparent morphological differences (e.g., variation in length of female tergite eight (Figs. S4, S5, S7), sculpturing of scutellum) complement genetic and ecological differences in supporting the hypothesis that at least clade 7, if not all four clades, represent distinct species.

### Unidentified *Ormyrus*

(clades 6, 14, 16, 23) are species that we were unable to identify morphologically. The lack of identification is either because they were represented in our collections only by males (Hanson (1992) provides only a key to females), or because physical specimens were not available and important characters were obscured in photos of extracted specimens. The first species (clade 6, Fig. S6) was reared from a leaf gall with internal radiating fibers in the genus *Atrusca* on Sonoran scrub oak [*Q. turbienlla*]. The second species (clade 14, Fig. S15) is represented by one individual reared from *Disholcaspis quercusvirens*, a stem gall on southern live oak [*Q. virginiana*] from Gainesville, FL. The third putative species (clade 16) emerged from *Neuroterus saltarius*, a saucer-shaped leaf gall on white oak [*Q. alba*] in IA. Finally, the last unidentified putative species in the mtCOI tree (clade 23, Fig. S23) emerged from *Andricus robustus*, a midrib (leaf) cluster gall on post oak [*Q. stellata*] from St. Louis, MO.

Two **unknown species of *Ormyrus*** were present in our sampling. These differ from the unidentified *Ormyrus* in that while physical specimens and detailed photos were available to discern key traits, the unknown species did not fit any existing species descriptions based on Hanson (1992). The first species (clade 10, Fig. S11) was reared from two gall hosts. One host was *A. quercusfoliatus*, a bud gall on two species of live oak: sand live oak [*Q. geminata*] in St. Teresa, FL and southern live oak [*Q. virginiana*] in Hammond, Citrus, and Lithia Springs in FL. The other gall host of this species is *Callirhytis quercusclavigera*, a spring stem gall on scarlet oak [*Quercus coccinea*] in Gainesville, FL. All five *Ormyrus* from *A. quercusfoliatus* overwintered in the gall. The second unknown species (clade 21, Fig. S21) emerged from two unidentified leaf galls on sand laurel oak [*Quercus hemisphaerica*] in Florida. Exact emergence dates were not collected for clade 21 wasps, but they were collected in late March.

### Ormyrus labotus

Walker is a complex of 16-18 putative species, which we refer to by their clade assignments in Fig. 2 based on our bPTP and ASAP results:

Clade 9 wasps (Fig. S10) emerged from three gall species: *Andricus quercusstrobilanus*, a cluster of stem galls on swamp white oak [*Quercus bicolor*], *Acraspis villosa*, a spiny leaf gall on bur oak [*Q. macrocarpa*], and *A. quercuspetiolicola*, a midrib/petiole swelling on swamp white oak [*Q. bicolor*]. All galls were collected in Iowa City, IA, and *Ormyrus* from *A. villosa* overwintered, emerging the following summer after the gall was induced. Wasps in this clade were the only “*Ormyrus labotus*” wasps that did not group in the larger *labotus* clade (Fig. 2), though we caution overinterpretation of evolutionary relationships from a single gene tree.

Clade 18 wasps (Fig. S18) were reared from *Andricus pattoni*, a woolly midrib cluster gall on the leaf of a post oak [*Q. stellata*] in Peducah, KY, and from *Bassettia pallida*, a stem swelling gall on sand live oak [*Q. geminata*] from Inlet Beach, FL.

Clade 19 wasps (Fig. S19) included wasps reared from *Andricus dimorphus*, a midrib cluster gall on bur and dwarf chinquapin [*Quercus prinoides*] oaks in Lansing, IA and Konza, KS, respectively. Emergences from both gall types occurred in the calendar year after galls were collected (Figure 3).

Clade 20 wasps (Fig. S20) were reared from *Callirhytis pigra*, a midrib leaf swelling on red oak [*Quercus rubra*] in Nashville, TN. Clade 22 wasps (Fig. S22) were also reared from *C. pigra*, but on black oak [*Quercus velutina*] in Vestal, New York. Wasps from clades 20 and 22 were collected from different tree hosts and emerged a month and half apart from each other (Figure 3), suggesting some degree of temporal isolation. However, they were also collected in different geographic regions, leaving open the possibility that their COI sequence divergence (9.4%) is due to isolation by distance, and in which case their temporal differentiation may represent two generations using different gall hosts. Sequencing of additional loci and collection of additional specimens from a wider geographic area may help clarify the status of these two clades as one or two distinct species.

Clade 24 wasps (Fig. S24) emerged from June to July from an unidentified leaf vein swelling on pin oak [*Quercus palustris*] collected in May in Iowa City, IA.

Clade 25 wasps (Fig. S25) were reared from three gall species: *Andricus chinquapin*, an extended vein on the leaf, and *Phylloteras volutellae* and *Phylloteras pocoulum*, two spangle leaf galls collected in early fall. All three galls were collected from leaves of swamp white oak [*Q. bicolor*] in Iowa City, IA. Wasps from *A. chinquapin* galls emerged in June, while those from the two *Phylloteras* galls emerged from July-September (Figure 3). It is possible that these may represent two generations using two types of galls on the same trees at different times during the year, but further molecular and ecological work is required.

Clade 26 wasps comprise two subclades (clade 26a, Fig. S26 and clade 26b, Fig. S27) reflecting their assignment by bPTP into two molecular species (Figure 2). Because ASAP did not separate them into two species, and because they were collected in TX and FL, respectively, COI differences could be due to geographic isolation and so we conservatively group them together into one putative species. Sub-clade 26a is represented by one individual reared from *A. quercuspetiolicola* on post oak [*Quercus stellata*] in Austin, TX. Sub-clade 26b is represented by two individuals: one reared from *Dryocosmus floridensis*, a bud gall on laurel oak [*Quercus laurifolia*] from Gainesville, FL, and the other from an unidentified fuzzy pink gall on overcup oak [*Quercus lyrata*] from Otter Springs, FL.

Clade 27 wasps (Fig. S28) were reared from *Andricus quercuslanigera*, a woolly leaf gall collected in Kyle, TX on live oak [*Q. fusiformis*], and *D. quercusvirens*, a stem gall collected in Gainesville, FL on southern live oak [*Q. virginiana*].

Clade 28 wasps (Fig. S29) were reared from *Belonocnema kinseyi*, a woody spherical leaf gall, on southern live oak [*Q. virginiana*] in Houston, Lake Jackson, and Ingleside, TX. Clade 29 (Fig. S30) was reared from three gall species: *Belonocnema treatae* on southern live oak [*Q. virginiana*] in MS, *Belonocnema fossoria* on sand live oak in FL, and *A. quercusfoliatus*, on both southern [*Q. virginiana*] and sand live oak [*Q. geminata*] in FL. While all three *Belonocnema* gall wasp species share gall morphology, they show strong differentiation along geography – a common pattern for species distributed around the U.S. gulf coast (Zhang, Egan, et al. 2021, Zhang, Hood, et al. 2021). The *Ormyrus* reared from these galls are separated into two species in Figure 2 because they differ by an average of 10.3% in COI sequence and clade 29 wasps have a distinct striped patterning on their lateral metasoma (Fig. S30), earning them the moniker “tigermorphs” in our working group. However, because clade 28 wasps were collected in TX and clade 29 wasps were collected in MS and FL, differences in COI sequence and body color could also represent geographic distances, so we count them here as representing either one or two species.

Clade 30 wasps (Fig. S31) were reared from two gall species: *A. quercuspetiolicola* on bur [*Q. macrocarpa*] and white oak [*Q. alba*] from Oxford and Iowa City, IA, and *Callirhytis seminator* (detachable woolly stem gall) on white oak [*Q. alba*] from Iowa City, IA. Both galls listed for this species were collected in early June with *Ormyrus* emergences in late June.

Clade 31 wasps (Fig. S32) were split into two species by both molecular species delimitation methods; however, in both clades (Figure 2) were reared from two of the same gall species (*Dryocosmus cinereae and A. quercusostensackenii*) on trees in the red oak section, so to be conservative we call them a single putative species. Each group has one additional gall species different from the other, but these also (*Dryocosmus quercuspalustris* and *Dryocosmus quercusnotha*) are both of the same genus, induce integral leaf galls, have internal free space (free rolling cell and radiating fibers, respectively) similar to other two galls within the host range, share phenology (May –June, Weld 1959), and are found on trees in the red oak section. There were also no obvious differences between the two groups in their emergence data (Figure 3).

Clade 32 wasps (Fig. S33) were represented by ten individuals reared from six gall species across six oak species (two in the white oak section and four in the red oak section) in IA, MI, and FL. Gall hosts include: *Andricus nigricens* (midrib cluster), *Andricus quercusfrondosus* (bud gall with bracts), *Melikaiella ostensackeni* (parenchyma thickening), *Callirhytis quercusgemmaria* (nectar-secreting stem gall), *Callirhytis quercusoperator* (woolly catkin gall), and *C. quercusclavigera* (woody stem swelling). Though these are a relatively diverse set of hosts, they may represent wasps in this clade specializing on different hosts at different times during the year. For instance, the earliest wasps emerged in April and May from *A. quercusfrondosus* bud galls collected in March and April (Figure 3, Clade 32 solid box), then a second set of wasps emerged in June from *A. quercusoperator* catkin galls collected in late May (Figure 3, Clade 32 dashed box), and finally a third set of galls emerged from August to October (with one wasp emerging the following May) from *M. ostensackeni* and *C. quercusgemmaria* galls collected in July and August, respectively. We do not have emergence data for the single wasp reared from *C. quercusclavigera* in Florida.

Clade 33 wasps (Fig. S34) were reared from *Callirhytis quercuscornigera* (horned woody stem galls) on pin oak [*Quercus palustris*] in St. Louis, MO and St. Peters, MO. These galls were collected in early June and wasps emerged from late June to late July of the same year, though one wasp emerged in May of 2018 more than 1.5 years after its *C. quercuscornigera* host gall was collected in September of 2016 (this wasp is not shown on Figure 3).

Clade 34 wasps (Figs. S35-36) were reared from four gall species, all leaf galls, and all on white oak [*Q. alba*] from IA, IL, WI, and WV. The gall hosts were *A. erinacei* (spiny leaf gall), *Andricus quercusflocci* (woolly leaf gall), *Callirhytis quercusfutilis* (integral leaf blister gall), and *Philonix nigra* (globular gall with felt). All gall hosts except *C. quercusfutilis* are detachable leaf galls. Most wasps in our collections emerged from late June to early September from galls collected between early June and early September (Figure 3, Clade 34 solid box). Two wasps emerged the following May from *A. erinacei* and *A. labotus* galls collected from leaf litter (Figure 3, Clade 34 solid box).

Clade 35 wasps (Figs. S37-38) were reared from two leaf galls, *A. erinacei* and *Acraspis pezomachoides* (textured globular or ellipsoidal galls) on white oak [*Q. alba*] from IL, KY, PA, and NY. All host galls for this putative species were collected in September and October, with emergences in September and November, and one in June of the following year.

Clade 36 wasps (Fig. S39) were reared from three gall species, which were all leaf galls in the genus *Acraspis* from IA and IL: *Acraspis macrocarpae* (morphologically like *A. pezomachoides* but on bur oak – *Q. macrocarpa*), *Acraspis prinoides* (globular with cone-shaped projections) on chinquapin oak [*Quercus muehlenbergii*], and *Acraspis villosa* (similar to *A. erinacei* but on bur oak – *Q. macrocarpa*). All host galls were collected between August and September and wasps emerged between August and October, with one wasp emerging in May of the following year. Though it is possible that clades 34-36 are one species with three extremely divergent haplotypes (8-12%; Table 2), their use of different host gall species and sympatry at one site (Urbana, IL) suggests that they are reproductively isolated. Clade 34 and 35 wasps were both reared from one of the same hosts (*A. erinacei*).

In total, we find evidence for 31–36 species of *Ormryus* present in our samples, including 17–19 species that matched the morphological description of *O. labotus*. These included one species, *Ormyrus rosae* Ashmead, which Hanson (1992) previously separated from *O. labotus* based primarily on its host association with galls on roses (Rosaceae) and blueberries (Ericaceae). Thus, we find a total of 16–18 oak-gall-associated putative species fitting the morphological description of *O. labotus*. Additionally, we found three to four putative species matching the morphological identification of *O. distinctus* (clades 11– 13 and 17; Figure 2), two to four species matching *O. venustus* (clades 3–5 and 7; Figure 2), four species that we were unable to determine a morphological identification for (clades 6, 14, 16, 23; Figure 2), and two species that do not match any existing species descriptions (clades 10 and 21, Figure 2). The other *Ormyrus* species represented in our sampling included: *O. dryohizoxeni, O. reticulatus, O. thymus*, and *O*. nr *turio* (clades 15, 8, 2, 1; Figure 2).

## Discussion

We find that the supposed generalist *O. labotus* is apparently a complex of several species, each with a far narrower host range than had previously been reported (Hanson 1992, Noyes 2021). Our combined molecular and ecological analysis yielded 31-36 putative species present across all samples, including 16– 18 species nested within larger clades of wasps that all ran to *O. labotus* in the Hanson (1992) key (Figure 2). Though we were not explicitly testing for differentiation within other *Ormyrus* species, we also discovered apparent cryptic species present in each of two other morphologically defined species, *O. distinctus* and *O. venustus*, each again with a narrower range of hosts than previously reported. While a formal taxonomic revision of the genus is beyond the scope of this work, we note that there appear to be some emergent morphological, phenological, and host-associated differences that may prove useful in delimiting and describing some of these putative new species.

The discovery of morphologically cryptic, host-specific diversity in insect taxa generally, and oak-gall-associated natural enemies specifically (e.g., Kaartinen et al. 2010, Nicholls et al. 2010), is not unusual. For example, the Palearctic species *Ormyrus pomaceus* Geoffroy has been reported from more than 56 species of cynipid galls, but molecular and ecological data suggests *O. pomaceus* is a complex of cryptic species (Kaartinen et al. 2010, Gómez et al. 2017). Similarly, Zhang *et al*. (2014) found two species matching the description of *Eurytoma spongiosa* Bugbee (Hymenoptera: Chalcidoidea: Eurytomidae) that differ in their host associations. In Ward *et al*. (2020), five of eleven previously described species of *Synergus* (Hymenoptera: Cynipidae: Synergini) were shown to be complexes of more than one putative species, each with a small host range and specializing on galls of similar morphologies restricted to a single oak tree section.

### Host ranges, host shifts, and evolution in Ormyrus

Far from being generalists in their associations, most *Ormyrus* species were reared from galls on just one or two tissue types and from a single oak tree section (Figure 2). In clades where we observe relatively larger hosts ranges (e.g., clades 31 & 34), hosts tended to be similar in ecology, phenology, and/or gall morphology, and/or there was some evidence that different hosts were used across the course of a year (e.g., clade 32). Additionally, we find several clades in which *Ormyrus* collected in different locations were assigned to the same species (e.g., clades 30, 35, and 36), as well as several sympatric wasps differing only in their host associations (Supplemental table 2) indicating that gene flow is not strongly restricted by geography but rather more by ecology (dimension of the host). This further suggests that host associations may have been important in *Ormyrus* diversification, though evaluating phylogenetic questions will require more than a single gene.

In Hanson’s (1992) study of genus *Ormyrus*, he noted that some of his named species attacked galls with shared characters – for instance, shared morphology and plant organ among the gall hosts of *O. hegeli*, or common gall wasp host genera for both *O. acylus* and *O. reticulatus*. In discovering many more putative species, each with narrower host ranges, we hoped that we might reveal a specific common axis of adaptation (e.g., host galler species, gall morphology or phenology, tree section or species) that was most important for defining host ranges in *Ormyrus* wasps. However, beyond finding that each *Ormyrus* species is specialized on a narrow range of hosts, we note no single unifying theme. Like Hanson (1992), we find examples of species with multiple hosts that share similar characters, but the apparent axes of specialization are highly variable (e.g., clades 35 and 36, reared only from gallers in genus *Acraspis*; clade 25, reared only from leaf galls from swamp white oak [*Quercus alba*]; clade 32, with three generations possibly specialized on particular gallers (or plant tissues), but found on trees in two different sections of the oak family). It may therefore be that while each species specializes on one or more ecological dimensions of the host, no single dimension is paramount. In other words, specialization may be opportunistic, with adaptations to different aspects of the gall environment varying across lineages or even among populations. Alternatively, it may be that we have failed to analyze one or more important aspects of *Ormyrus* specialization, or that this large sample is not yet sufficiently complete for common patterns to emerge.

While we were able to formulate species hypotheses and investigate the relevance of various niche dimensions in the host ranges of *Ormyrus* species, resolution below the species level is fairly poor and relationships among putative species remain largely uncertain (Figure 2; Supplemental Figures 41 and 42). Future studies are needed to resolve relationships among these putative species and evaluate evolutionary patterns for *Ormyrus*, including whether shifts to new hosts are correlated with changes in particular niche dimensions, while others are more likely to be conserved. We see early hints of ecological differences between well-resolved sister clades. For example, some appear to reflect conservation of galler genus alongside shifts to different tree hosts (e.g., clades 35 and 36 attack *Acraspis* gallers on different trees; Figure 2) while others apparently undergo major shifts between tree section, tree species, and galler host (e.g., clades 33 and 34; Figure 2). However, we caution that the COI gene tree is not necessarily a reflection of the overall species tree. Additionally, our sampling of North American *Ormyrus* was haphazard in its approach, such that many additional species likely remain to be discovered. Despite these cautions, we submit that *Ormyrus* speciation events seem universally correlated with changes in host association. With additional collections and a multilocus phylogenetic approach, this should prove a rewarding system in which to evaluate hypotheses of host-shift speciation or other manifestations of ecological speciation.

One additional caveat to our study is that it relies strongly on mitochondrial COI to infer species hypotheses, which various authors have cautioned against and for various reasons (e.g., Funk and Omland 2003, Zamani et al. 2020). However, this approach has demonstrated to be a powerful tool in detection of otherwise cryptic species, especially when paired with ecological data, behavioral assays, and/or statistical tools (e.g., examples in Table 1; Forbes et al. 2012, Duran et al. 2019). Our COI analysis indicates considerable genetic divergence (6.5% – 15.6% among wasps with *O. labotus* morphology; Table 2), and in most cases where molecular methods suggested that collections were from different species, these molecular differences were correlated with ecological differences.

### Hidden specialists – Why it matters

Why does it matter that cryptic clades of small parasitic insects might often be lumped together as single species? Some reasons are economic. Parasitic insects include many forest and agricultural pests, as well as the natural enemies of those same pests. Discerning host ranges and patterns of host-use for insect pests and their parasitoids is useful for designing effective biocontrol strategies to regulate parasitic insect pests (Nicholls et al. 2018). For example, were *O. labotus* to be considered for biological control of an invasive oak-galling wasp, selecting a population to draw from based on a concept of *O. labotus* prior to this paper would likely result in failure (1/N; where N is the number of new OTUs), whereas we now know that host ranges are narrow and careful selection from appropriate sources would be important. Several prominent examples demonstrate the importance of this idea to parasitoids of insect agricultural pests (e.g., Heraty et al. 2007, Forbes et al. 2009, Hood et al. 2015, Paterson et al. 2016, Seehausen et al. 2020)

From a basic science standpoint, clarifying the putative axes along which lineages specialize, as well as which components play crucial roles in species diversification will improve our ability to study how parasites evolve. Many studies have used host range data to ask synthetic questions about the relationship between host specialization and diversification (Winkler and Mitter 2008, Armbruster and Muchhala 2009, Novotny et al. 2012, Ebel et al. 2015, Forbes et al. 2017). Conclusions arising from such work are highly dependent on both correct species delimitation and the completeness of host range investigation.

Incomplete understanding of host ranges might also hinder our ability to study actual generalists when they do occur. There are good theoretical arguments for why some parasitic species may settle on a generalist approach (e.g., Futuyma and Moreno 1988). Compared to occupying a narrow ecological distribution, a broader niche allows for diet mixing between different life stages, and a nutritionally balanced diet (MacFarlane and Thorsteinson 1980, Barbosa et al. 1986, Bernays and Minkenberg 1997). Generalism also confers the ability to bet hedge against changing environments by maintaining access to alternative hosts (Funk and Bernays 2001). But care should be taken to ensure the insect systems used to infer the ecological conditions and genetic/morphological tools that enable generalist lifestyles are not actually complexes of cryptic specialists. Functional studies, behavioral assays, morphometric analyses, and transcriptomic work with the potential to elucidate the processes that result in different feeding strategies require *true* generalists to compare against their closely related specialist counterparts.

Finally, in this current conservation crisis, work to refine our understanding of biodiversity has become more critical than ever. Across the tree of life, different species differ in how they interact with their environments, such that a species that is lost from a system cannot be easily substituted with even a closely related species. Integrative taxonomic efforts have regularly discovered new species even within large, charismatic taxa (e.g., giraffes: Fennessy et al. 2016), but some of the most species-rich organismal groups are also among the most understudied. In particular, parasitic wasps (a group to which *O. labotus* belongs) are likely the most species-rich group in class Insecta (Noyes 2012, Forbes et al. 2018), though also among the most resistant to taxonomic classification due to the “taxonomic impediment” of too few taxonomists and too many species (Taylor 1983, Giangrande 2003). If many parasitic wasp “species” are actually complexes of several more specialized species, then they are also more susceptible to extinction because their survival relies on the availability of a smaller number of host species.

### Conclusions

We submit that this study, alongside a steadily accumulating list of similar studies showing cryptic specialization among parasitic insects (Table 1), validate a need for a higher standard of evidence to call a parasitic insect species a “generalist”. To describe a parasite as a generalist should require 1) that molecular and ecological data allow for a strong rejection of the hypothesis that there are multiple reproductively isolated lineages and 2) ecological studies showing use of many hosts by a single lineage. We of course recognize that definitions of “generalist” may vary. In the current study for instance, wasps in clade 32 were reared from six oak gall species on six species of oak. Is this a generalist species because it attacks several different hosts? Or is it a specialist because it attacks only a limited number of hosts from among a much larger available pool? Regardless of the descriptor, what matters is that for various considerations – from economic to basic biology – we must work to increase accuracy in detection of reproductively isolated species and description of their respective host ranges and ecologies.

Finally, we strongly emphasize that this work is not a criticism of taxonomy or taxonomists (and especially not of Hanson (1992), which we consider a masterful study of a borderline intractable genus of tiny wasps). Rather, we caution that host ranges reported in taxonomic publications might sometimes (or often) represent amalgamations of host use data from across several parasitic species. Strictly morphological descriptions of species are, like anything in science, subject to revision, and many elegant examples (Table 1) demonstrate that species hypotheses improve with added molecular, ecological, and other biological data. Indeed, corroboration of morphological methods with molecular data, ecological studies, and natural history records across the geographical range of a species is already standard practice in taxonomy today, but such work is expensive and requires broad methodological expertise. Further, taxonomy, including training of new taxonomists, is increasingly underfunded, such that there are fewer experts to conduct such studies. Funding for integrated taxonomy, including creating training opportunities and permanent positions for taxonomists should be prioritized in the biological sciences.

## Acknowledgements

We thank several individuals who helped with gall collections, including Robert Busbee, Robin Bagley, Will Carr, Sarah DeLong-Duhon, Alaine Hippee, Glen Hood, Kyle McElroy, Daniel McGarry, Kevin Neely, Jim Ott, Monzer Shakally, Eric Tvedte, Joseph Verry, Kelly Weinersmith, and Caleb Wilson. Funding for collections was provided to A.K.G.W. and S.I.S. from The Center for Global and Regional Environmental Research at the University of Iowa, and to A.A.F by a James Van Allen Natural Sciences Fellowship. YMZ was funded by the Oak Ridge Institute for Science and Education (ORISE) fellowship. Mention of trade names or commercial products in this publication is solely for the purpose of providing specific information and does not imply recommendation or endorsement by the USDA. USDA is an equal opportunity provider and employer.

## Author Contributions

S.I.S., A.K.G.W., and A.A.F. designed the study. All authors made collections and/or reared animals. S.I.S., A.K.G.W., Y.M.Z., C.D., and L.Z. obtained the sequence data. S.I.S. and A.A.F. conducted the analyses, and S.I.S., Y.M.Z. and A.A.F. wrote the manuscript. All authors contributed to revisions and approved the final manuscript.

## Notes

### Competing Interest Statement

The authors have declared no competing interest.

### Summary of Updates

Found a small error in the abstract that required correction.

