## Supplemental Figures S1-S39 for "*Ormyrus labotus* Walker (Hymenoptera: Ormyridae): another generalist that should not be a generalist is not a generalist"

**Figure S1. *Ormyrus* near *turio* (clade 1):** 881-013-7, female from *Callirhytis flavipes* on *Quercus macrocarpa* – Oxford, IA.

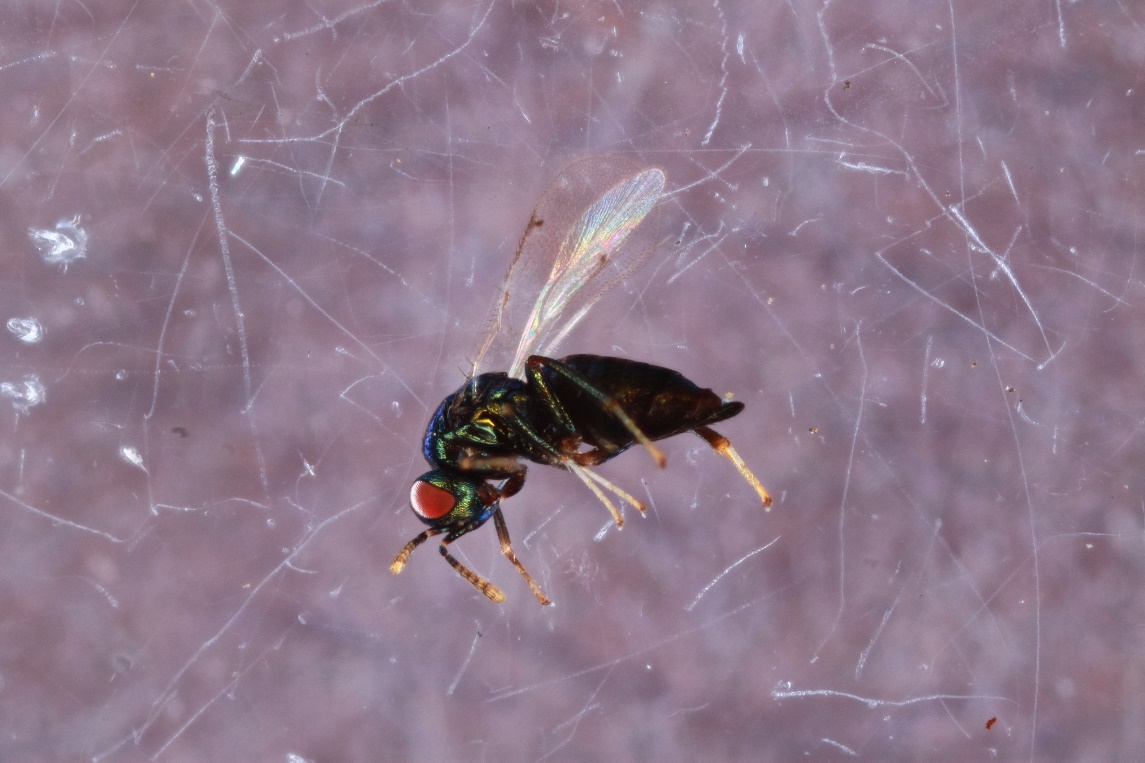

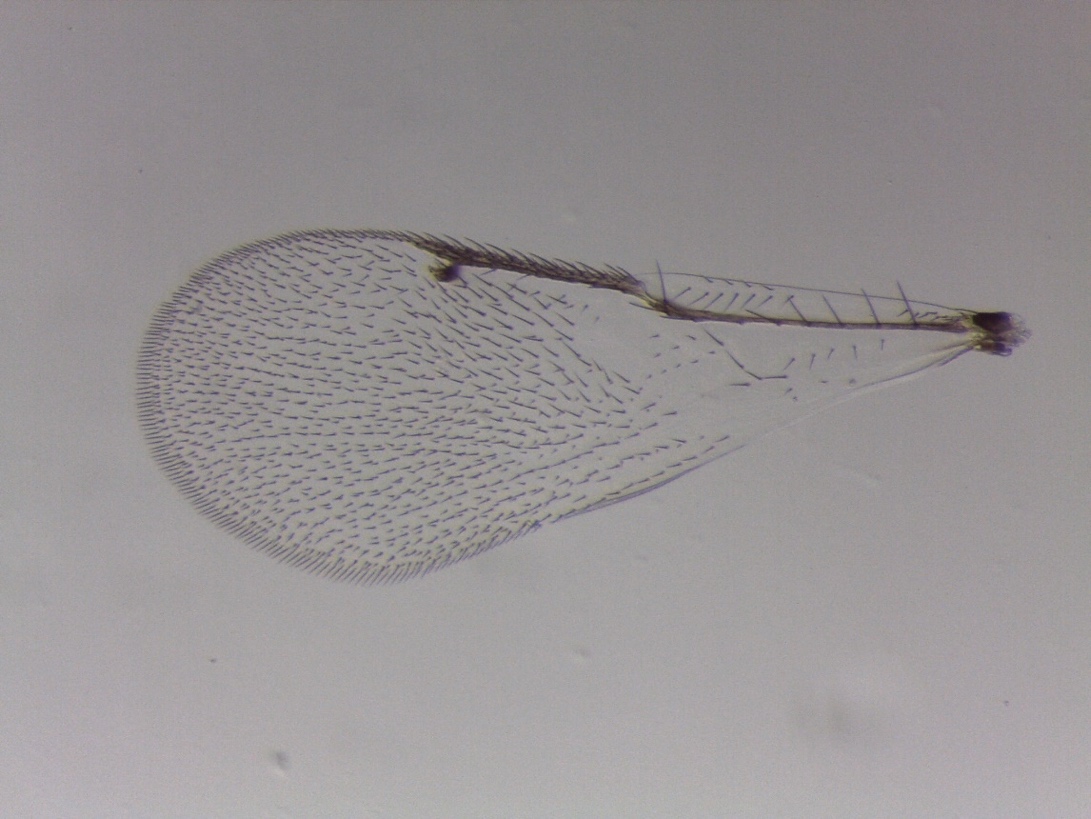
**Figure S2. *Ormyrus thymus* (clade 2):** KW004, male from *Bassettia pallida* on *Quercus geminata* – Inlet Beach, FL (Weinersmith *et al*. 2020).

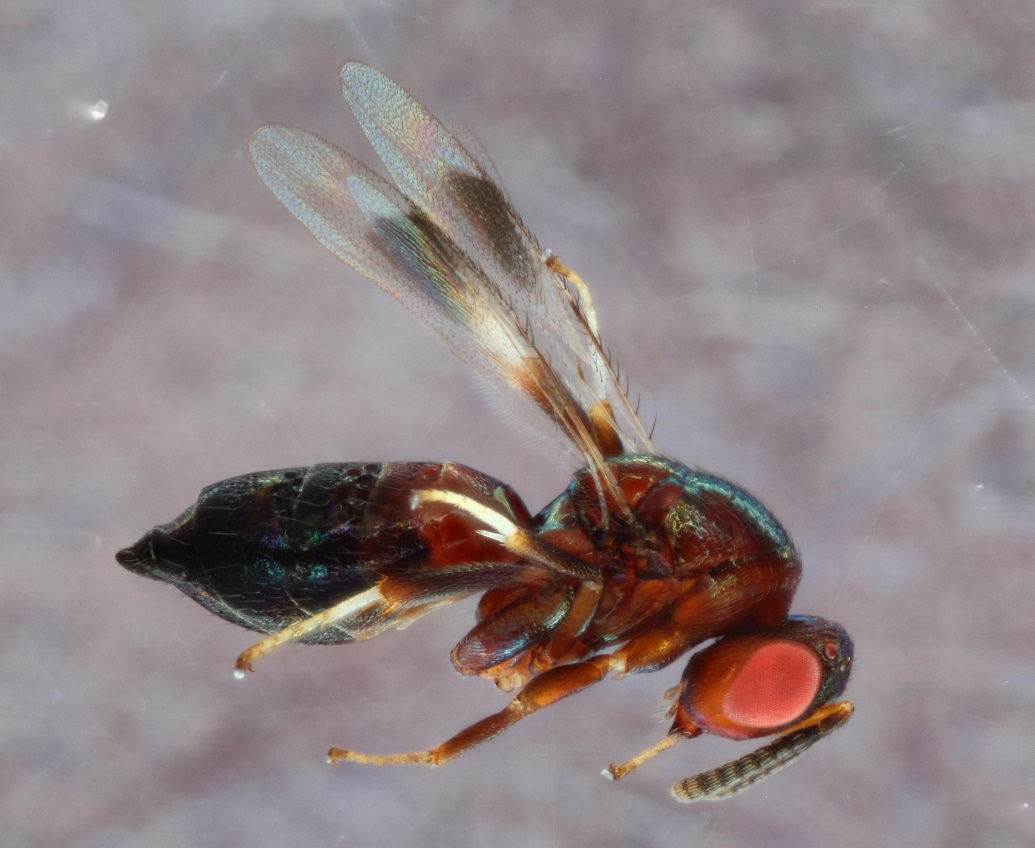

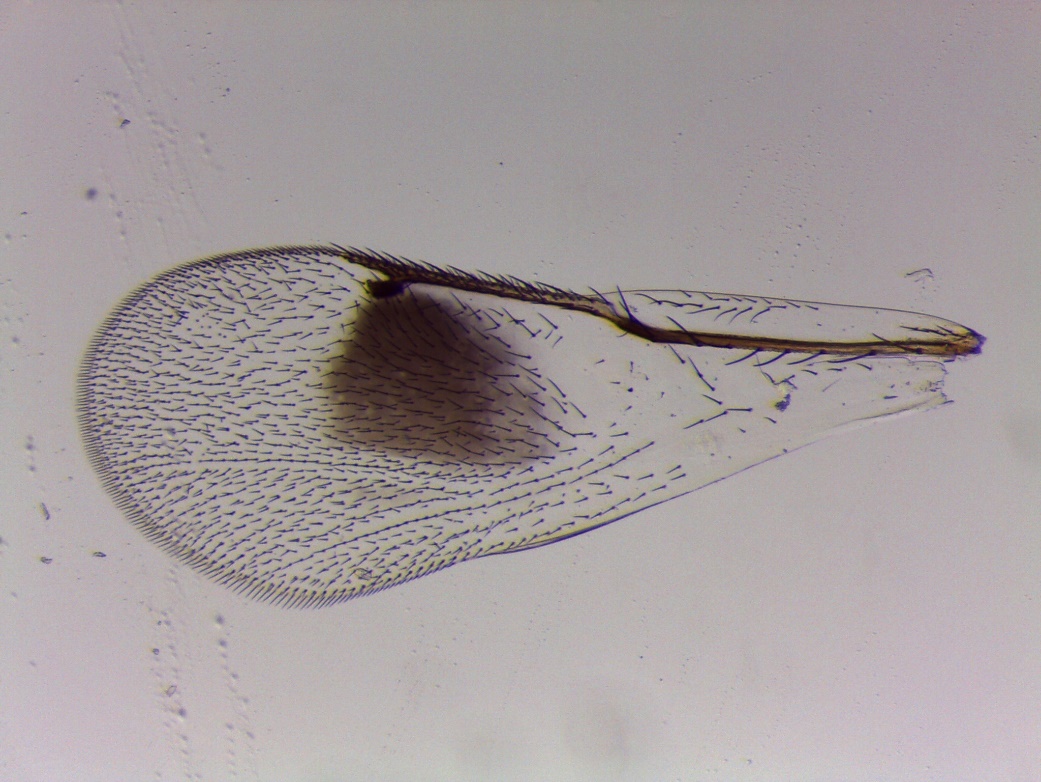

**Figure S3. *Ormyrus venustus* (clade 3):** 1703-227-1, male from *Disholcaspis pedunculeides* on *Quercus turbinella* – Rio Verde, AZ.

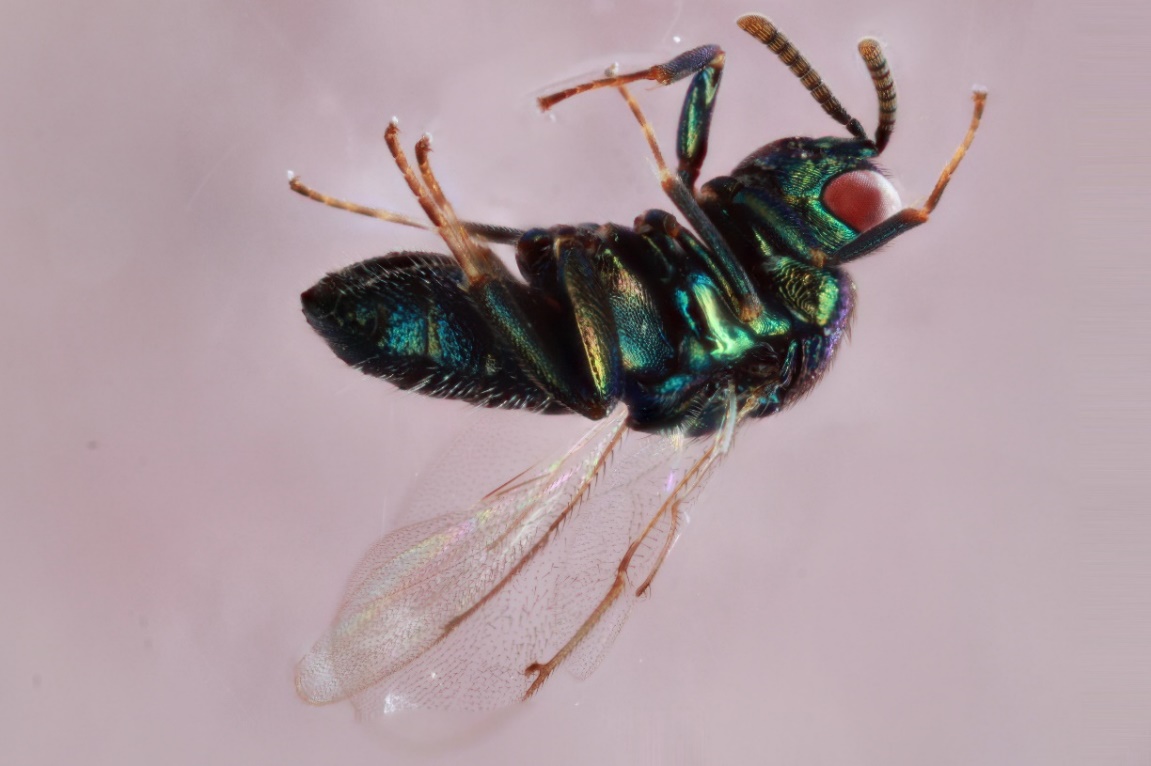

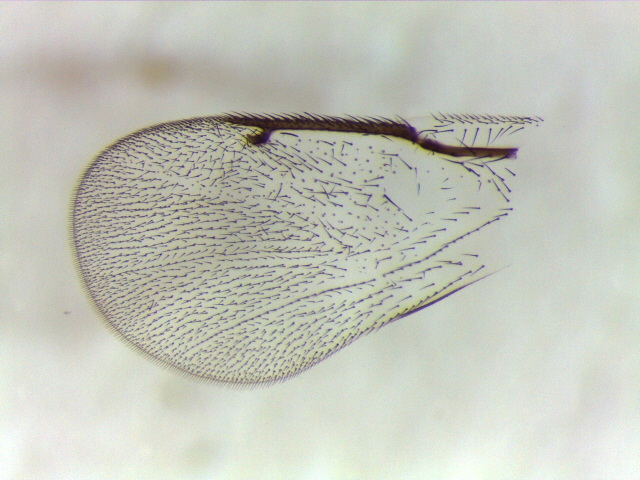

**Figure S4.** ***Ormyrus venustus* (clade 4):** 476-002-1A, female from *Acrapis erinacei* on *Quercus alba* – Iowa City, IA.

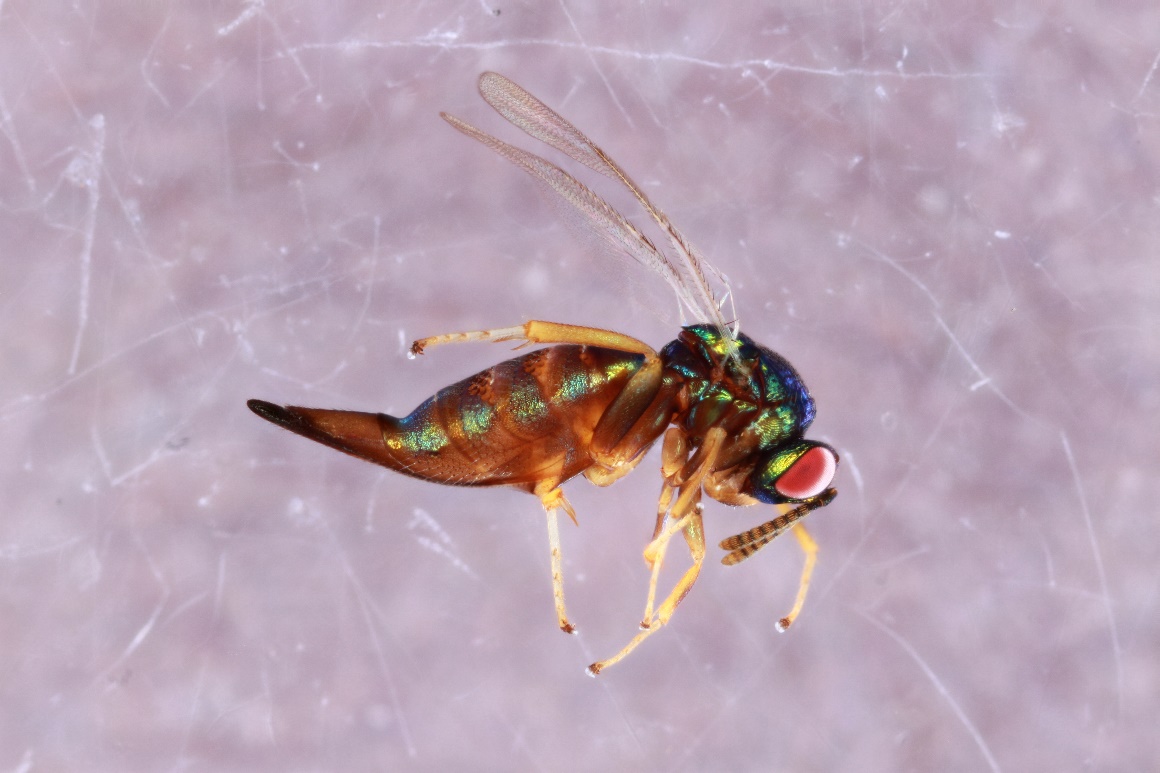

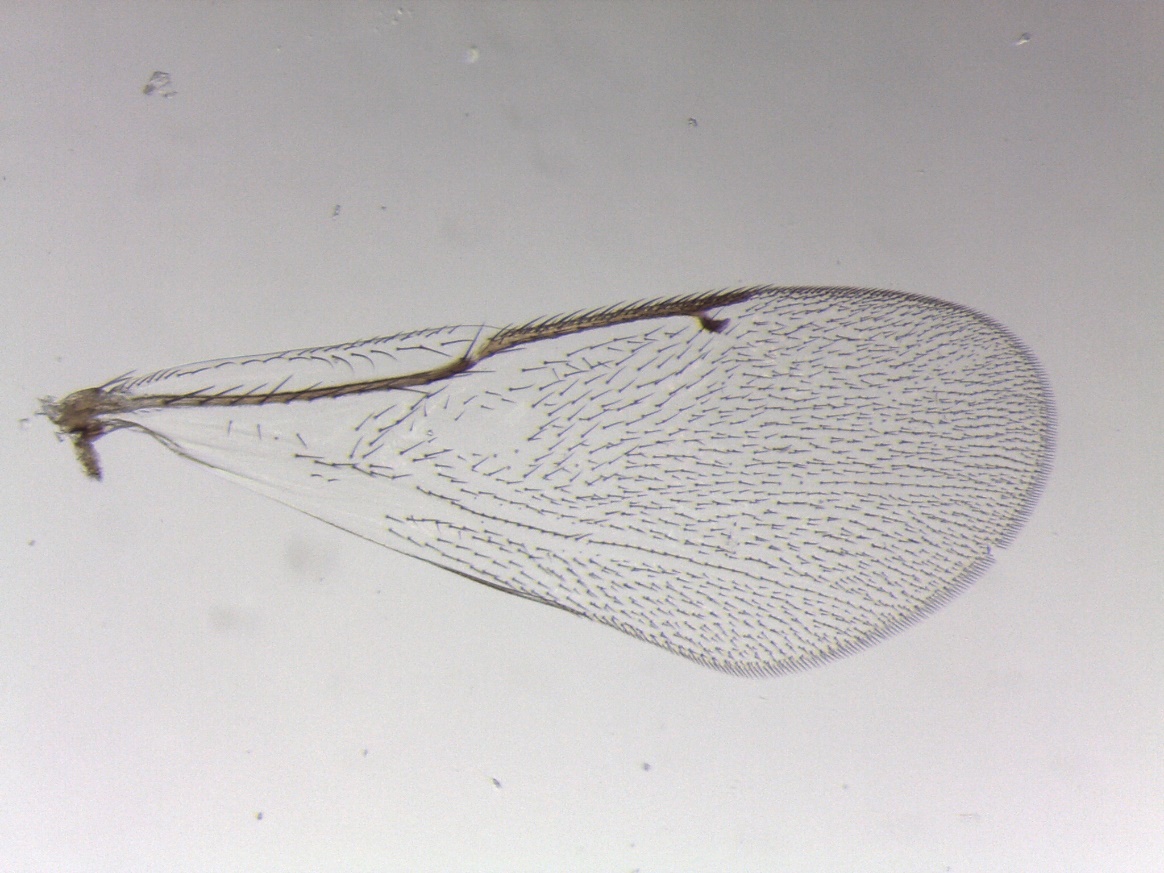

**Figure S5. *Ormyrus* *venustus* (clade 5):** 1238-117-6, female from *Andricus quercuspetiolicola* on *Quercus stellata* – Austin, TX.

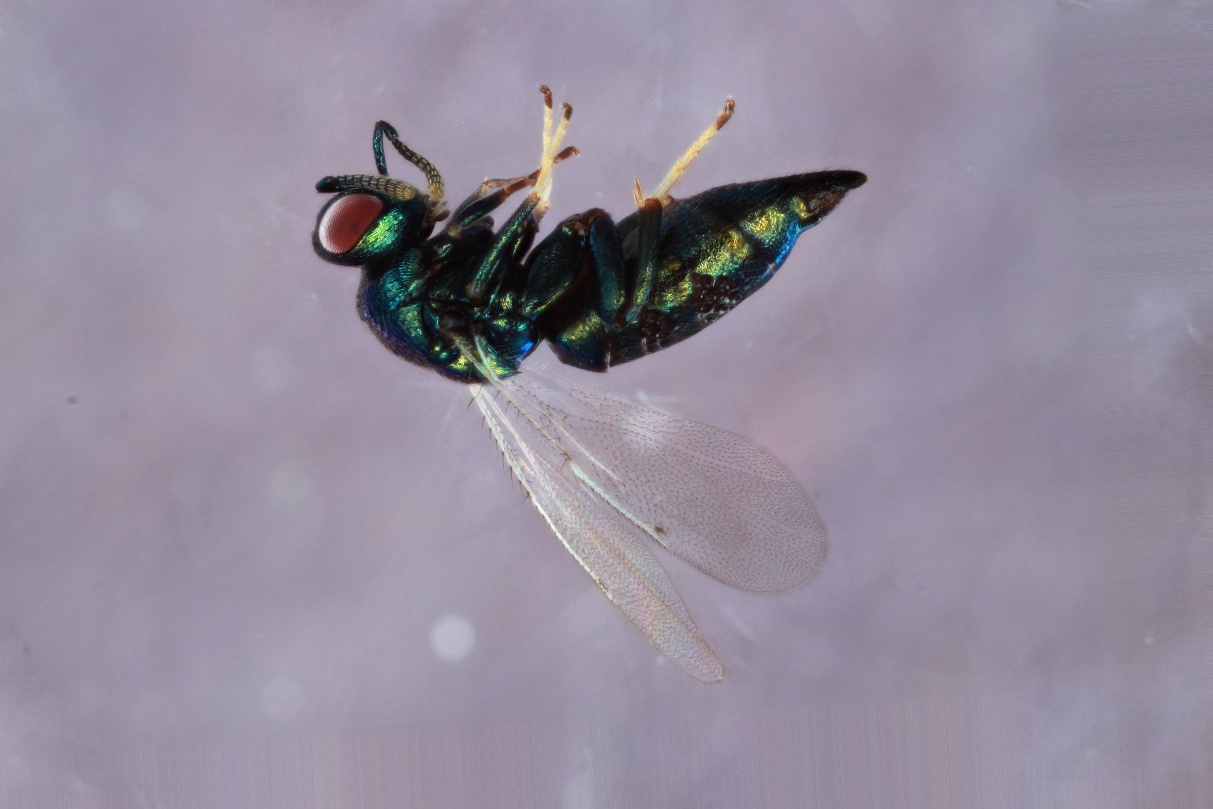

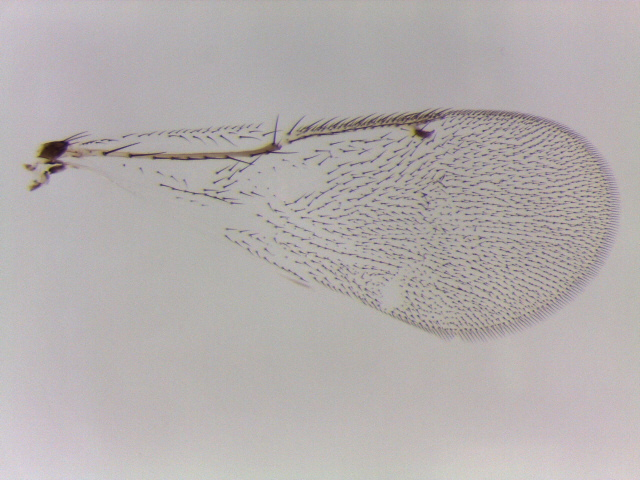

**Figure S6. *Ormyrus* unidentified (clade 6):** 1695-223-1, female from an *Atrusca* sp. (radiating fibers gall) on *Quercus turbinella* – Payson, AZ.

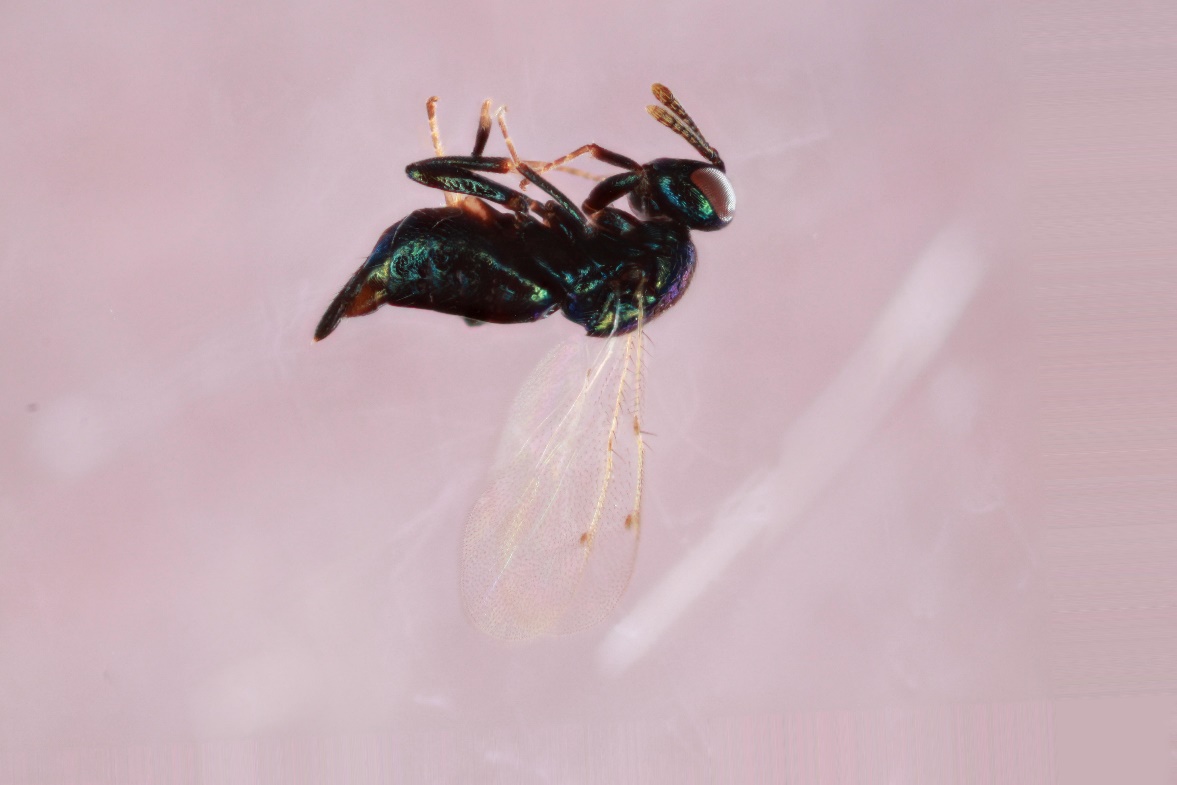

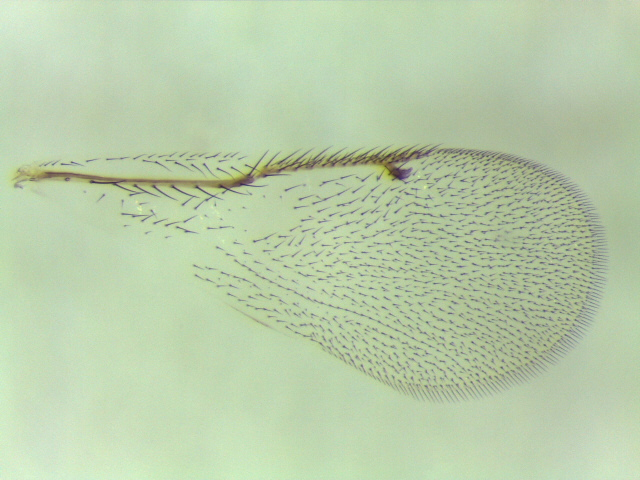

**Figure S7. *Ormyrus* *nr venustus* (clade 7):** 1664-199-4B, female from *Xanothoteras eburneum* on *Quercus gambelii* – Show Low, AZ.

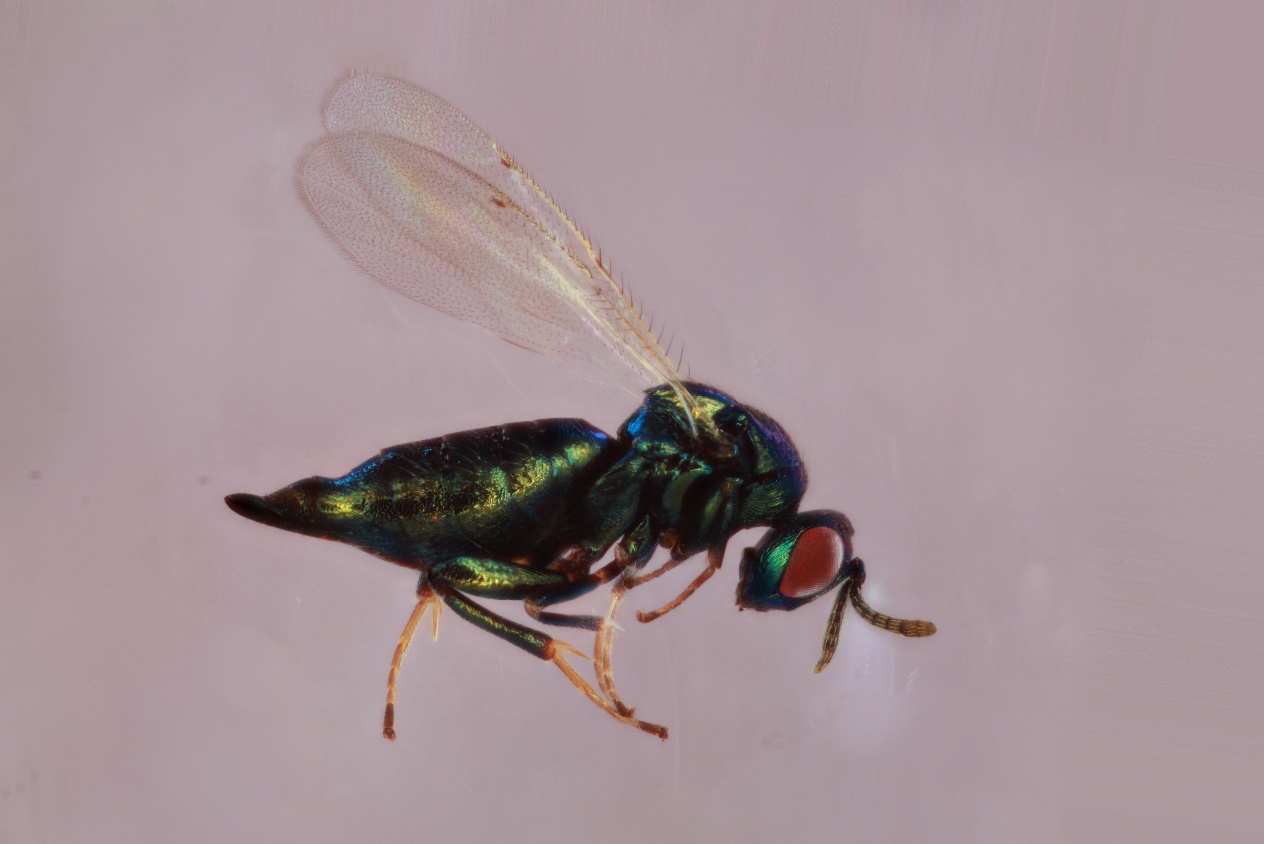

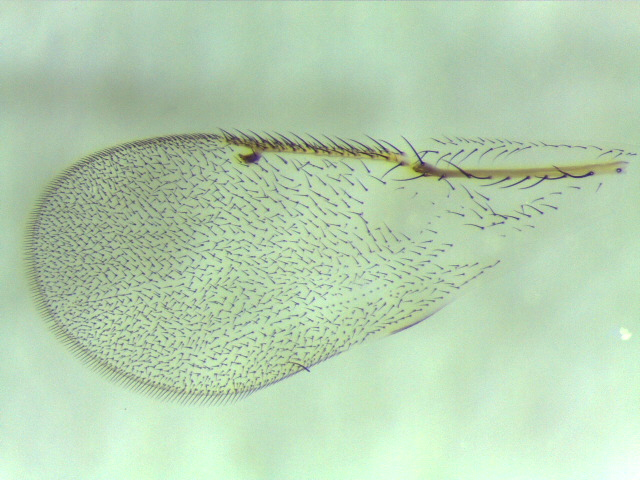

**Figure S8. *Ormyrus* *nr venustus* (clade 7):** 1664-199-4A, male from *Xanothoteras eburneum* on *Quercus gambelii* – Show Low, AZ.
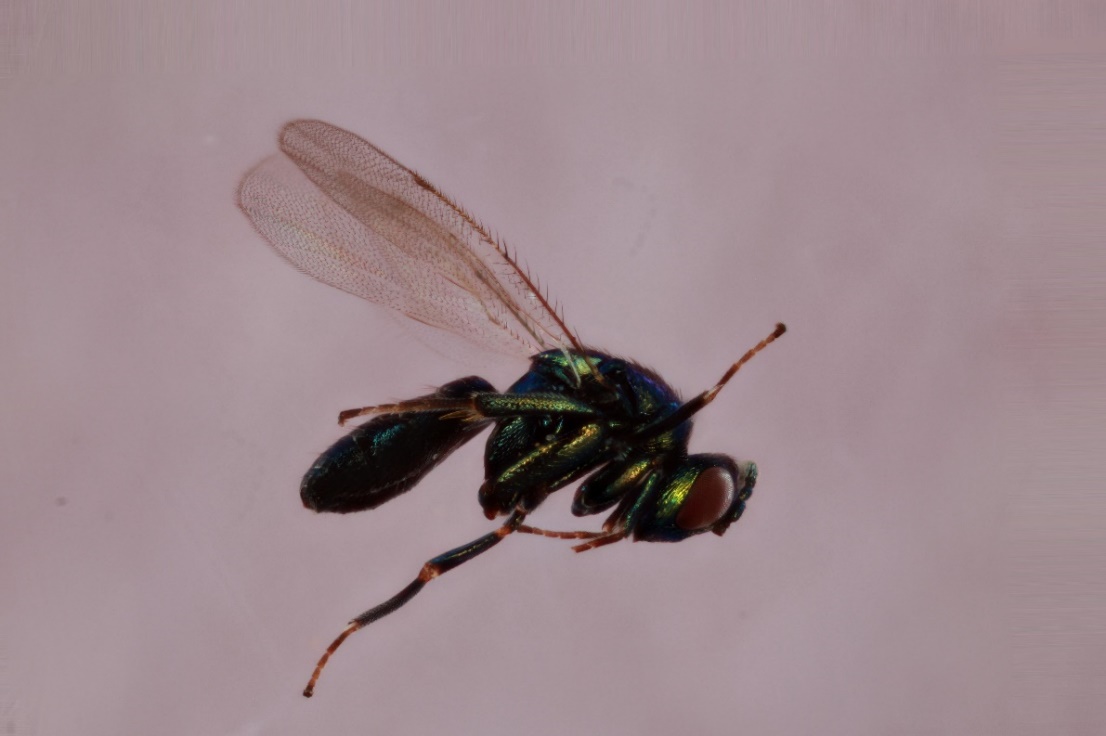

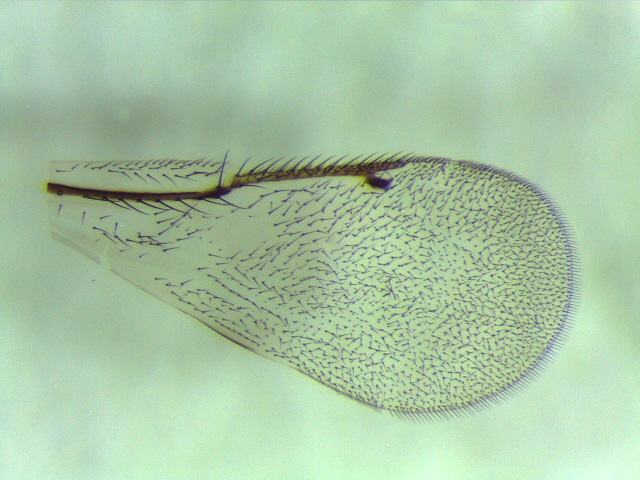

**Figure S9. *Ormyrus reticulatus* (clade 8):** 701-019-026 (pinned), representative female of this species from *Disholcaspis quercusglobulus* on *Quercus alba* – Iowa City, IA. This specimen’s COI was not sequenced, however an *Ormyrus* from the same collection (701-019-026) is present in the phylogeny.

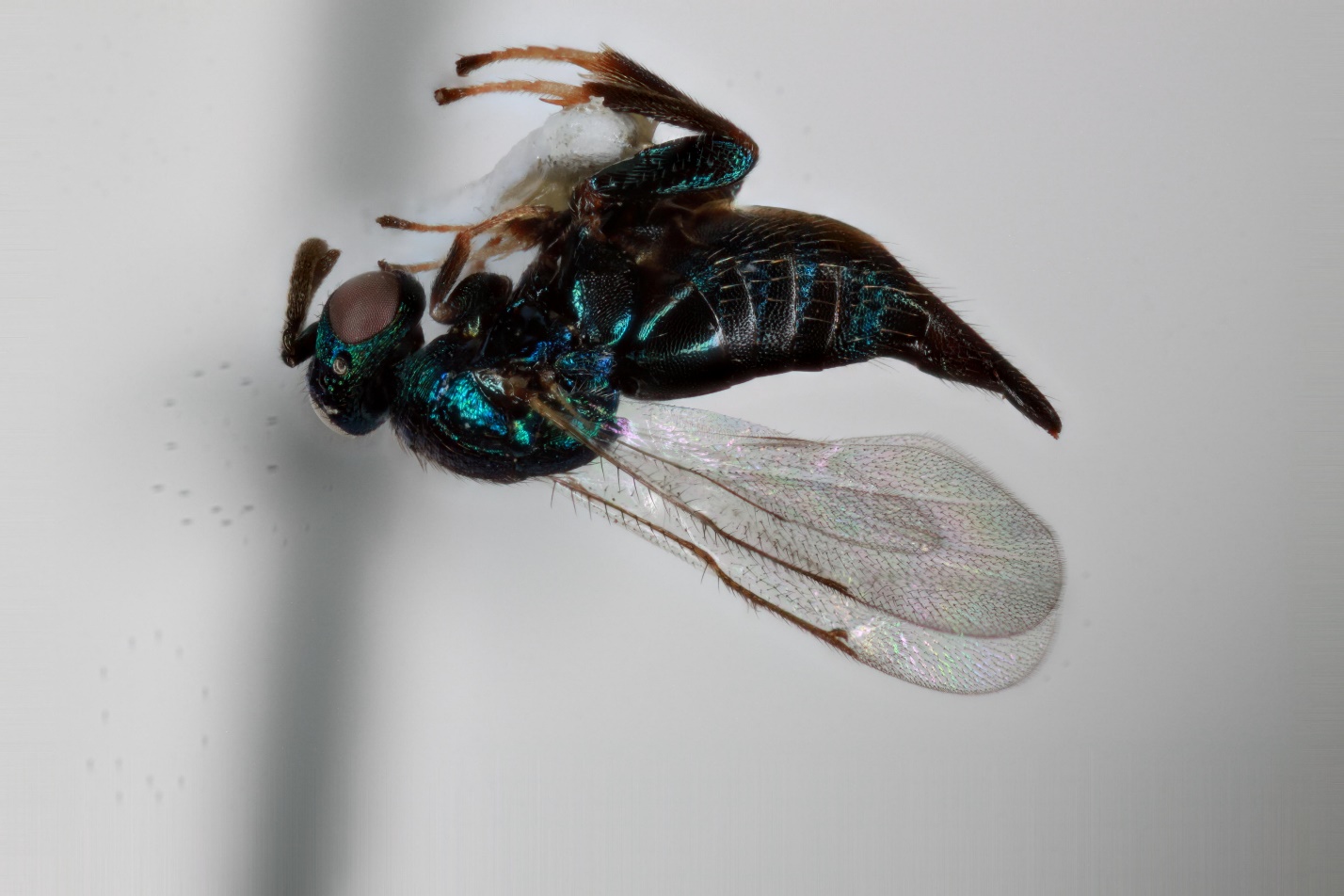

**Figure S10. *Ormyrus* *labotus* (clade 9)** 643-004-5, male from *Acraspis villosa* on *Quercus macrocarpa* – Iowa City, IA.

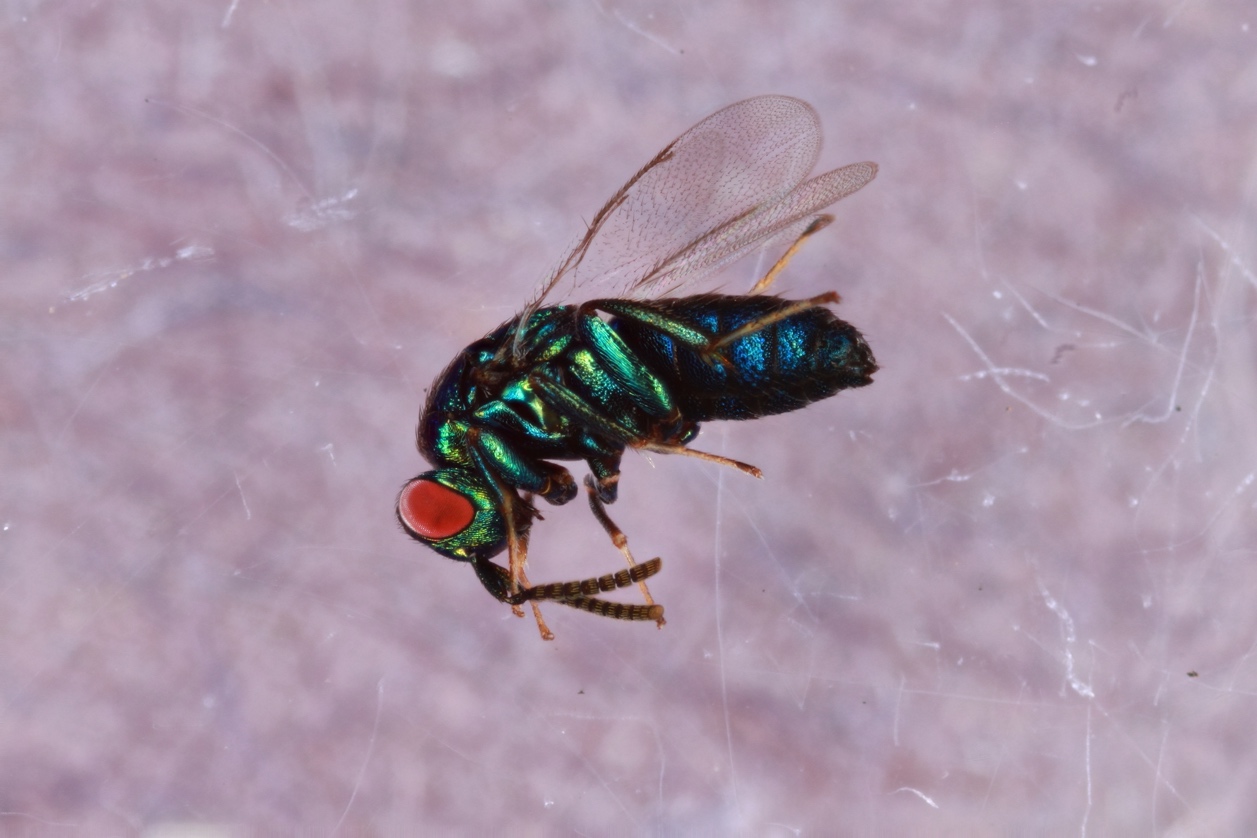

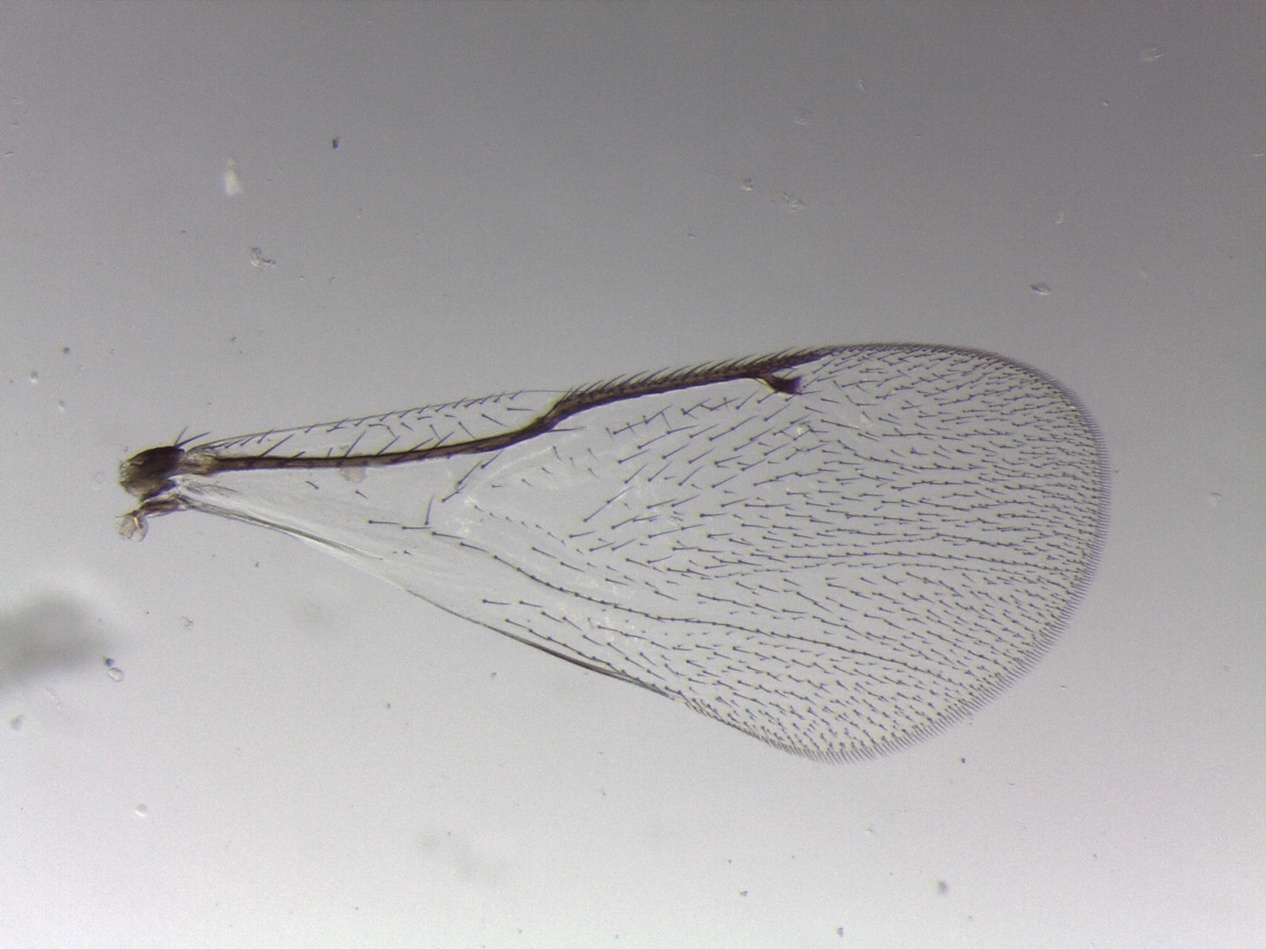

**Figure S11. *Ormyrus* unknown sp. 1 (clade 10)**: YMZ1A, female from *Callirhytis quercusclavigera* on Quercus coccinea – Gainesville, FL

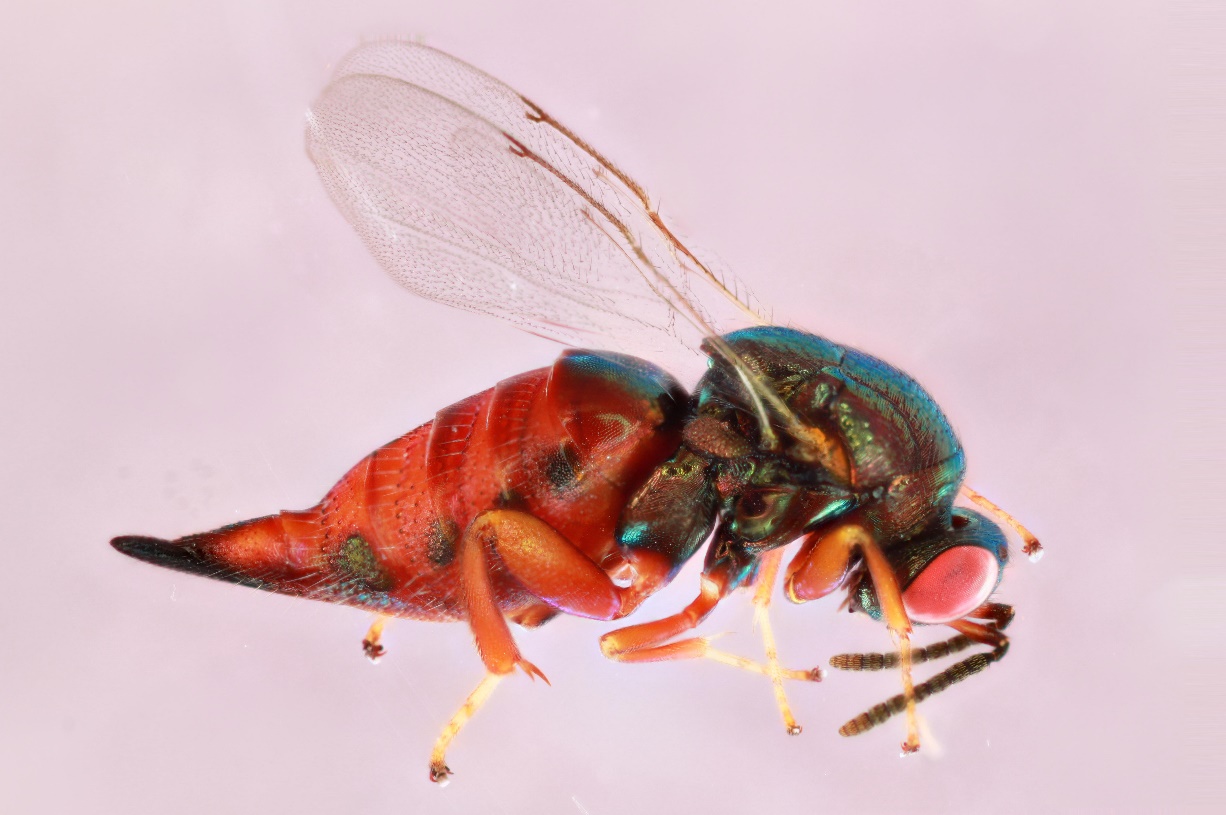

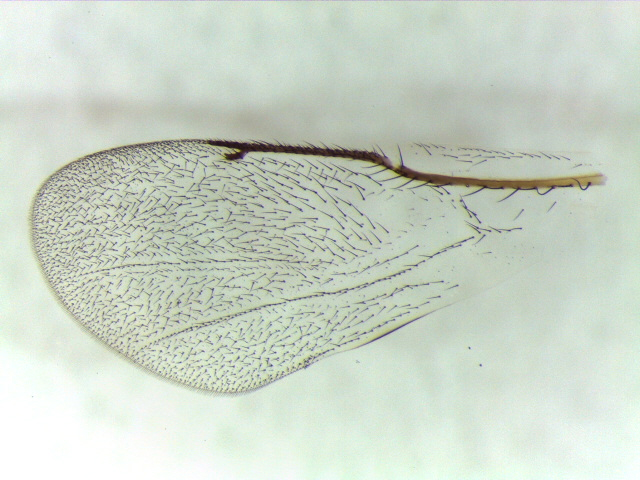

**Figure S12. *Ormyrus* *distinctus* (clade 11):** 1466-126-2, male from *Cynips douglasii* *on Quercus lobata* – Folsom, CA.

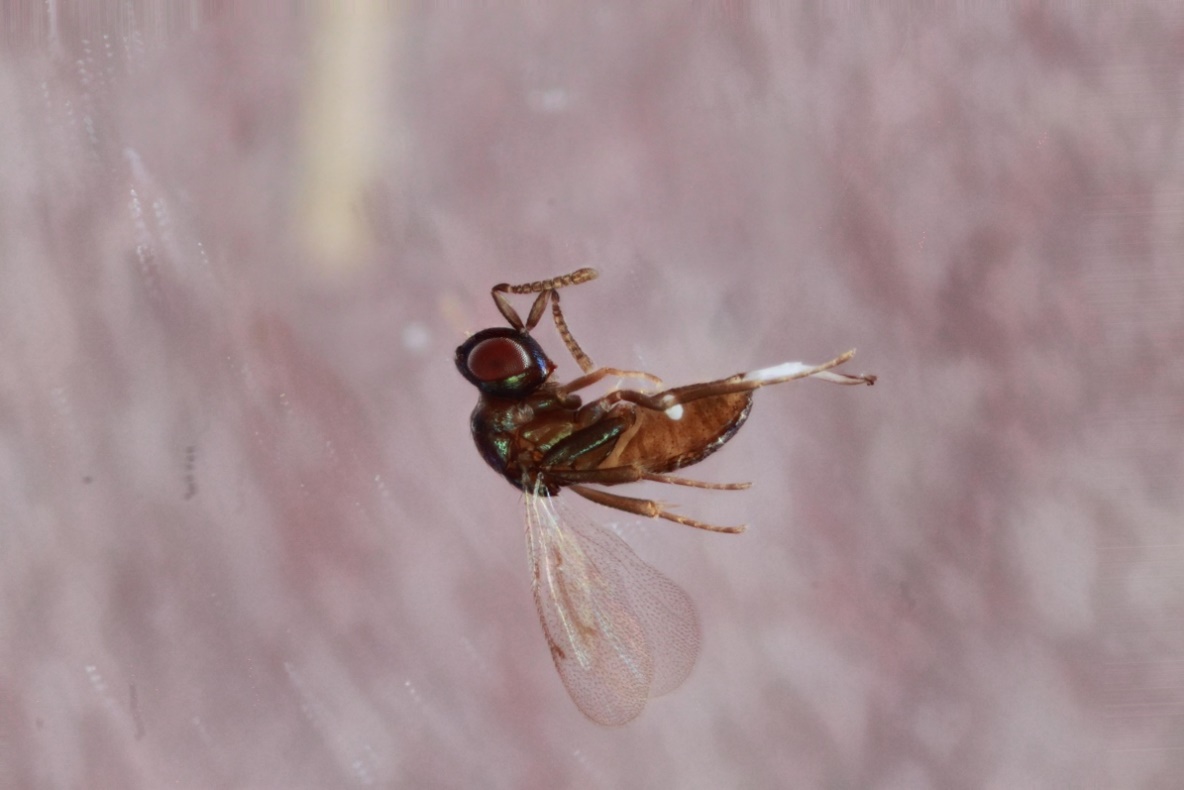

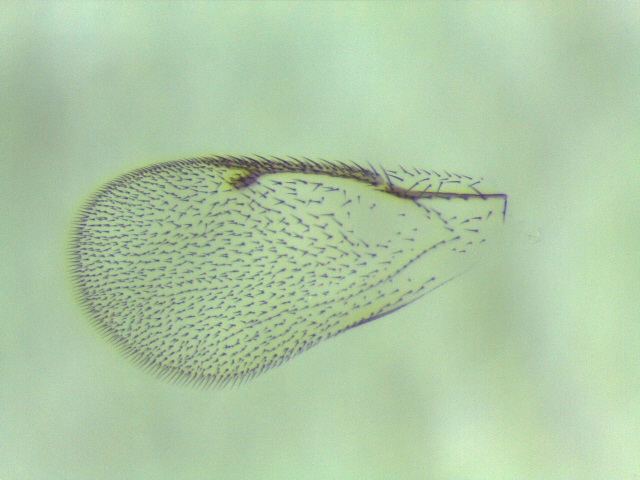

**Figure S13. *Ormyrus* *distinctus* (clade 12):** 1520-150-1A, male unknown spangle gall, similar in appearance to *Phylloteras cupella*, on *Quercus dumosa* – Borrego Springs, CA.

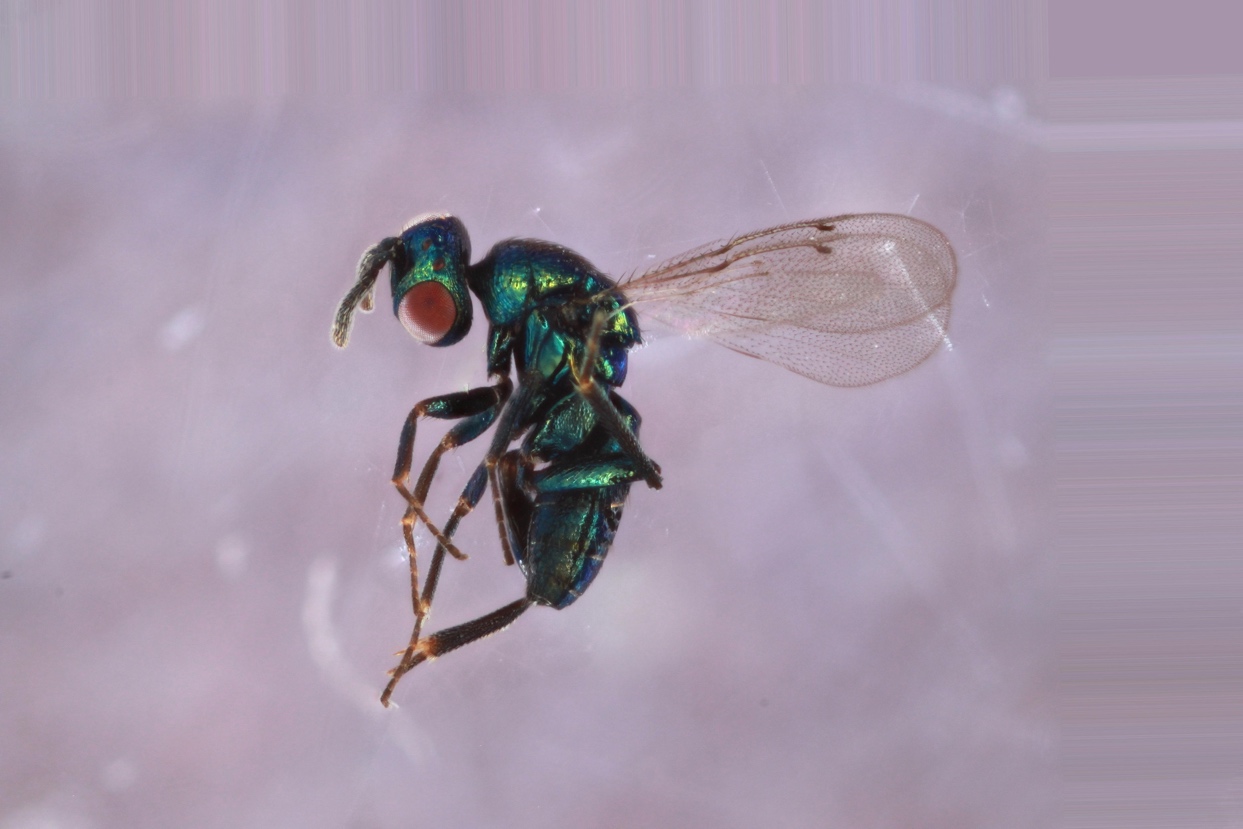

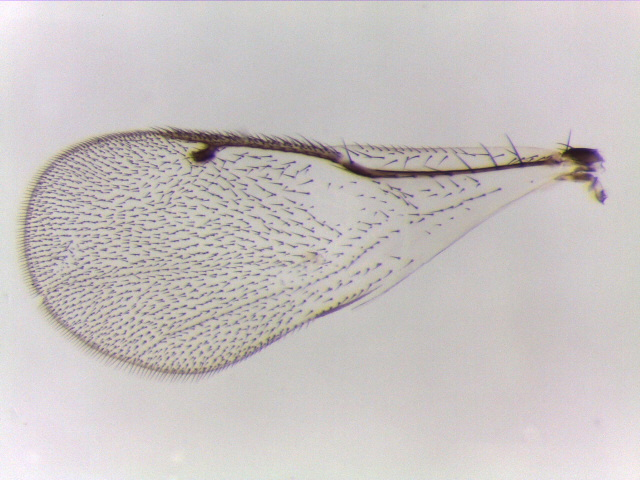

**Figure S14. *Ormyrus* *distinctus* (clade 13):** 1523-145-1, female from unknown stem gall on *Quercus dumosa* – Borrego Springs, CA.

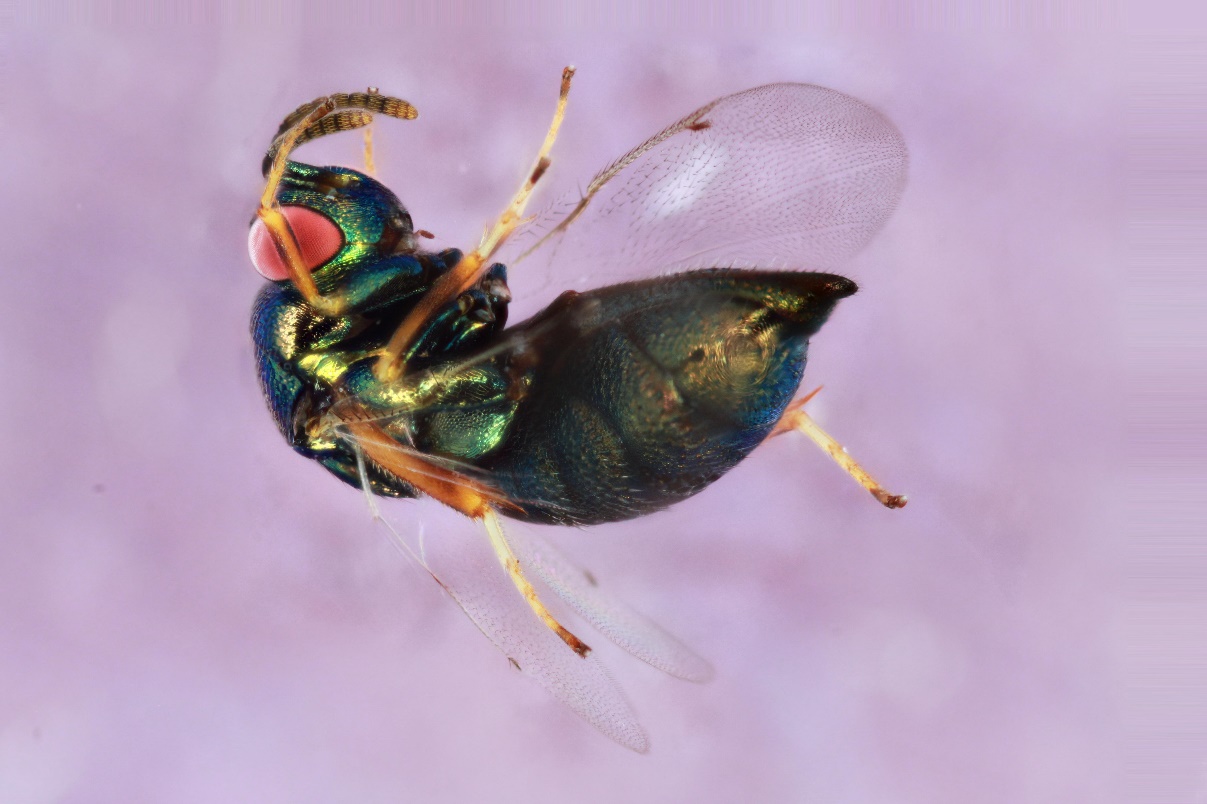

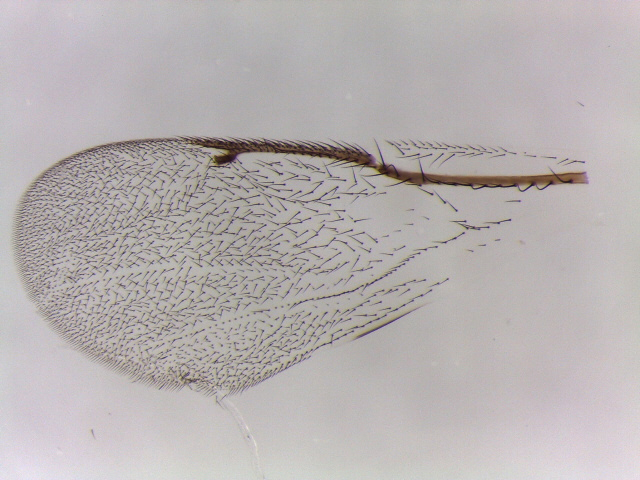

**Figure S15. *Ormyrus* unidentified (clade 14):** YMZ6A, female from *Disholcaspis quercusvirens* on *Quercus virginiana* – Gainesville, FL.

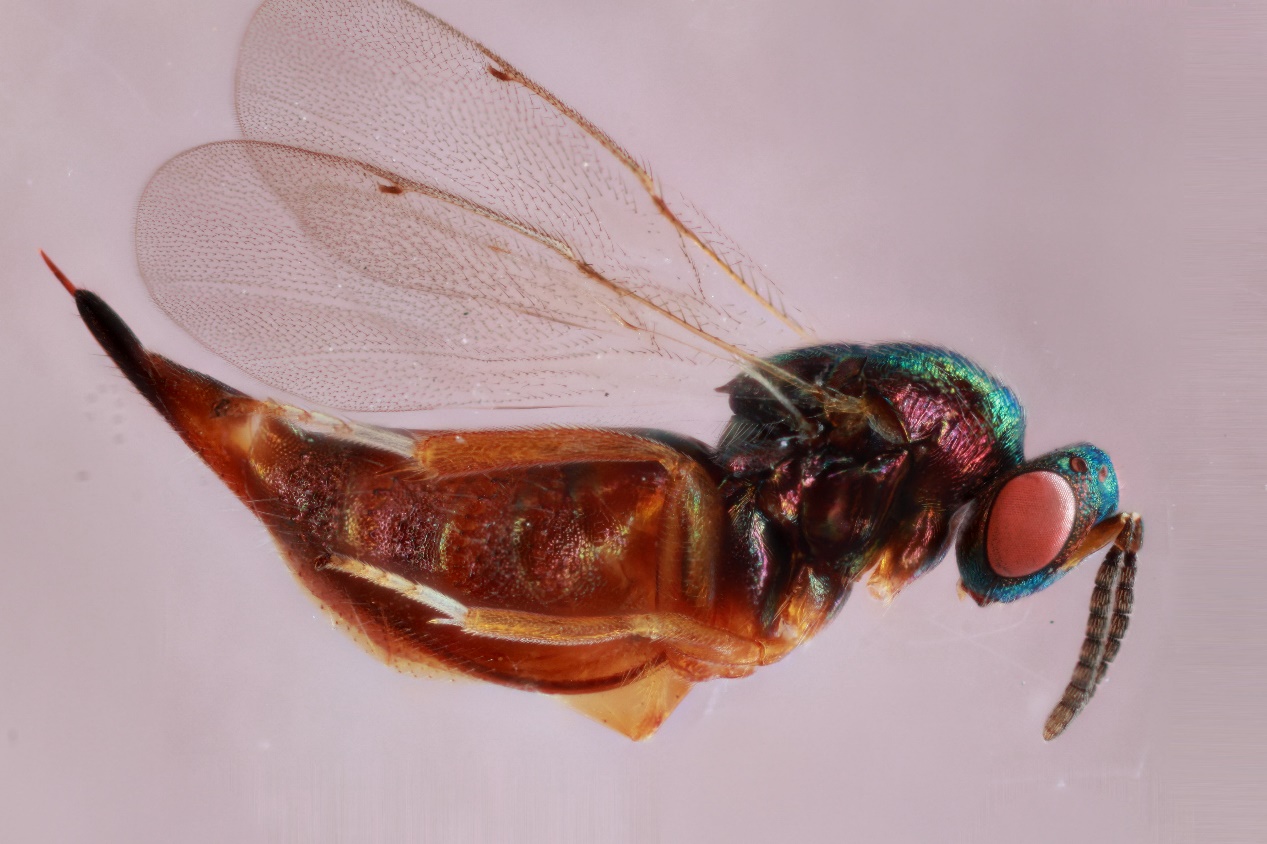

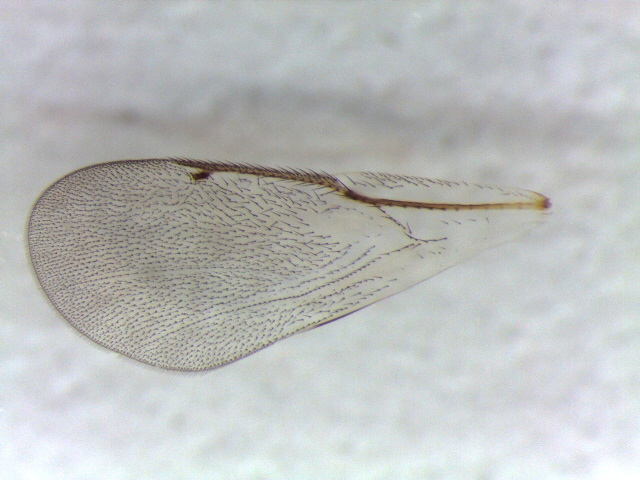

**Figure S16. *Ormyrus* *dryohizoxeni* (clade 15):** Body photo of LZ6608, female from Andricus foliatus on Quercus geminata – Archbold Biological Station, FL. Wing photo of F1001, *Ormyrus* from *Beloncnema treatae* on *Quercus virginiana* – Fort Macon, NC.

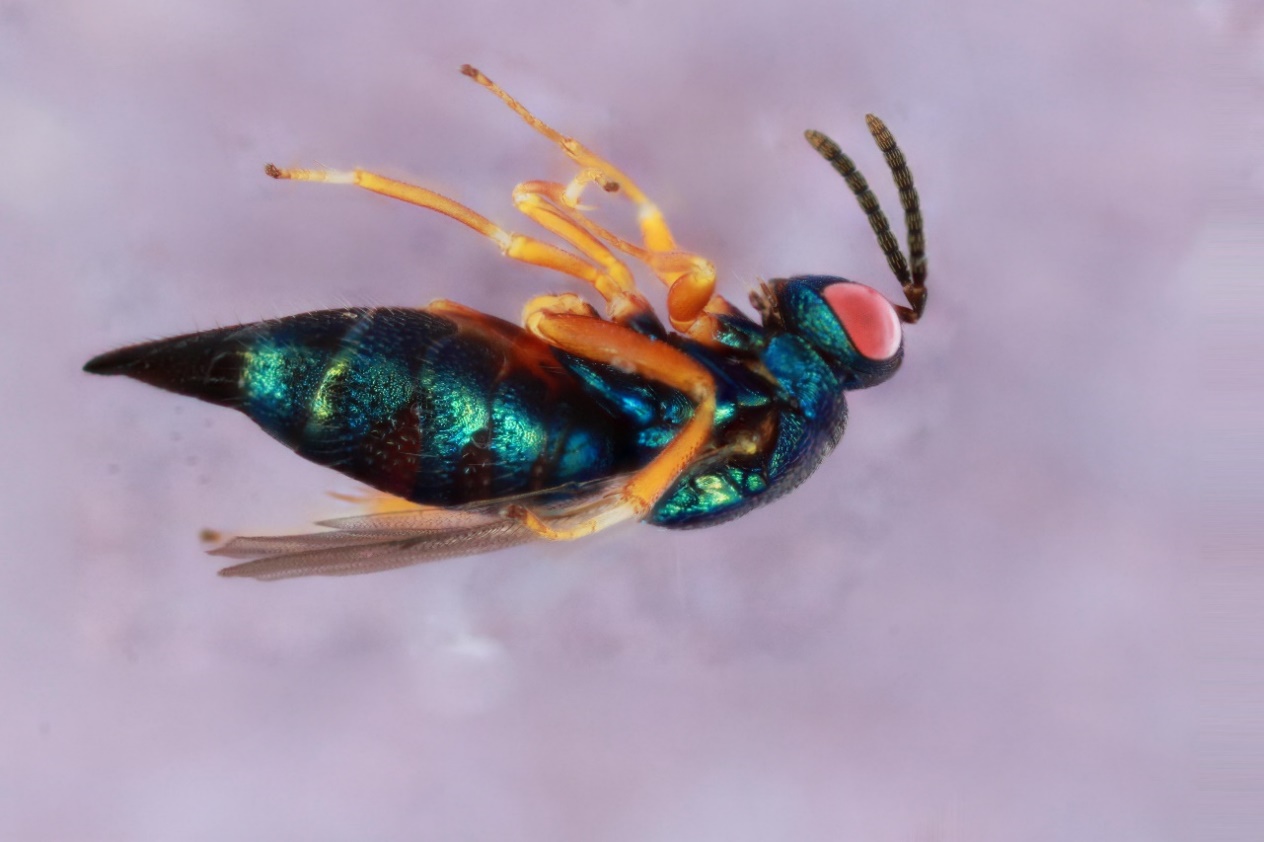

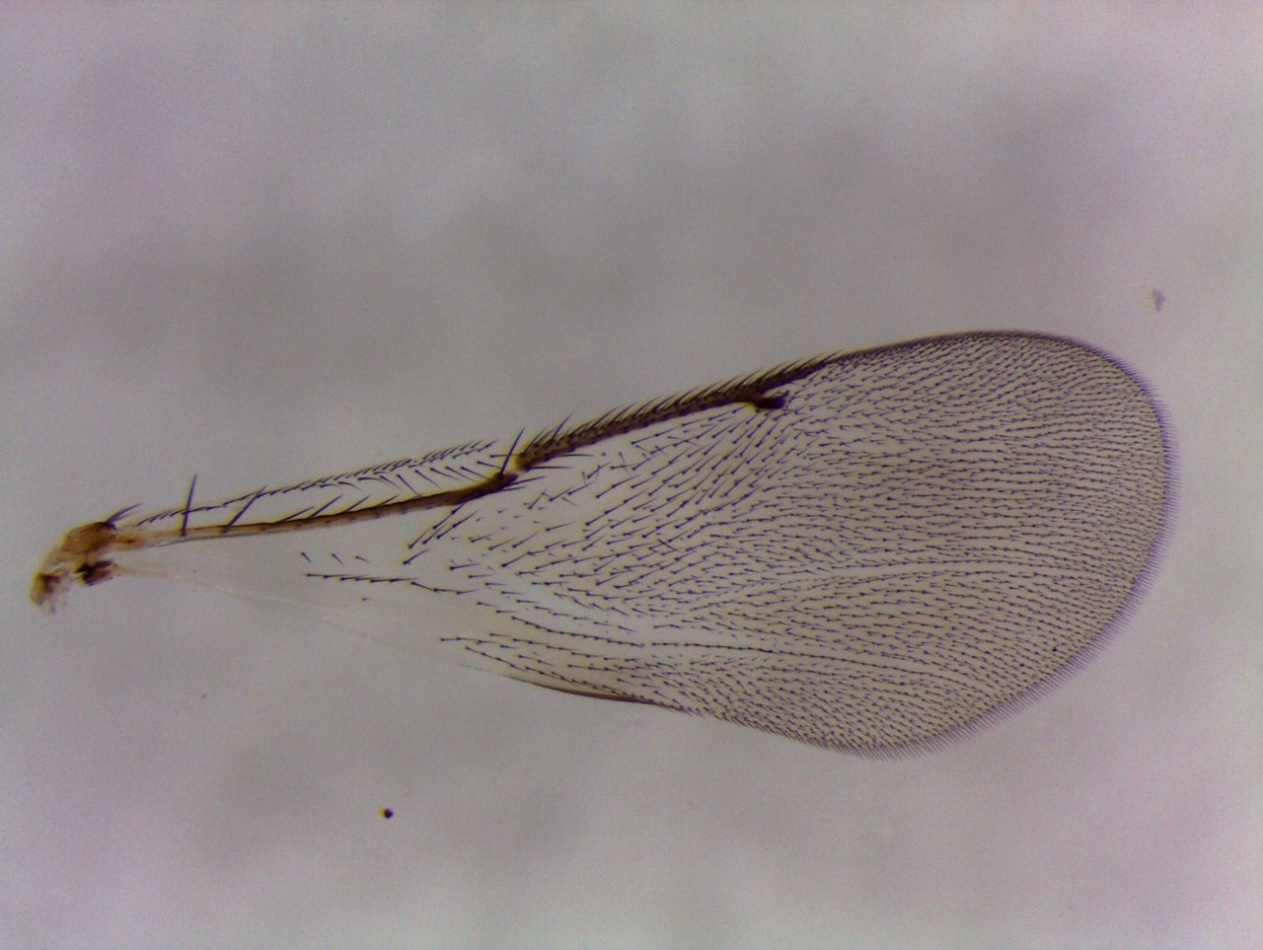

**Figure S17. *Ormyrus* *nr distinctus* (clade 17):** 1693-221-1 from an unidentified acorn gall on *Quercus turbinella* – Payson, AZ.

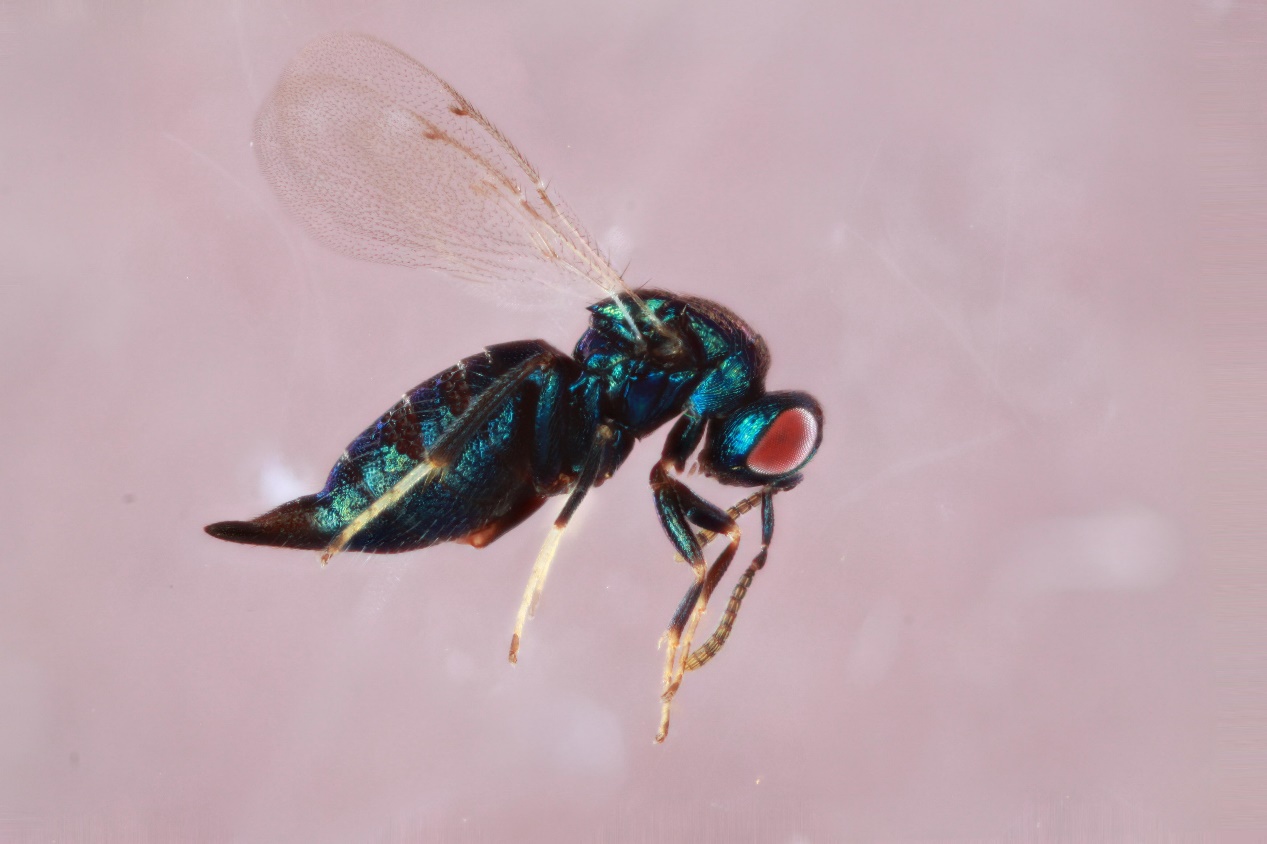

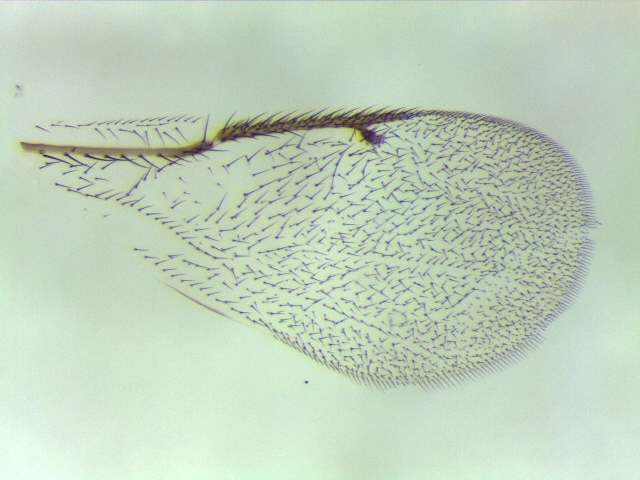

**Figure S18. *Ormyrus* *labotus* (clade 18):** 558-081-1A, female from *Andricus pattoni* on *Quercus stellata* – Peducah, KY.

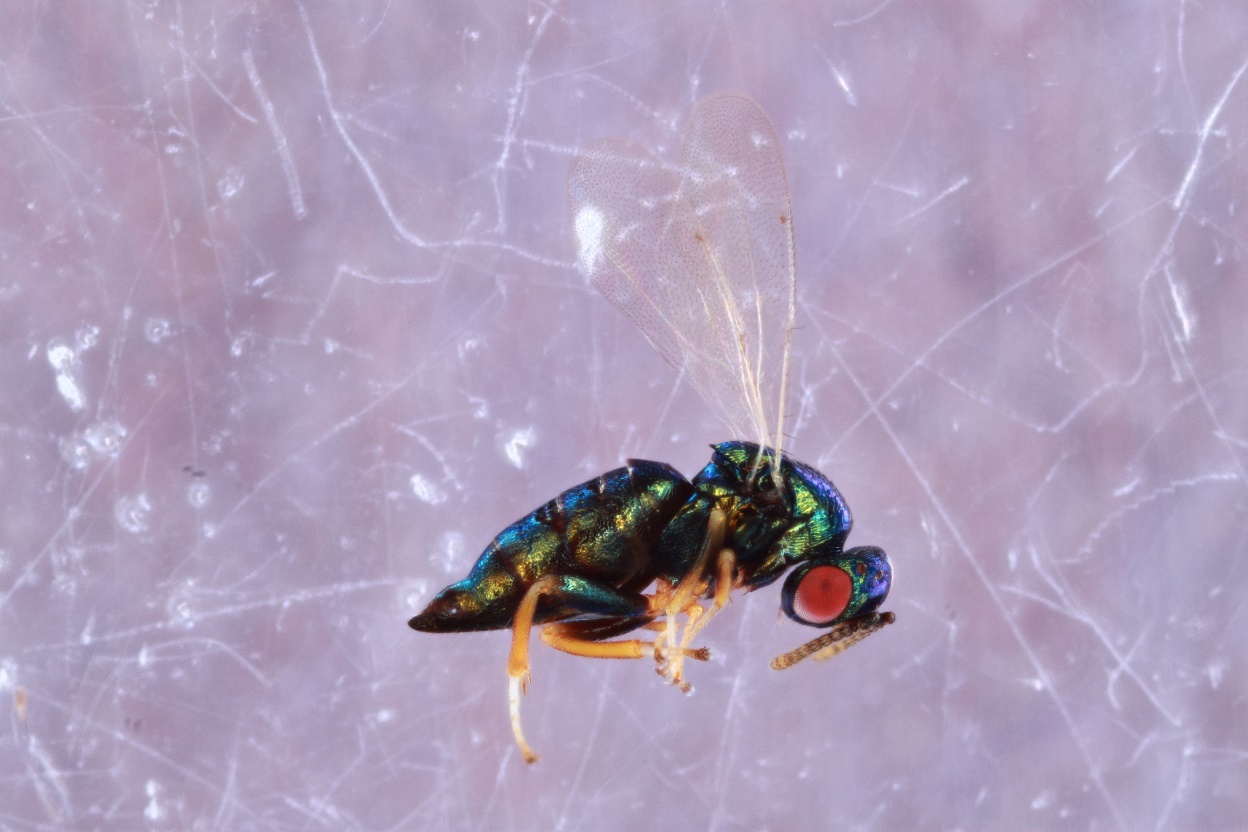

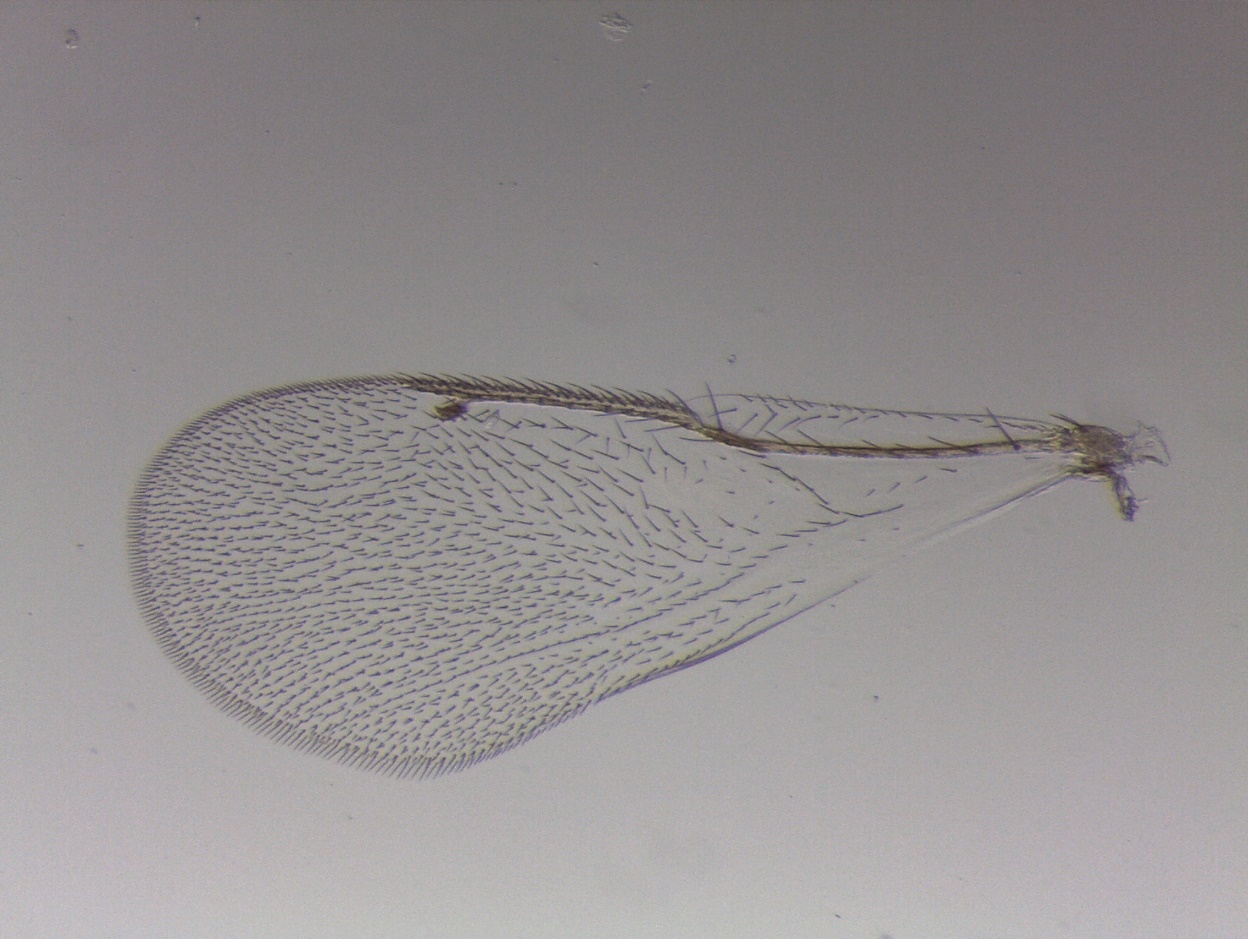

**Figure S19. *Ormyrus* *labotus* (clade 19):** 1553-005-8, female from *Andricus dimorphus* on *Quercus macrocarpa* – Lansing, IA.

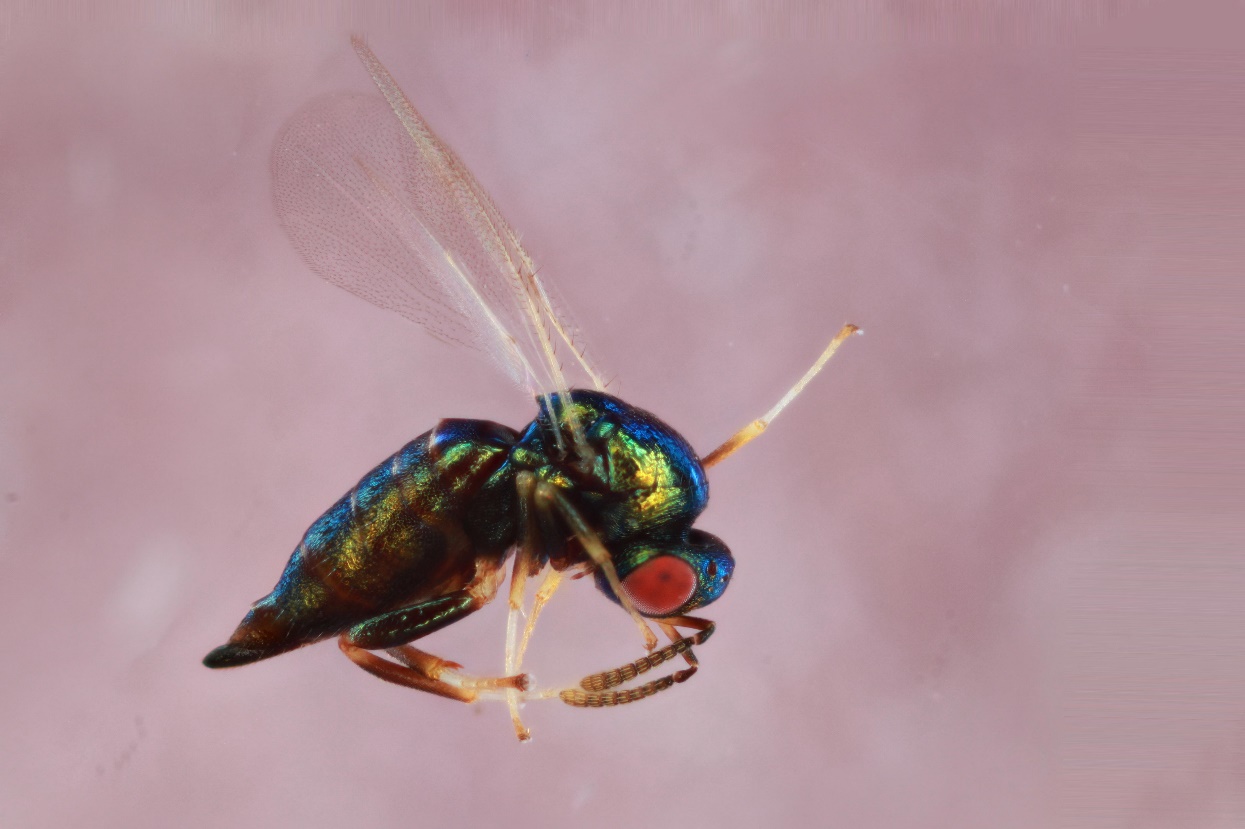

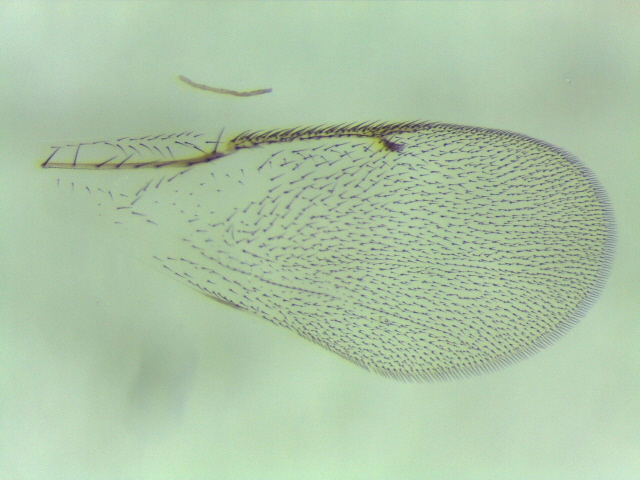

**Figure S20. *Ormyrus* *labotus* (clade 20)**: 1114-104-6, female from *Callirhytis pigra* on *Quercus rubra* – Nashville, TN.

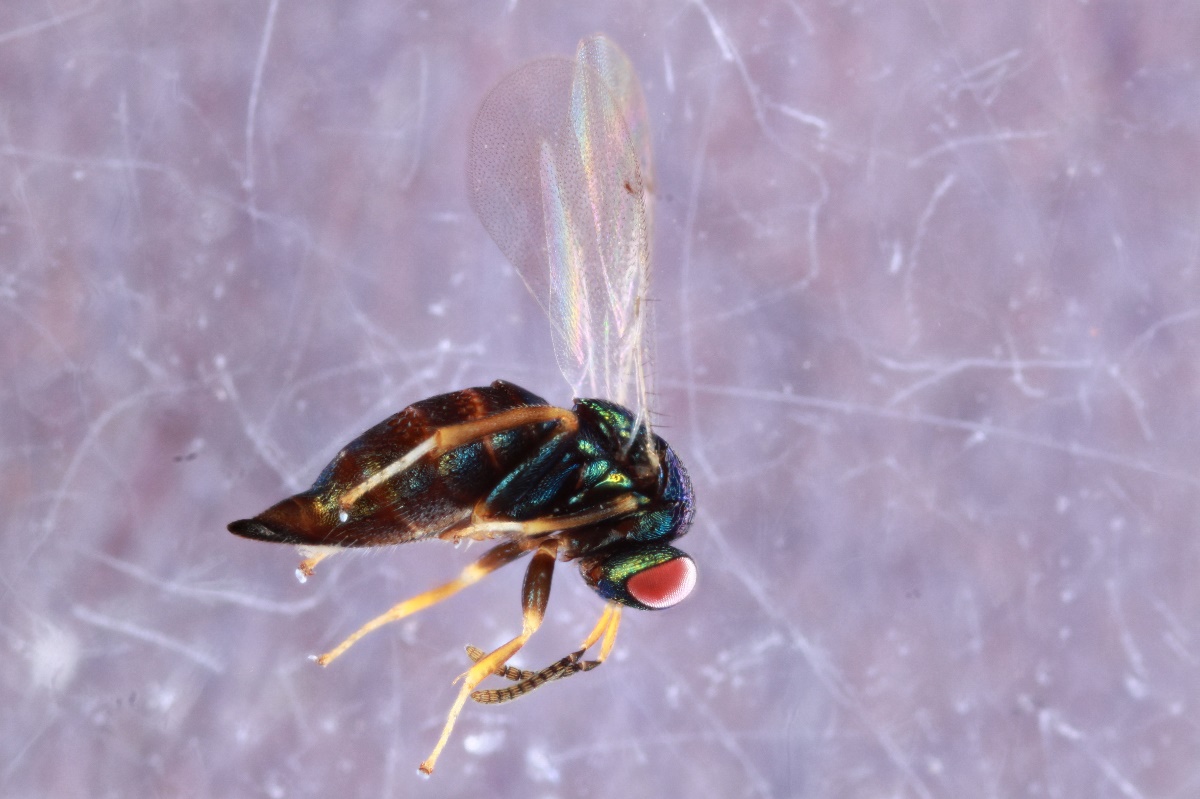

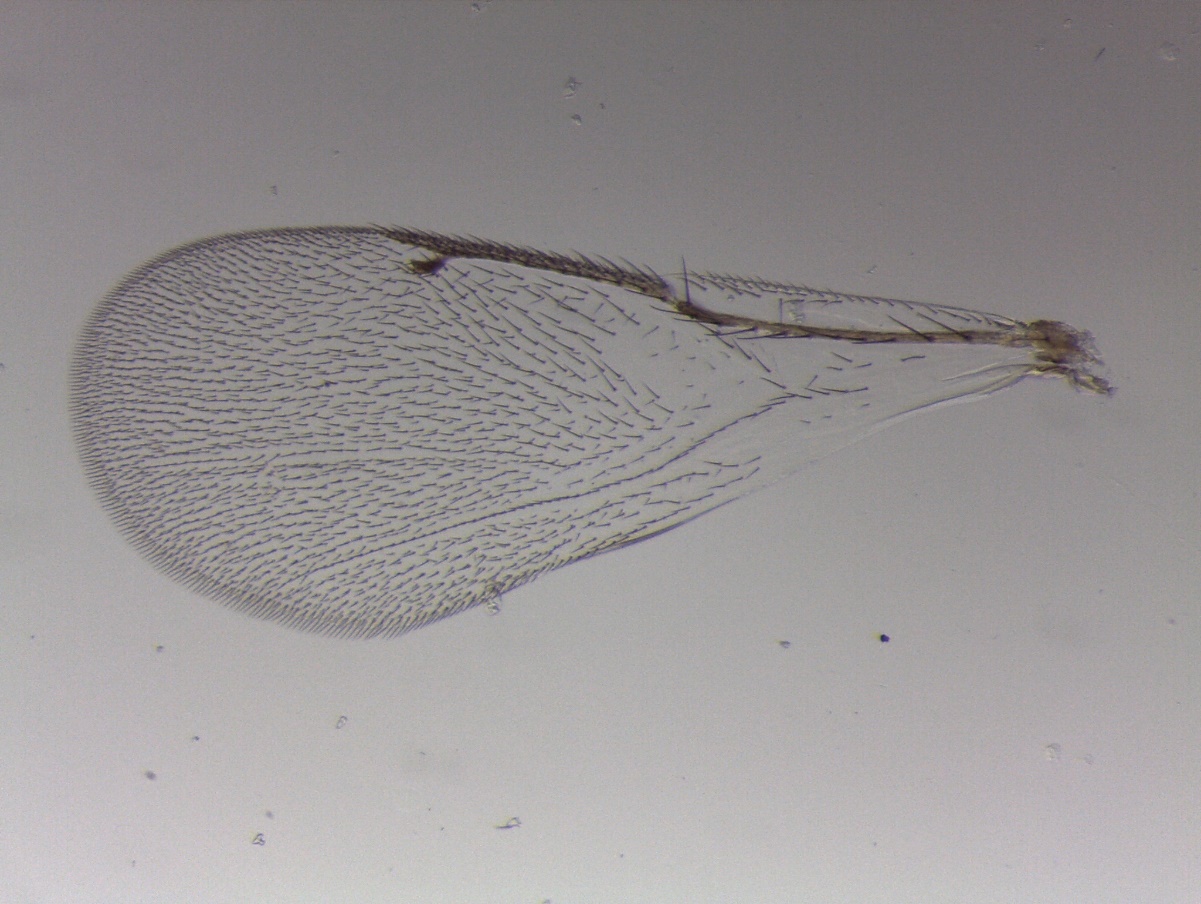

**Figure S21. *Ormyrus* unknown sp. 2 (clade 21):** YMZ5, female unidentified disc-like leaf gall on *Quercus hemisphaerica* – University of Florida, FL.

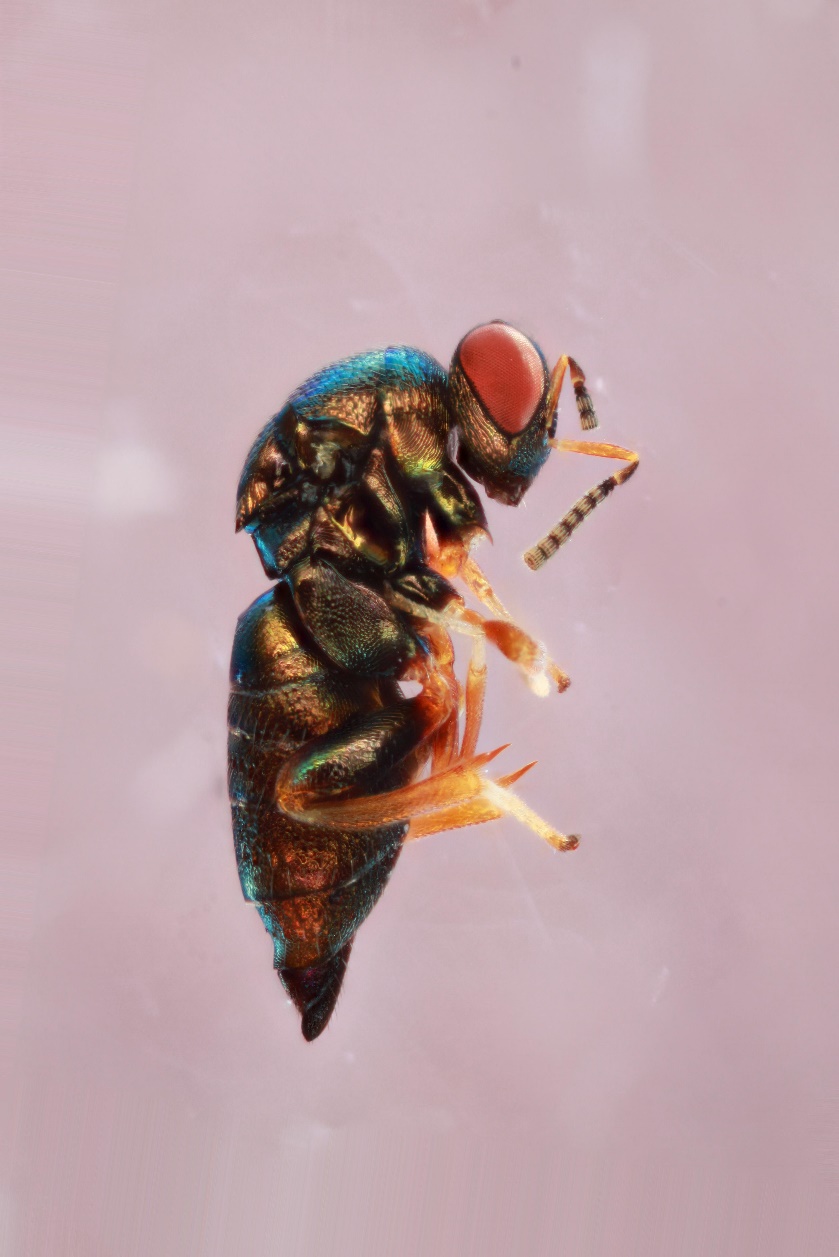

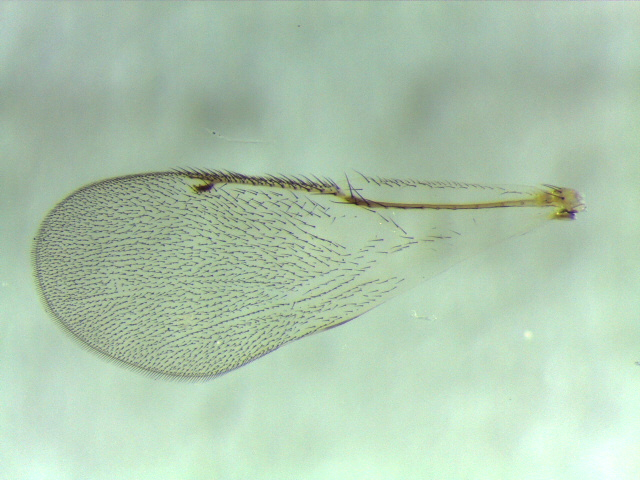

**Figure S22. *Ormyrus* *labotus* (clade 22):** 1574-104-6, female from *Callirhytis pigra* on *Quercus velutina* – Vestal, NY.

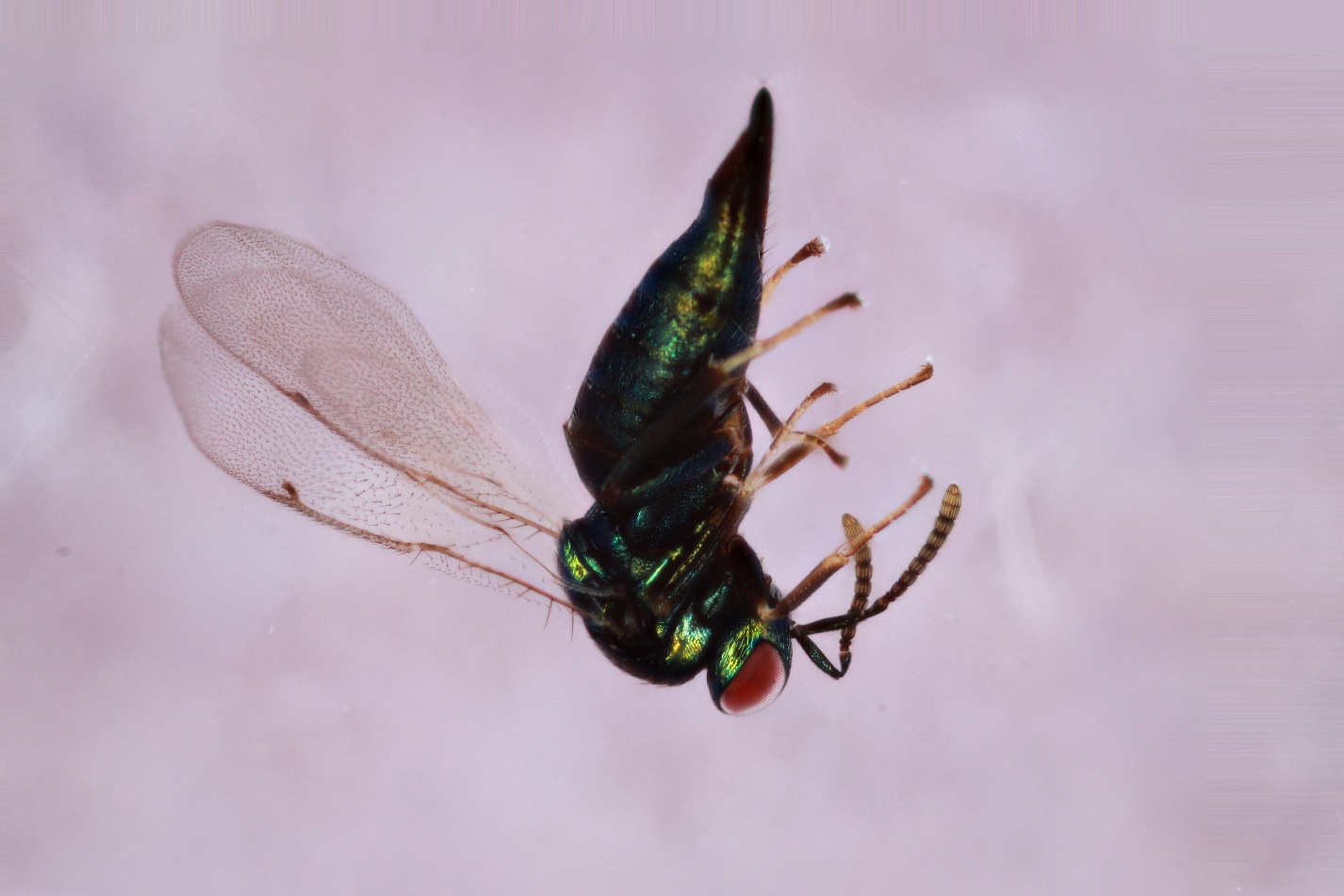

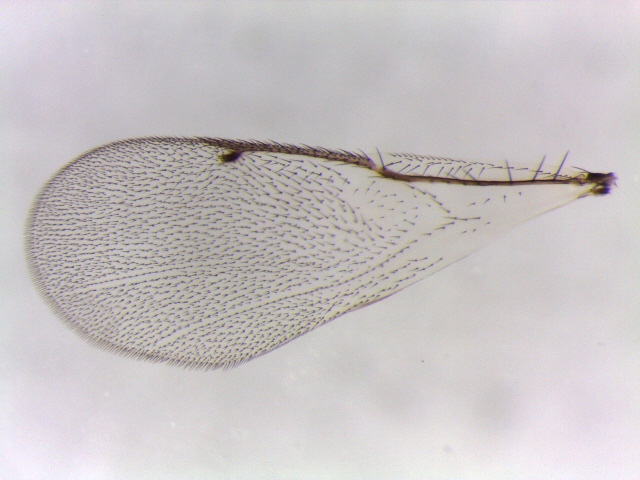

**Figure S23. *Ormyrus* unidentified (clade 23):** 539-067-5A, male from *Andricus robustus* on *Quercus stellata* – St. Louis, MO.

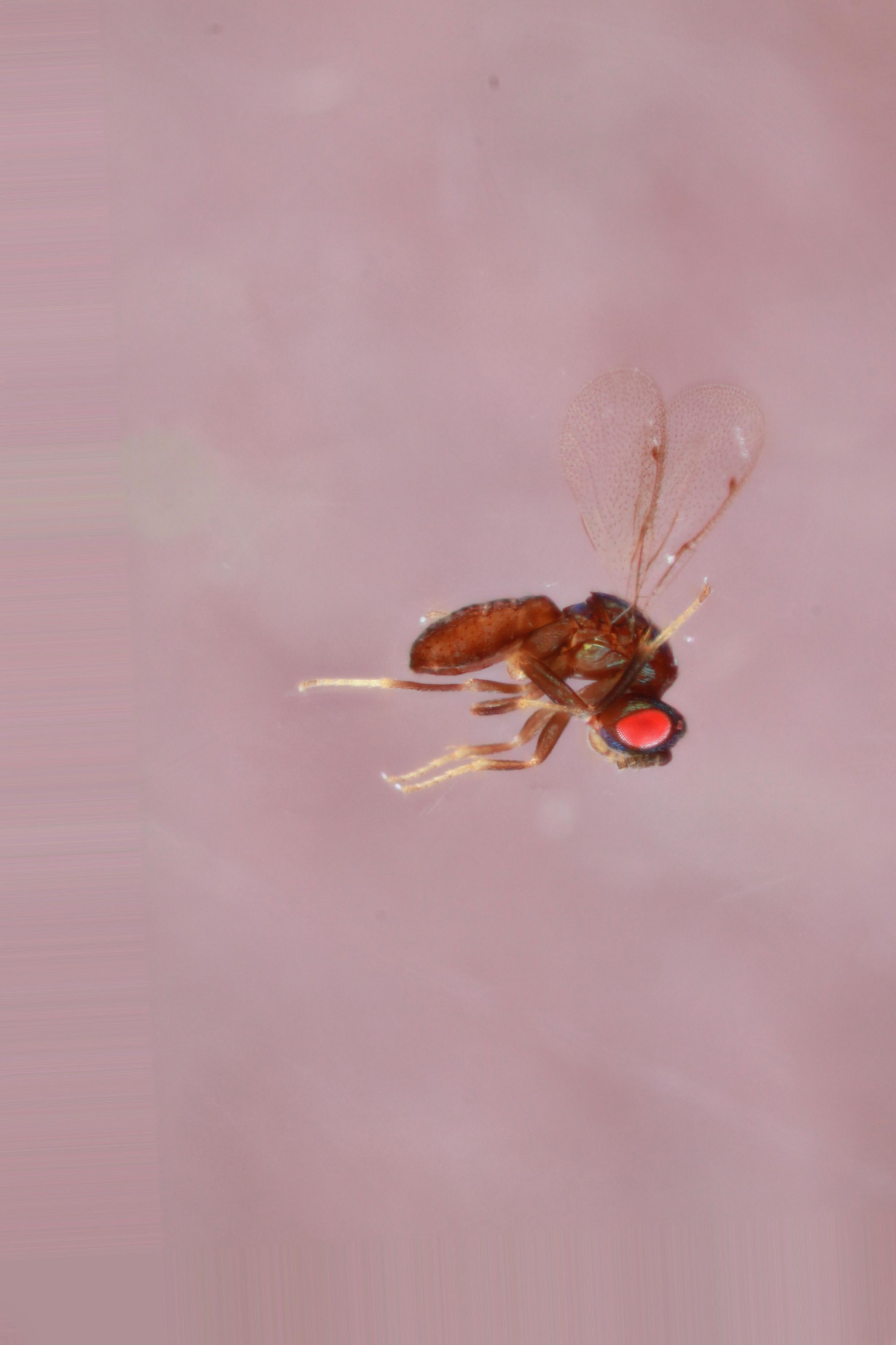

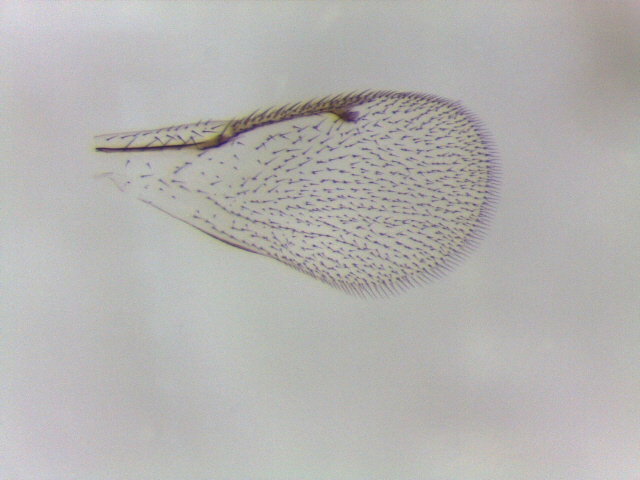

**Figure S24. *Ormyrus* *labotus* (clade 24):** 1335-097-1A, male from unidentified raised vein gall on *Quercus palustris* – City Park, IA.

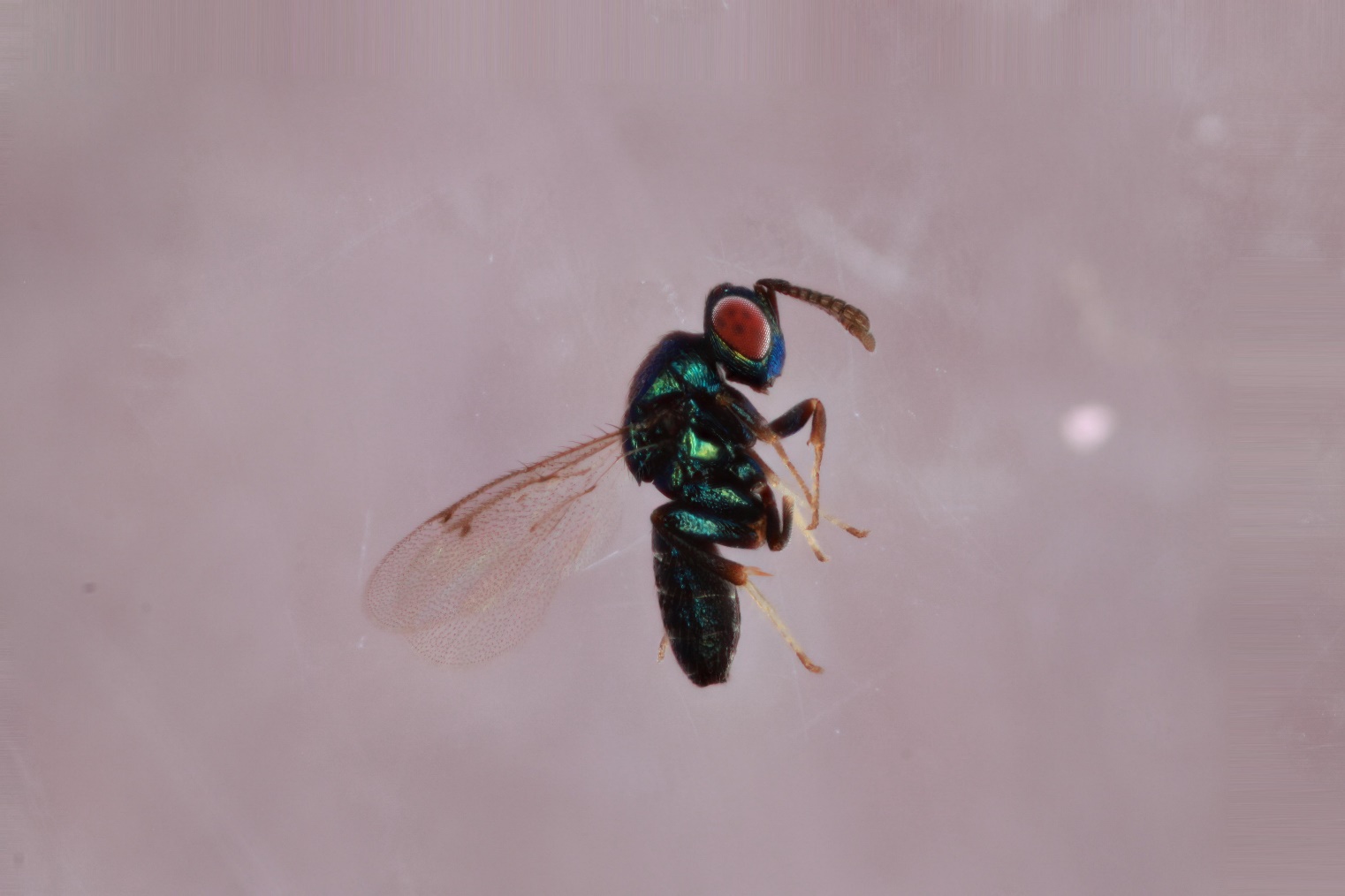

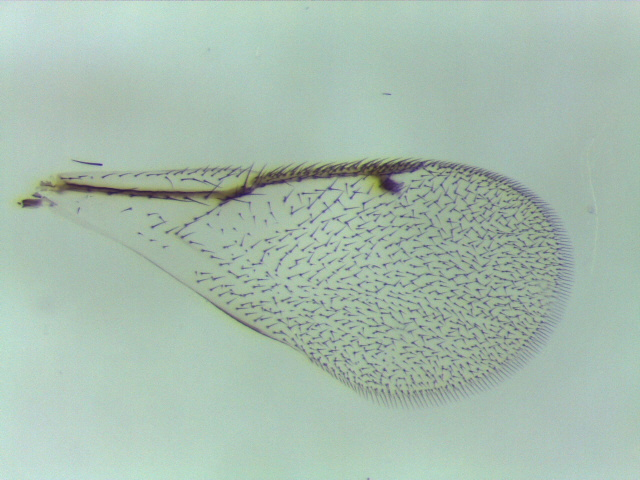

**Figure S25. *Ormyrus* *labotus* (clade 25):** 826-030-9A, female rom *Andricus chinquapin* from *Quercus bicolor* – Iowa City, IA.

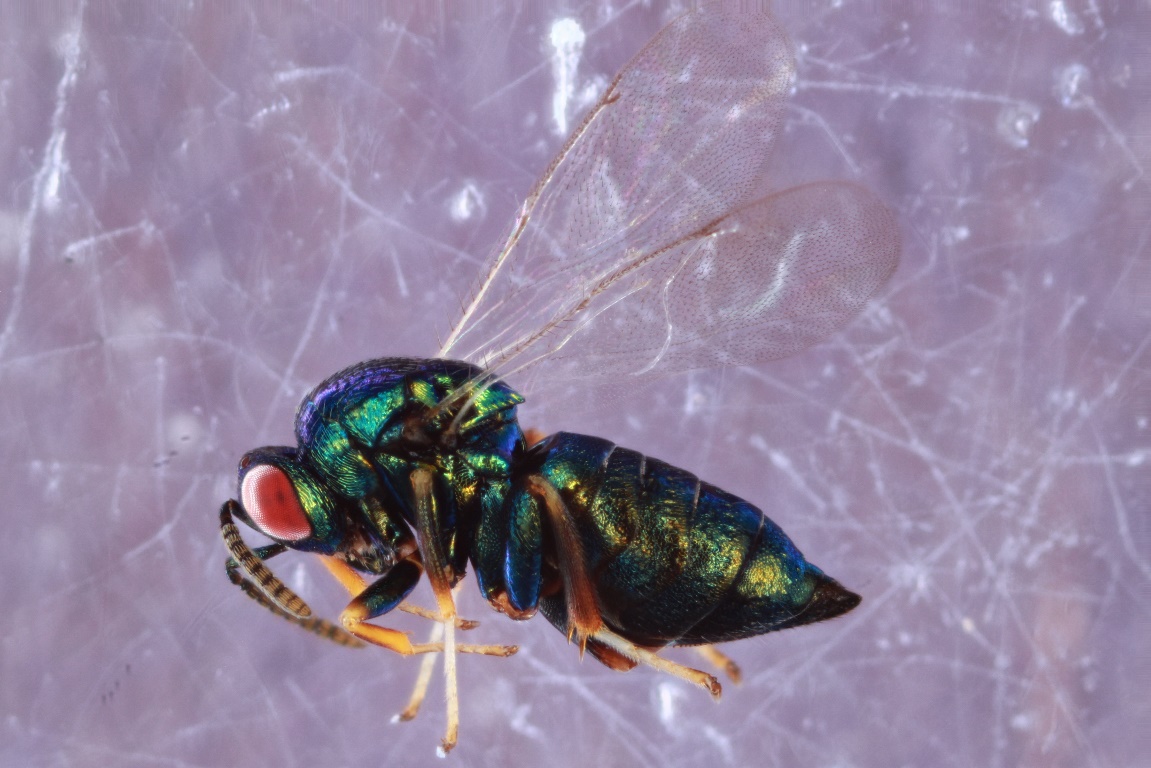

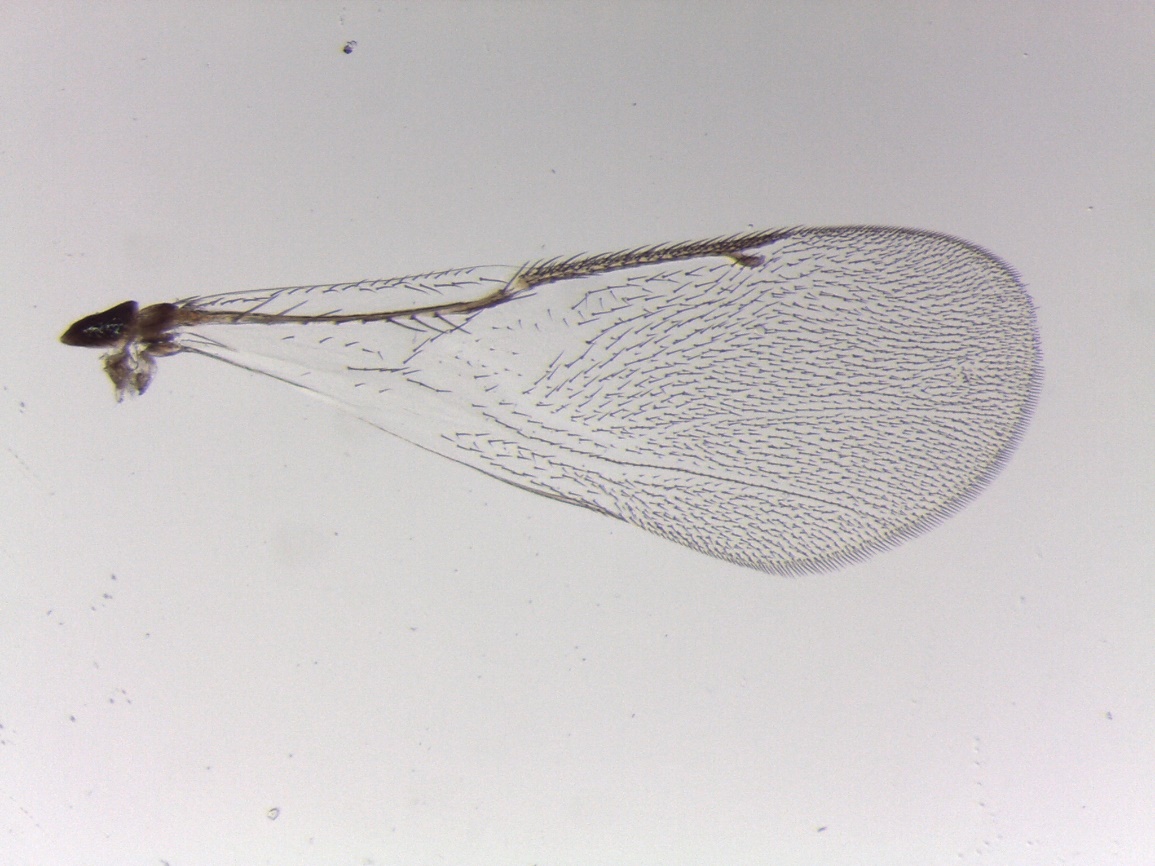

**Figure S26. *Ormyrus* *labotus* (clade 26a):** 1243-117-31, female from *Andricus quercuspetiolicola* on *Quercus stellata* – Austin, TX.

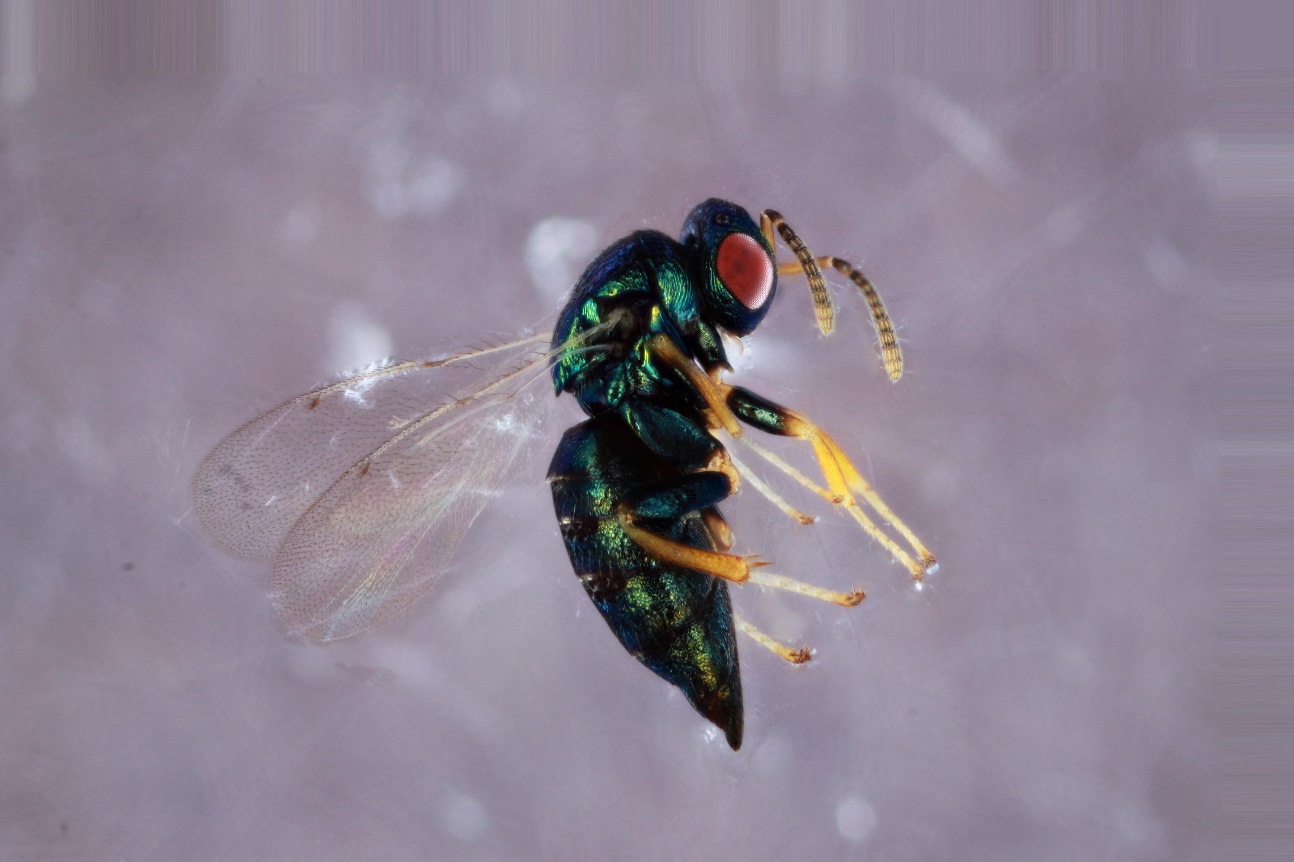

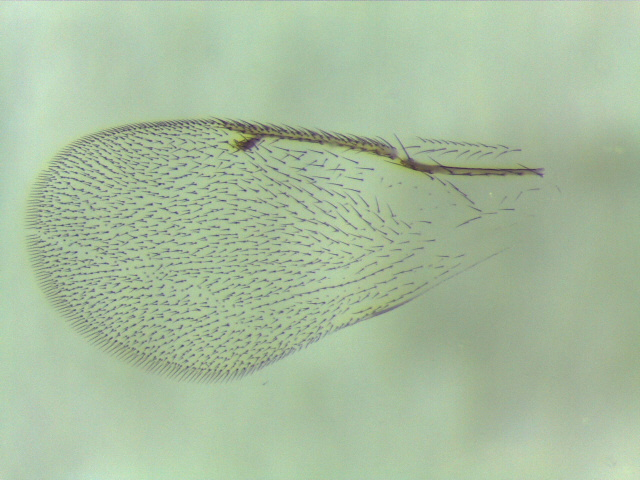

**Figure S27. *Ormyrus* *labotus* (clade 26b):** YMZ2, female from *Dryocosmus floridensis* on *Quercus laurifolia* – Gainesville, FL.

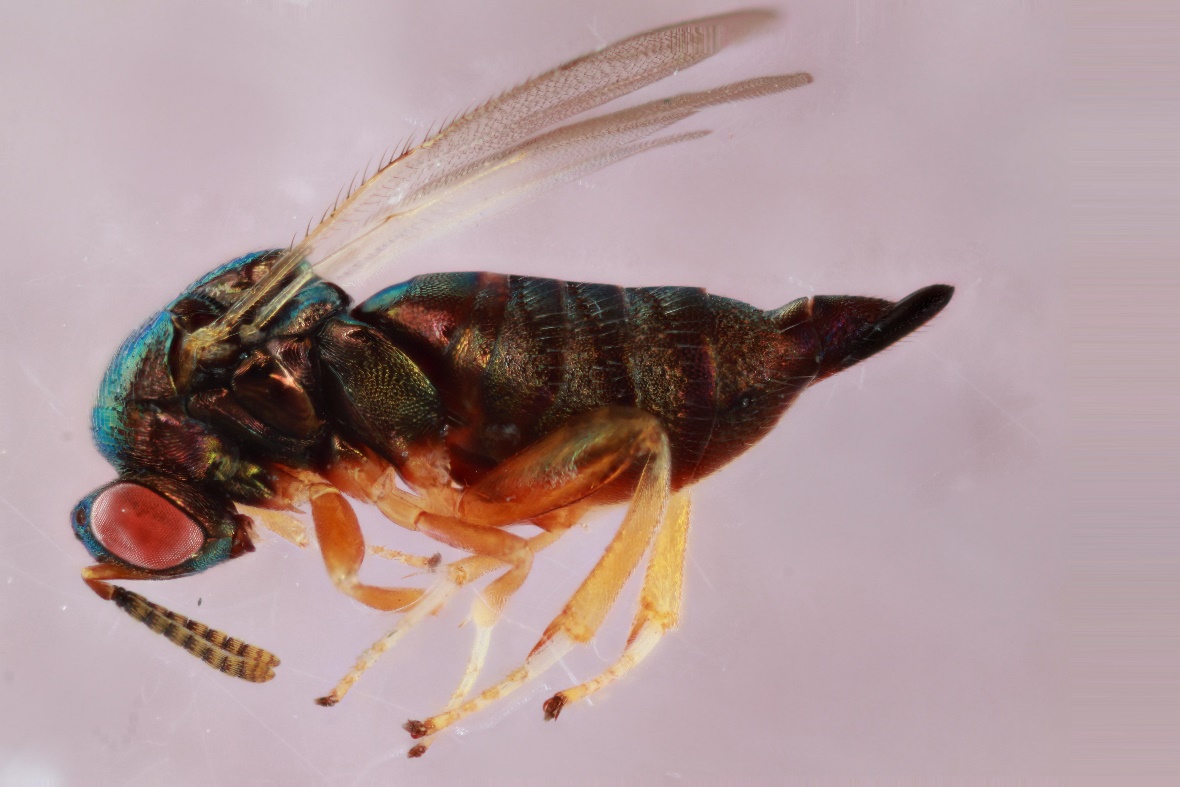

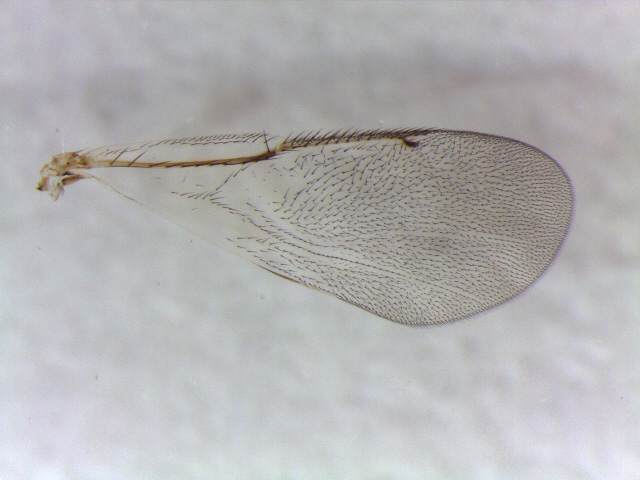

**Figure S28. *Ormyrus* *labotus* (clade 27)**: 1589-160-15A, female from *Andricus quercuslanigera* on *Quercus virginiana* – Lake Kyle, TX.

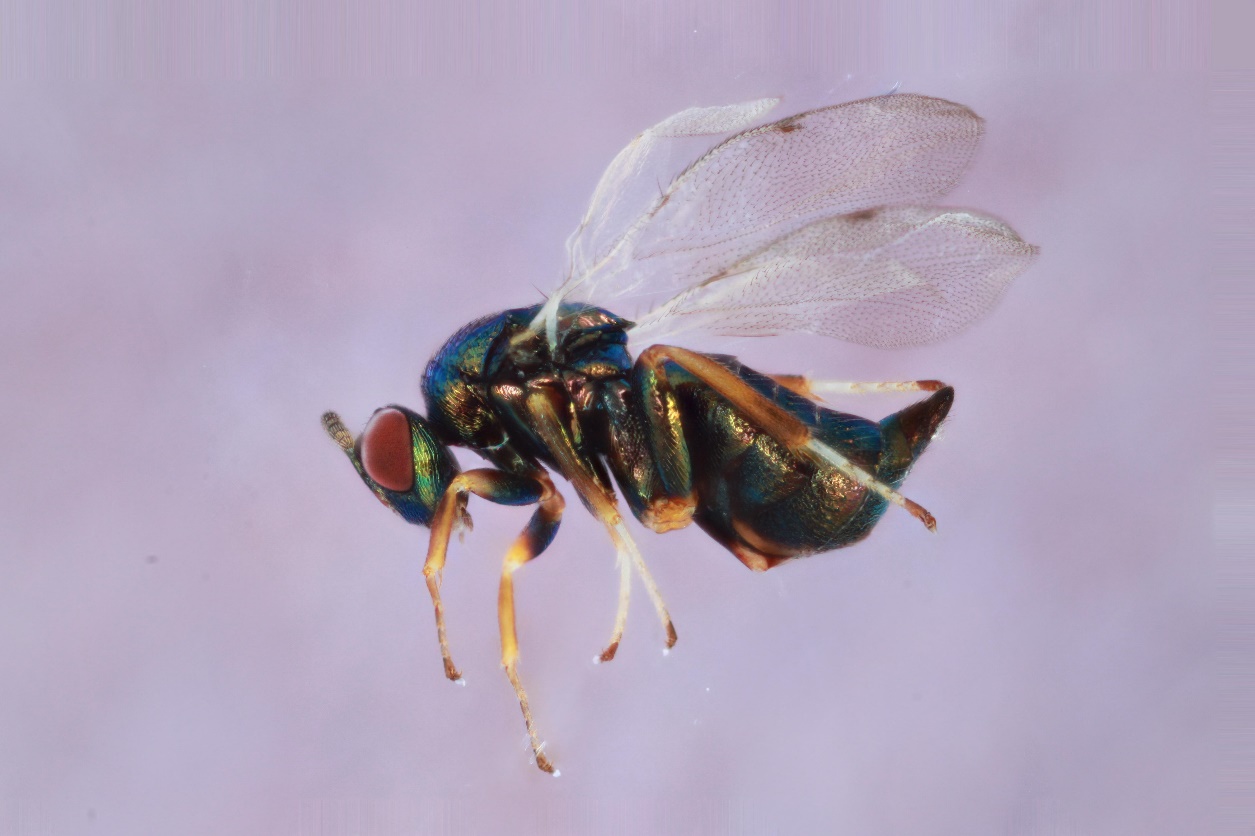

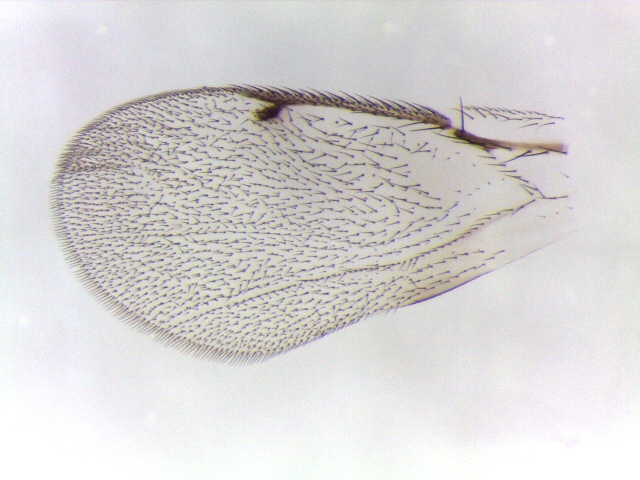

**Figure S29. *Ormyrus* *labotus* (clade 28):** P121, female (body photo only) from *Belonocnema treatae* on *Quercus fusiformis* – Ingleside, TX.

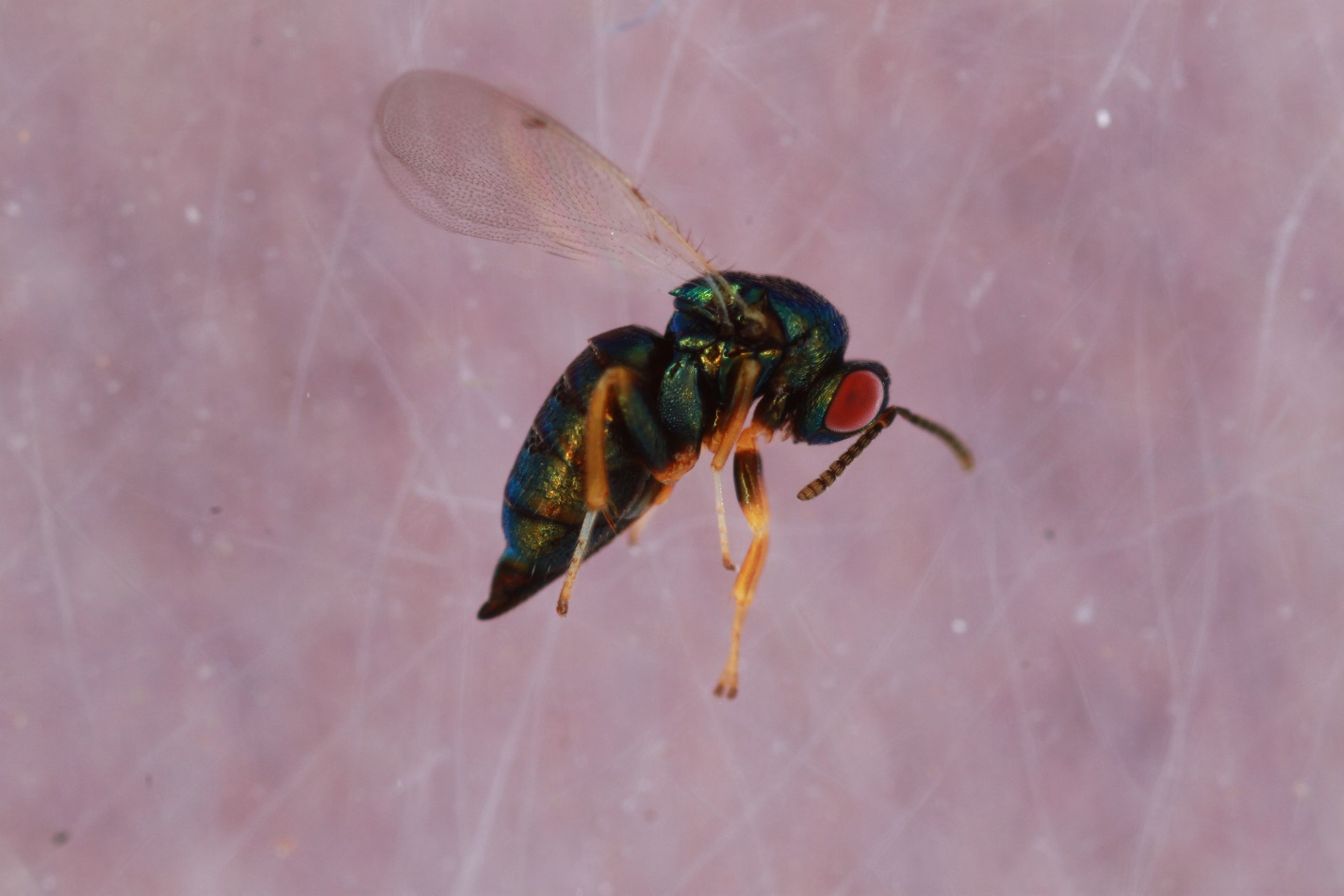

**Figure S30. *Ormyrus* *labotus* (“tigermorph”) (clade 29):** P184, female from *Belonocnema treatae on Quercus virginiana* – Gautier, MS.

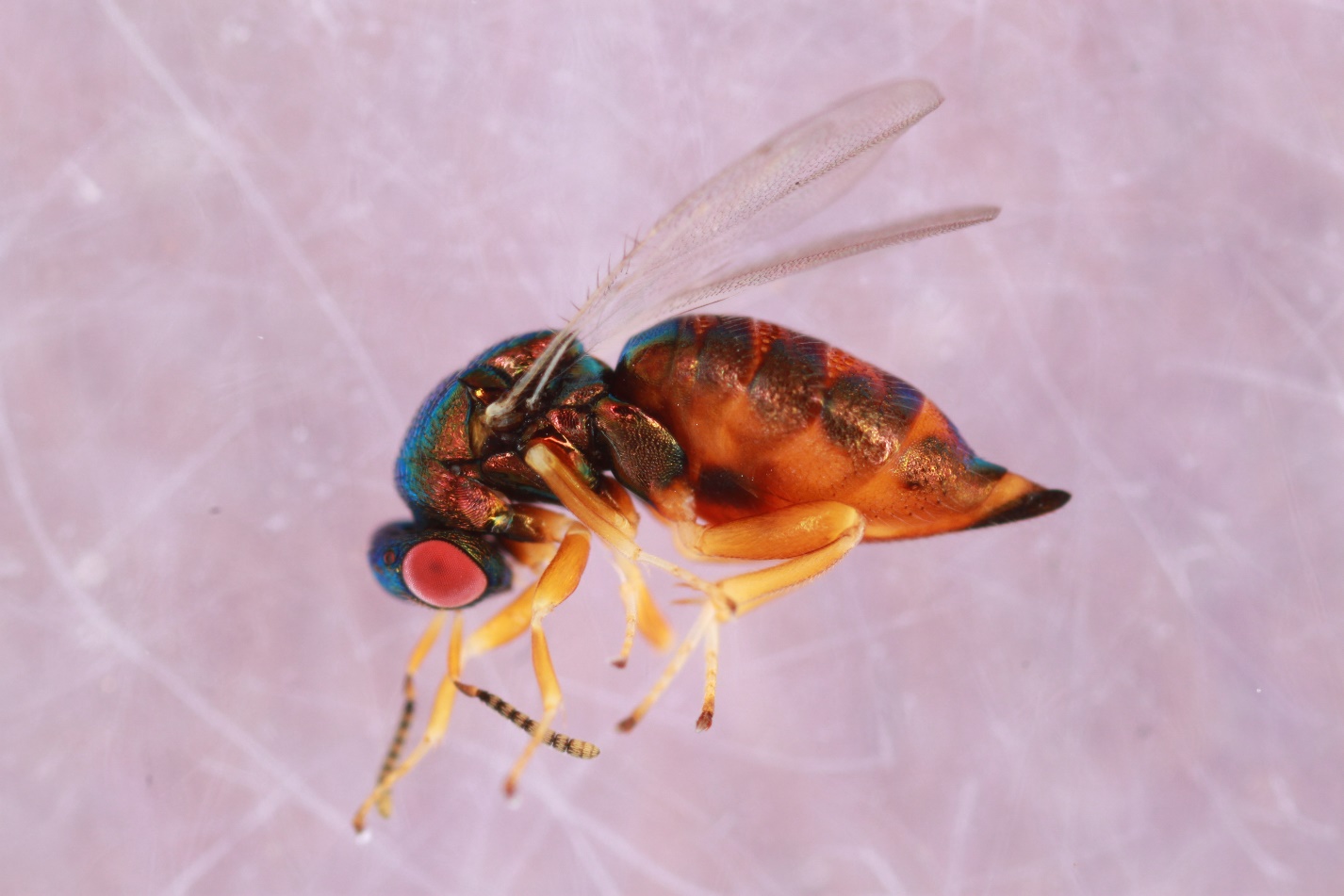

**Figure S31. *Ormyrus* *labotus* (clade 30):** 880-042-17G, female from *Andricus quercuspetiolicola* on *Quercus macrocarpa* – Oxford, IA.

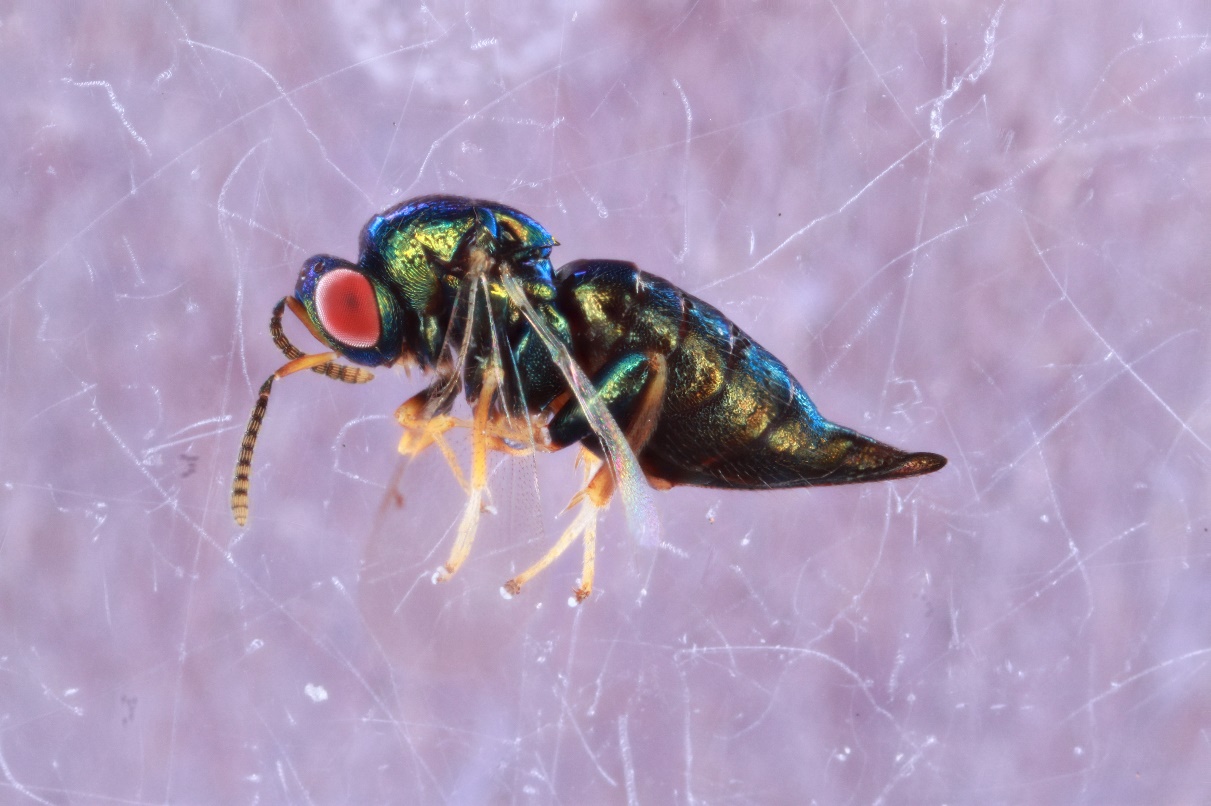

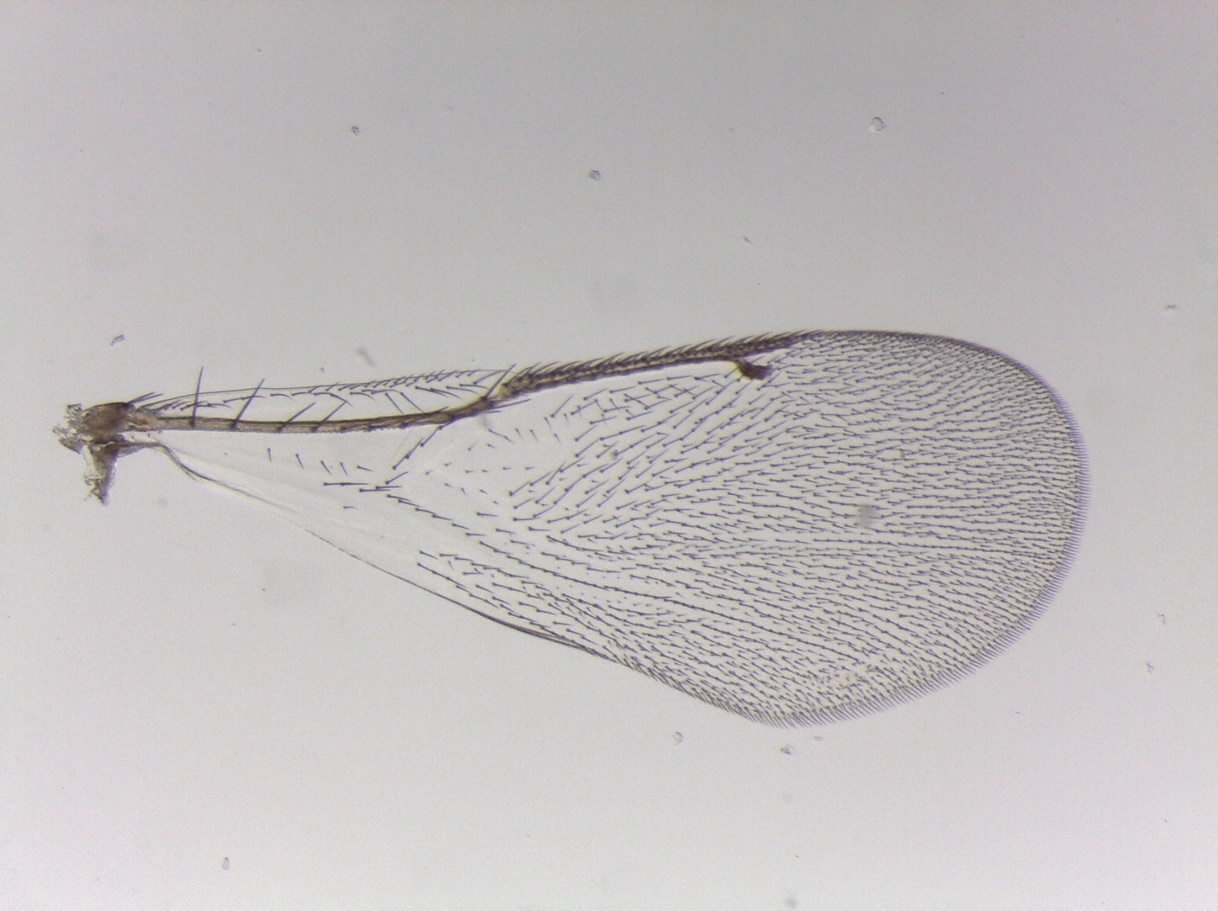

**Figure S32. *Ormyrus* *labotus* (clade 31)**: 1367-047-2, female from *Dryocosmus cinereae* on *Quercus imbricaria* – Iowa City, IA.

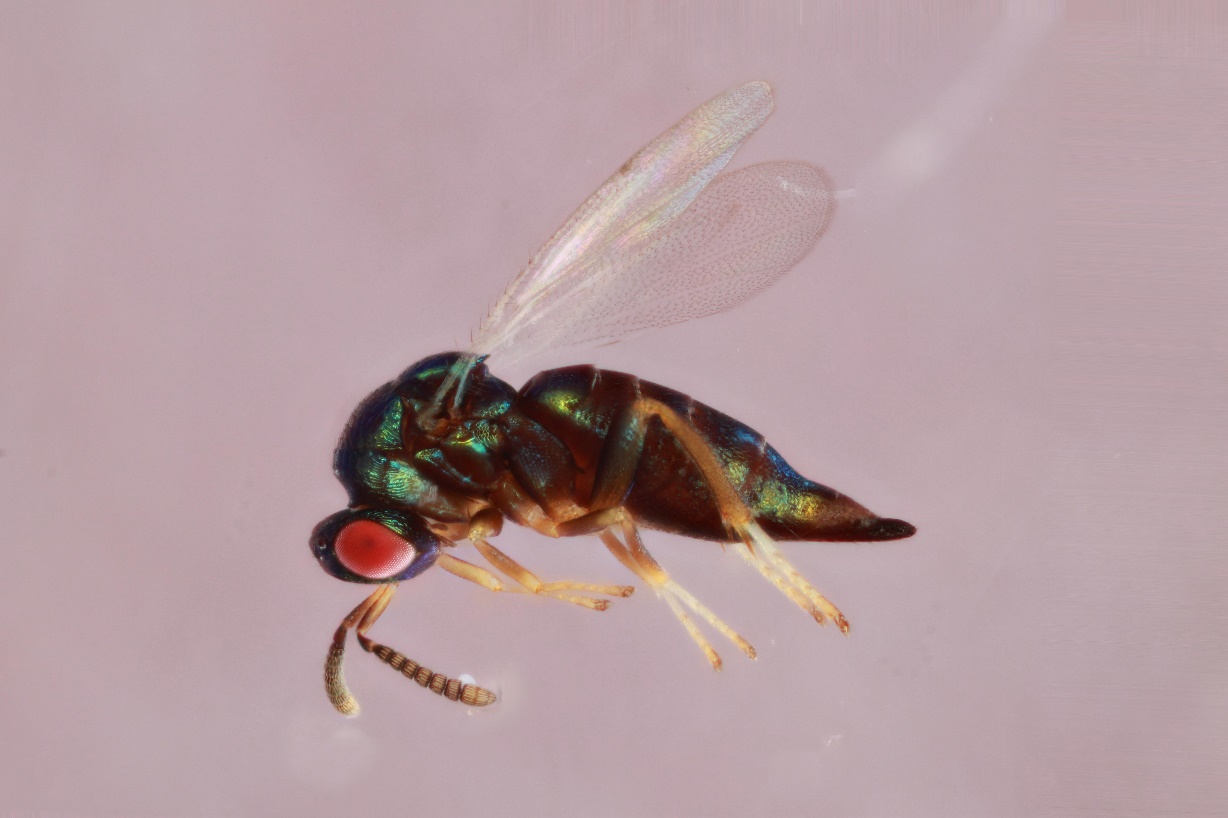

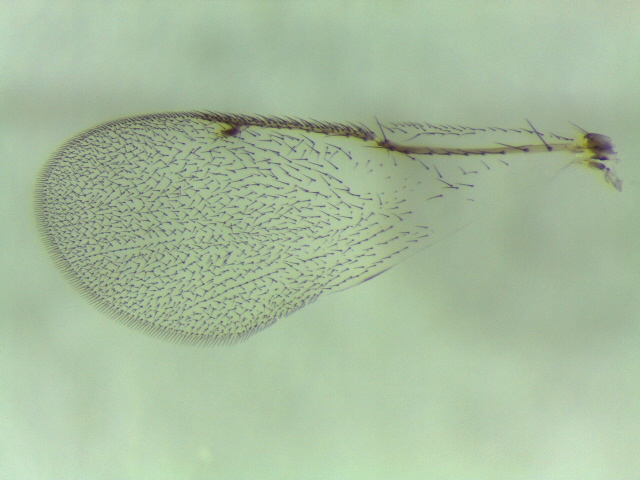

**Figure S33. *Ormyrus* *labotus* (clade 32):** 700-006-5, female from *Andricus quercusfrondosus* from Quercus bicolor – Iowa City, IA.

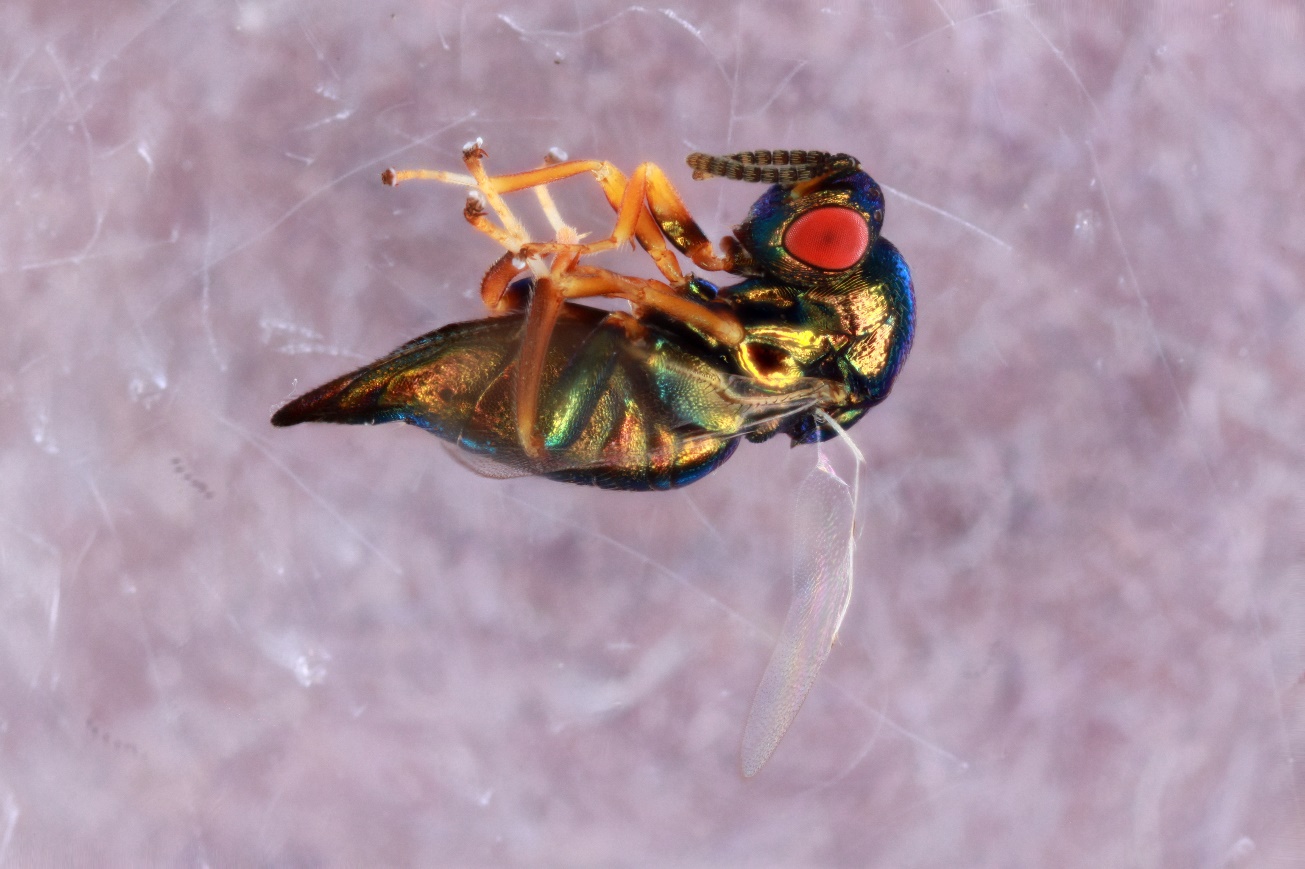

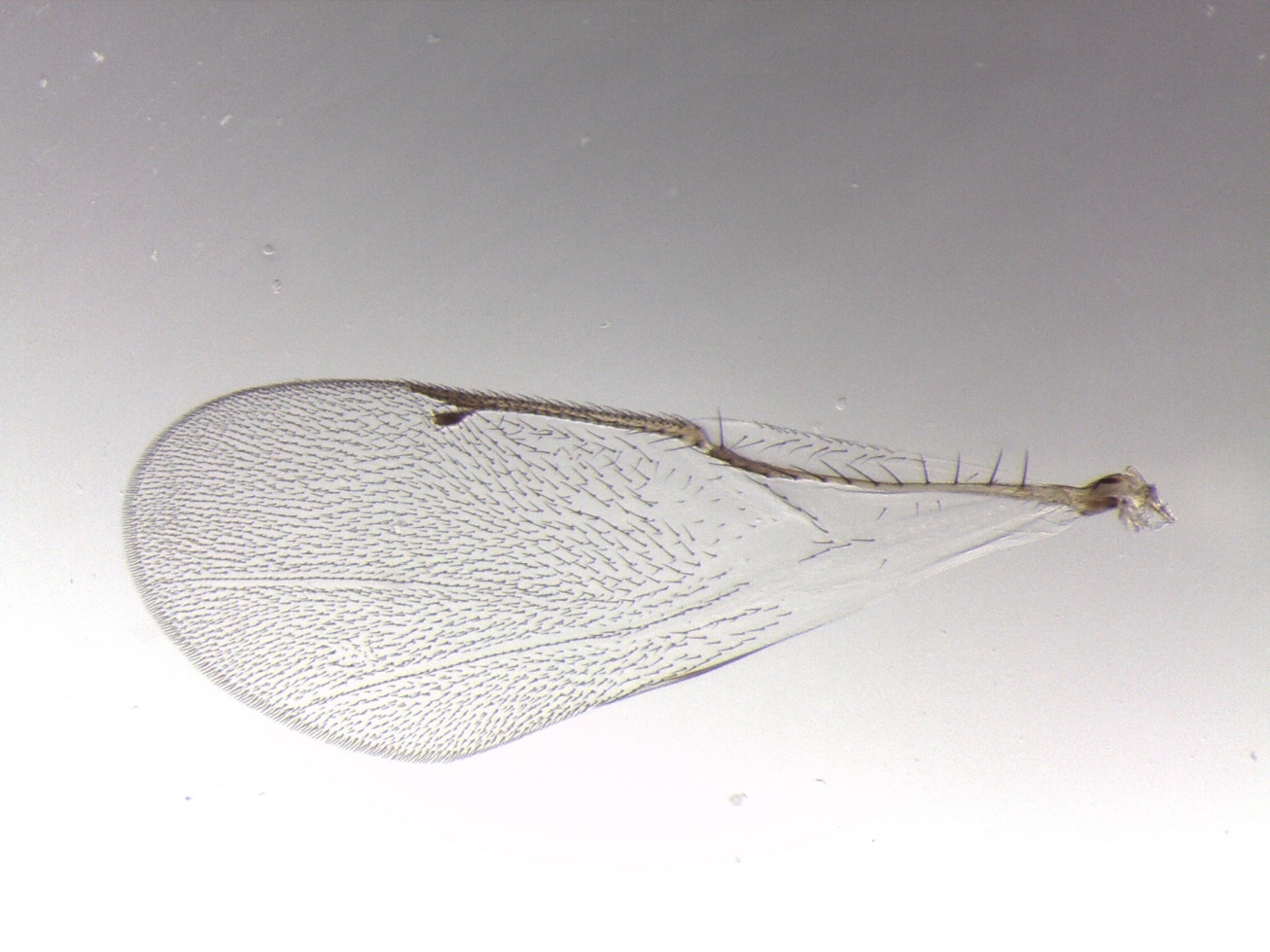

**Figure S34. *Ormyrus* *labotus* (clade 33):** 932-070-002, female from *Callirhytis cornigera* on *Quercus palustris* – St. Louis, MO.

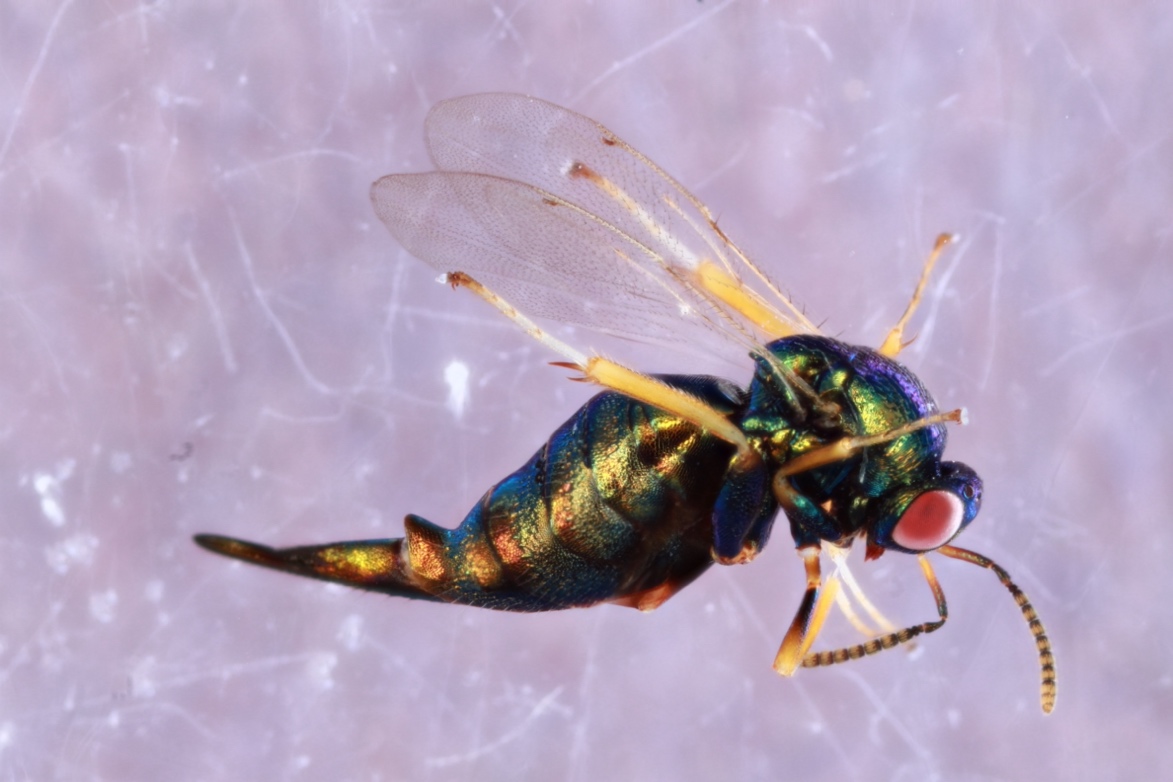

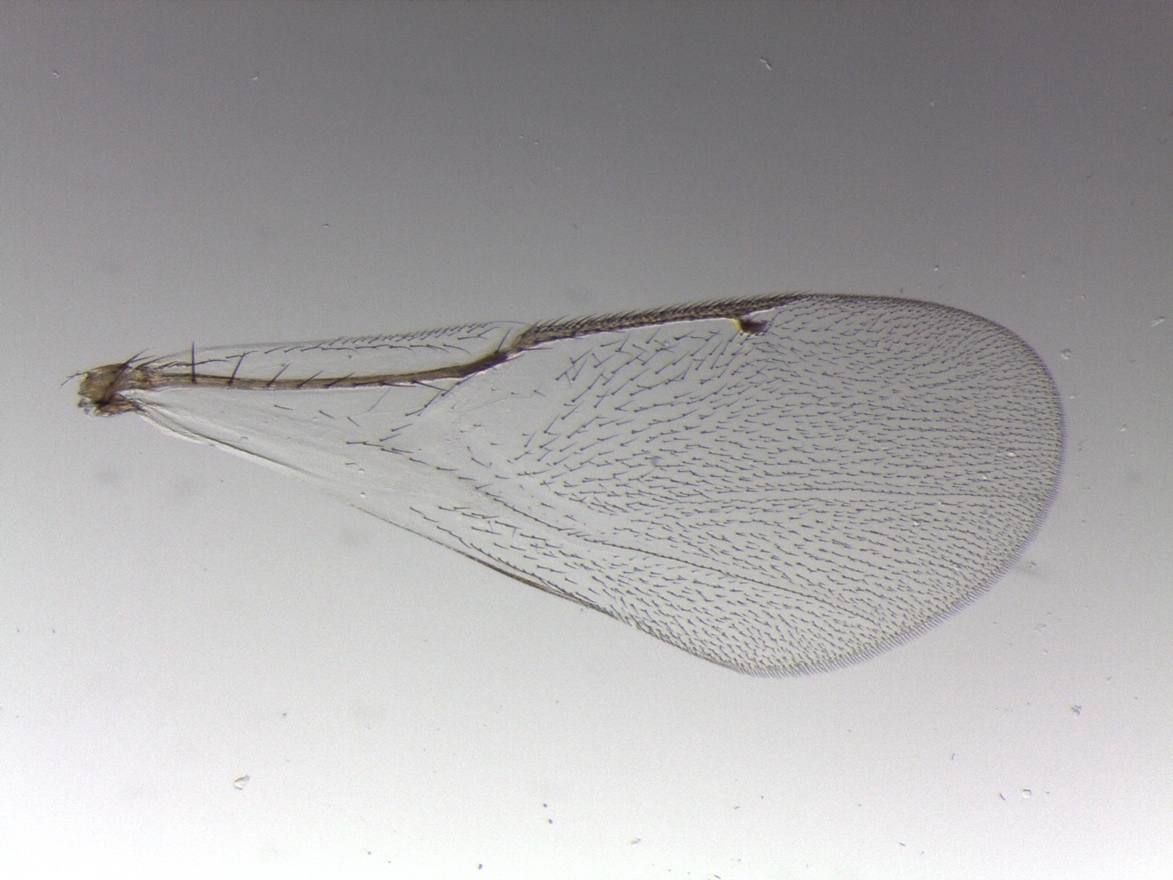

**Figure S35. *Ormyrus* *labotus* (clade 34):** JRO-1-5B, female from *Philonix nigra* on *Quercus alba* – Capon Bridge, WV.

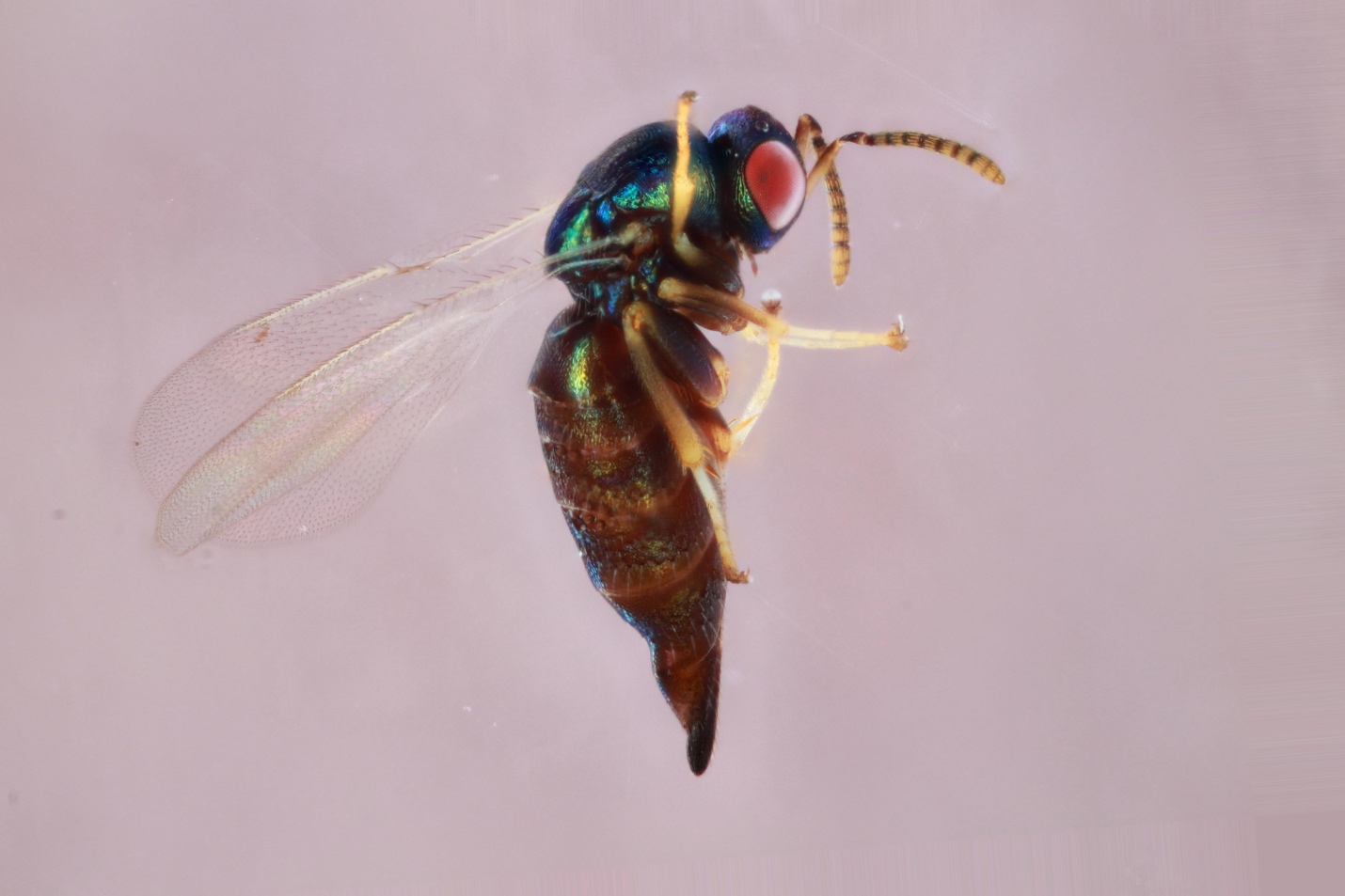

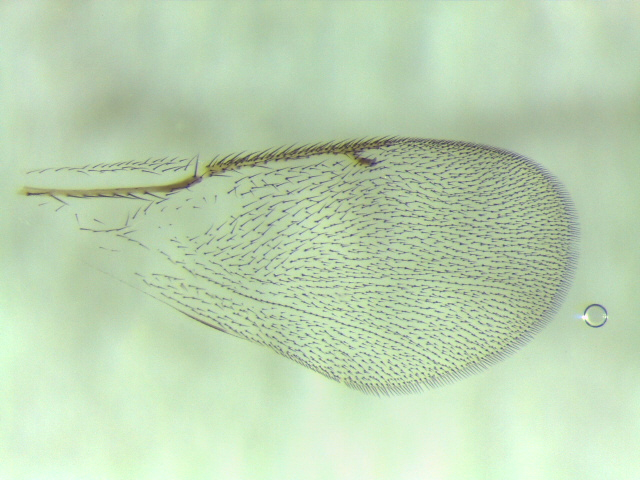

**Figure S36. *Ormyrus* *labotus* (clade 34):** 661-002-2A, *Acraspis erinacei* on *Quercus alba* – Oxford, IA.

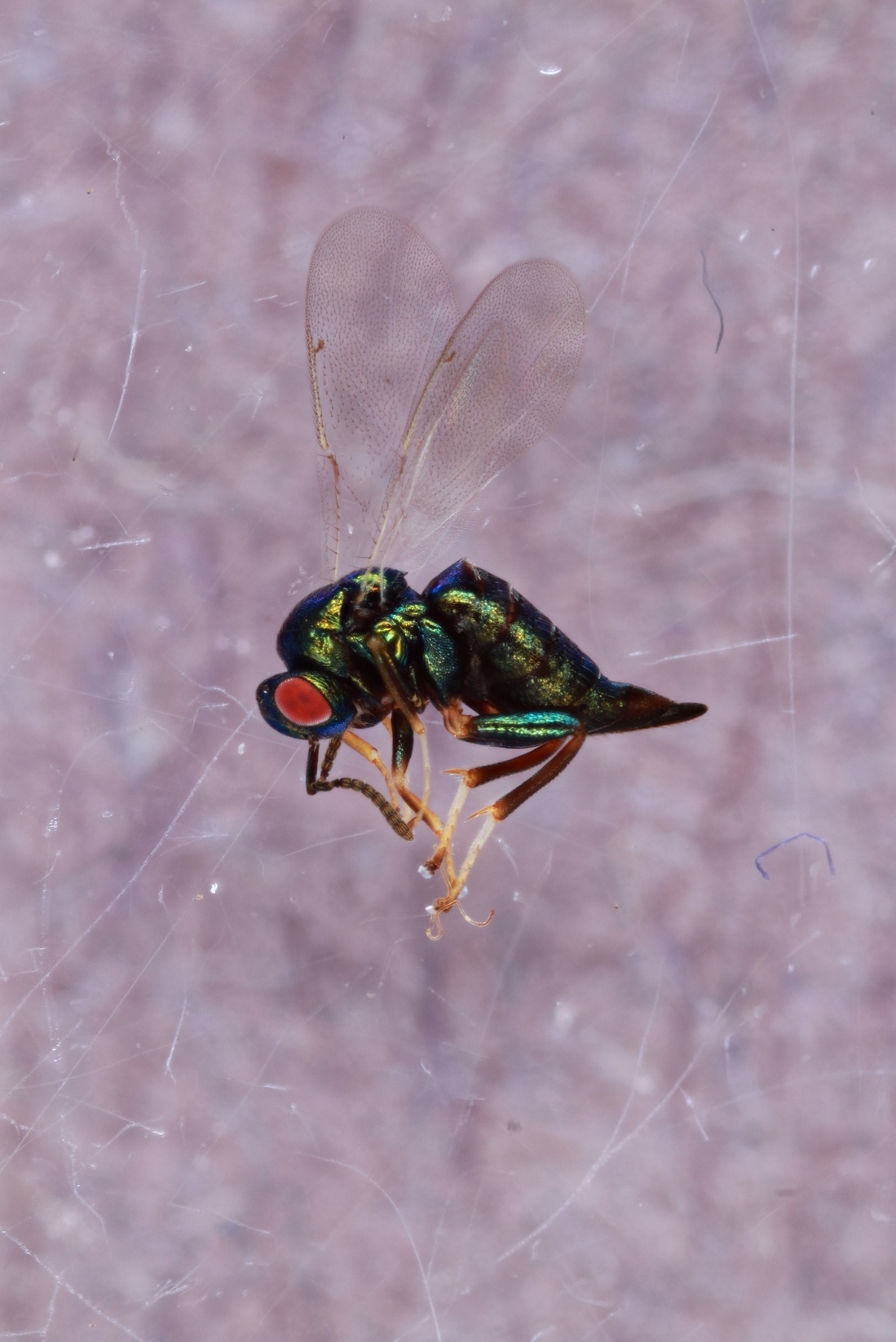

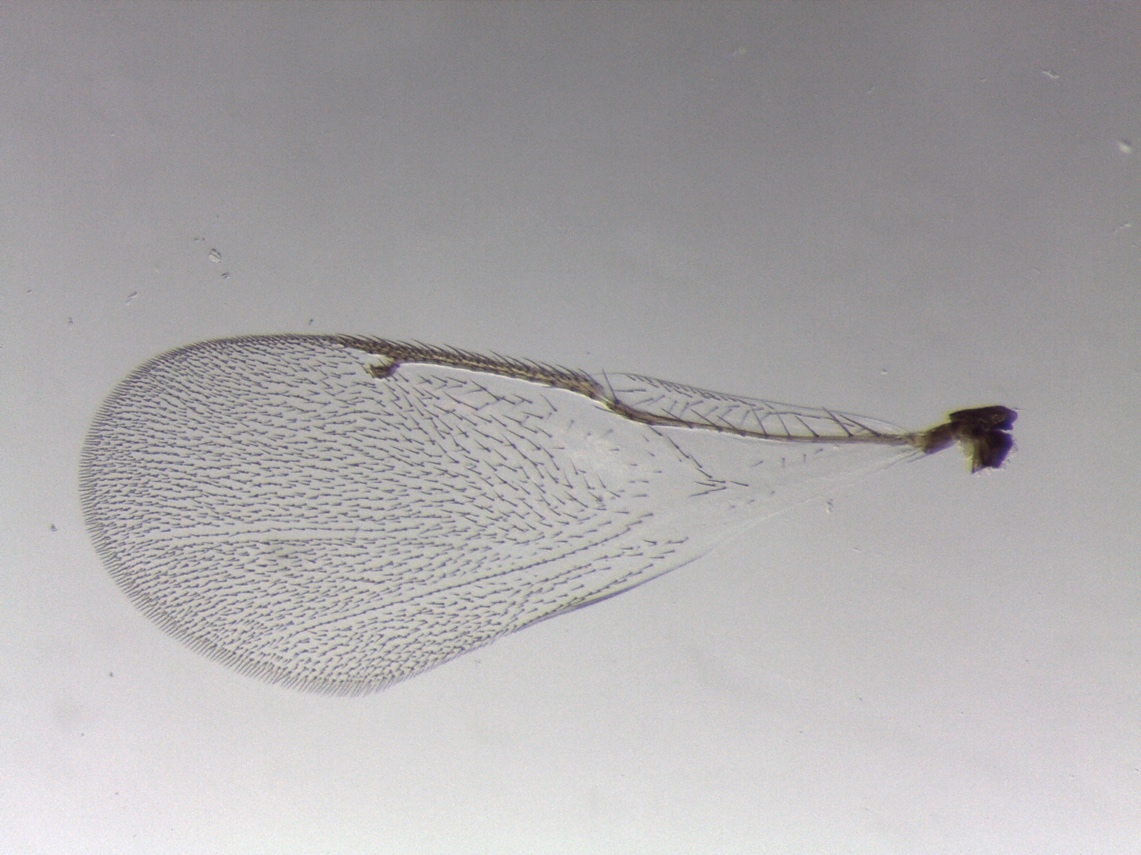

**Figure S37. *Ormyrus* *labotus* (clade 35):** 1181-003-2, female from *Acraspis pezomachoides* on *Quercus alba* – White Oak, PA.

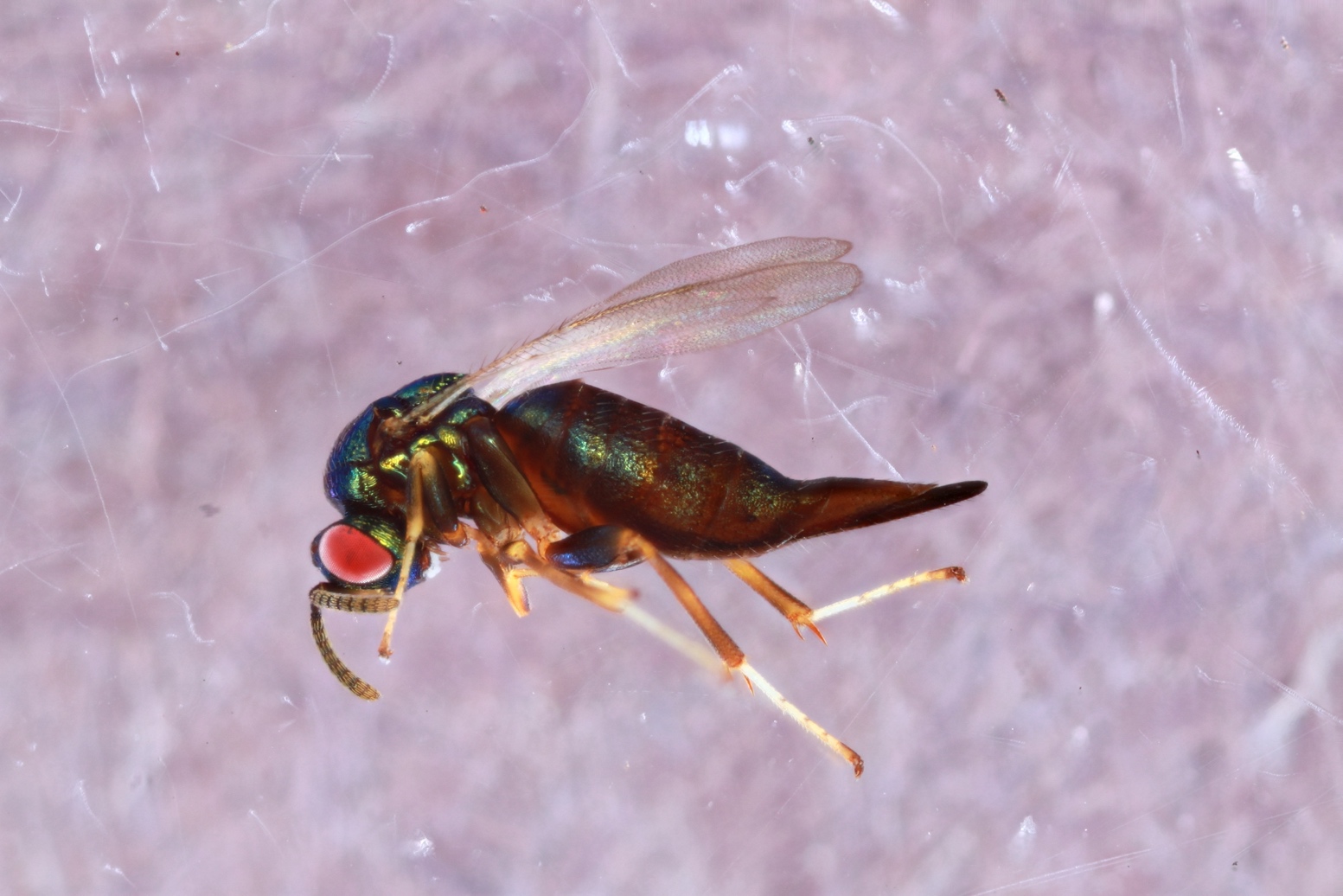

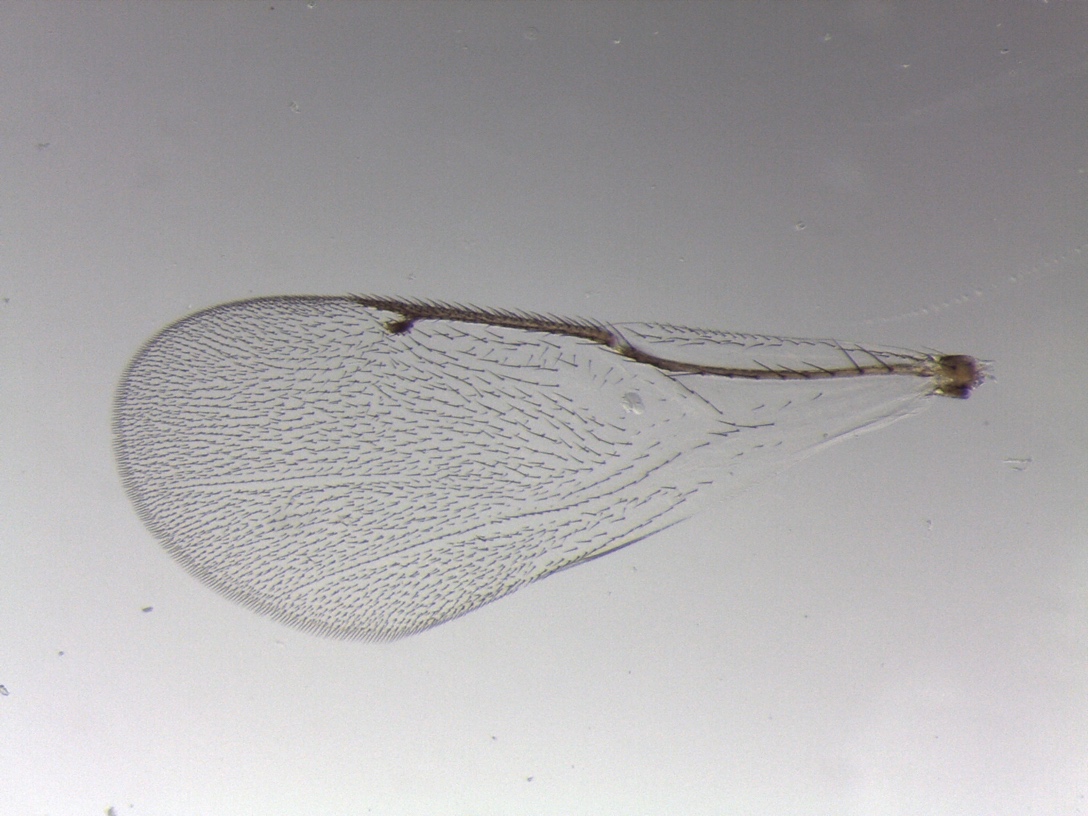

**Figure S38. *Ormyrus* *labotus* (clade 35):** 1566-002-5A, female from *Acraspis erinacei* on *Quercus alba* – Vestal, NY.
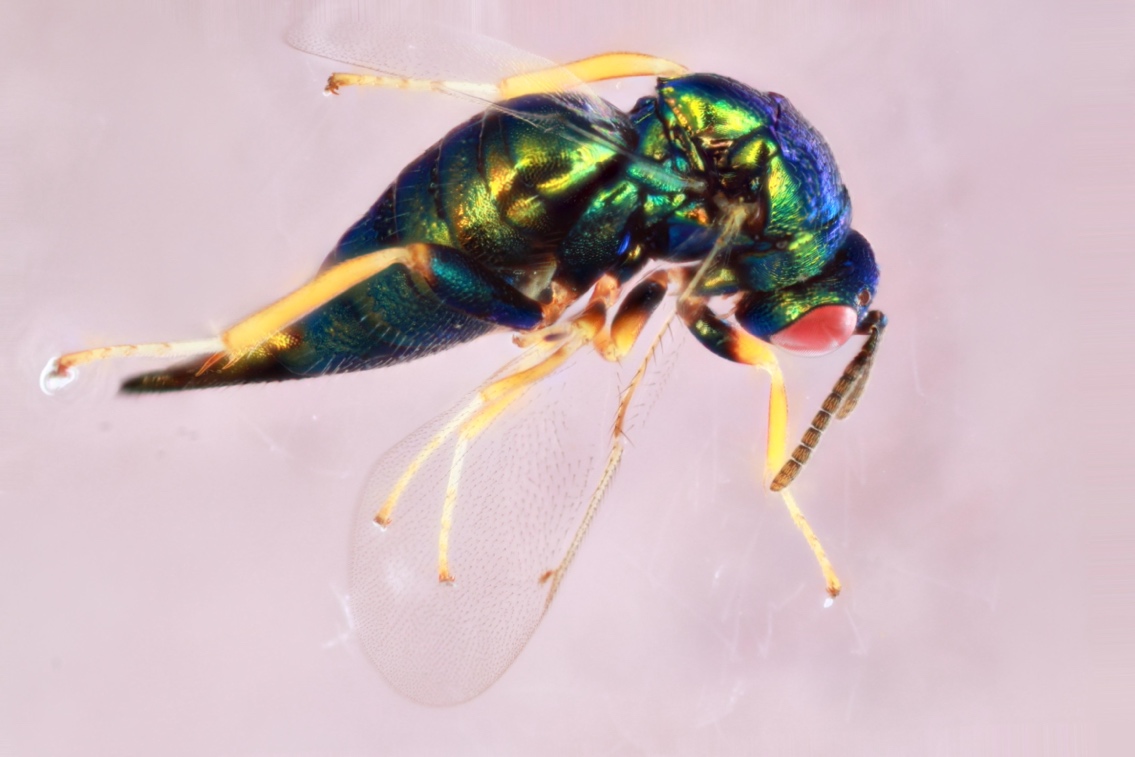

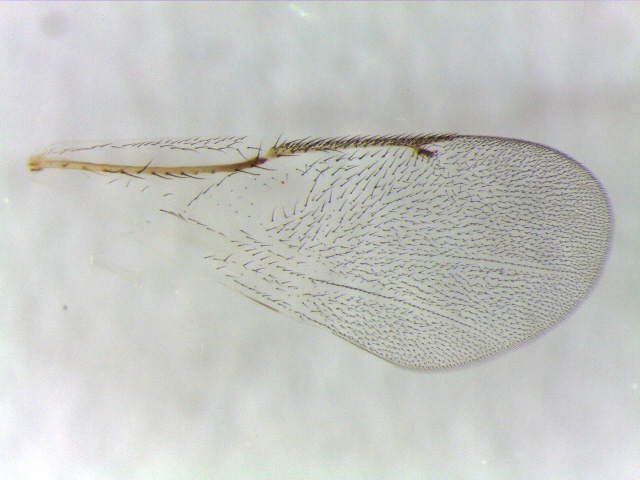

**Figure S39. *Ormyrus* *labotus* (clade 36):** 643-004-4, female from *Acraspis villosa* on *Quercus macrocarpa* – Iowa City, IA.

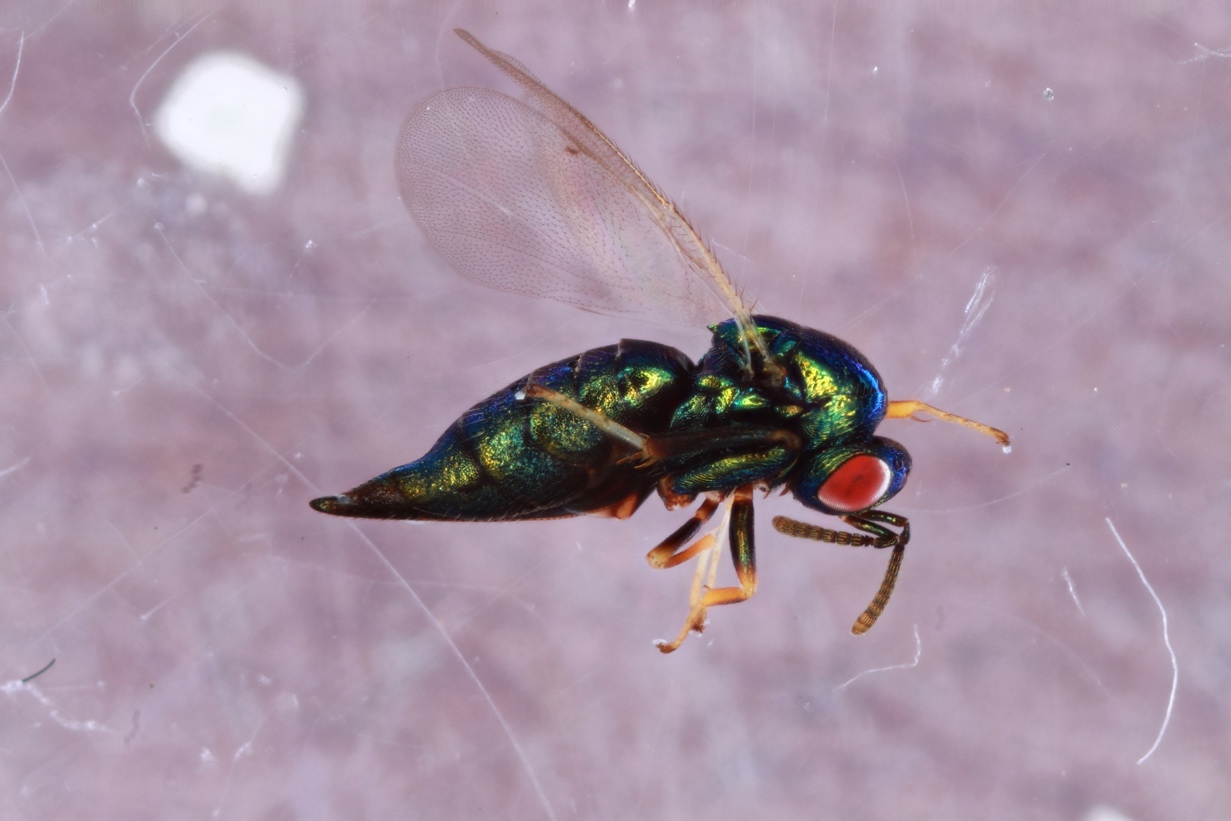

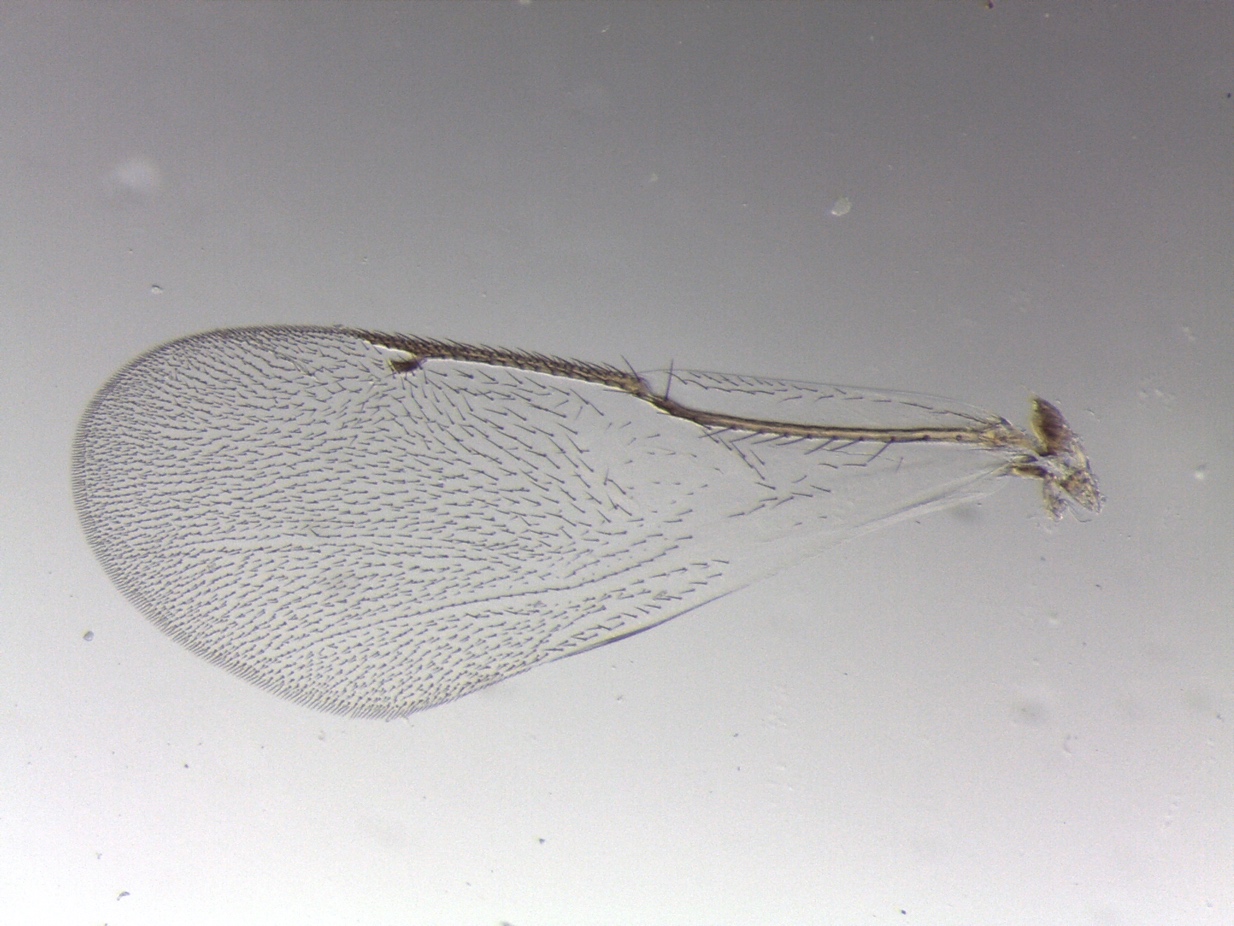
