## Supplementary figures and images for "*Ormyrus labotus* Walker (Hymenoptera: Ormyridae): another generalist that should not be a generalist is not a generalist"

### Supplemental Figure 40

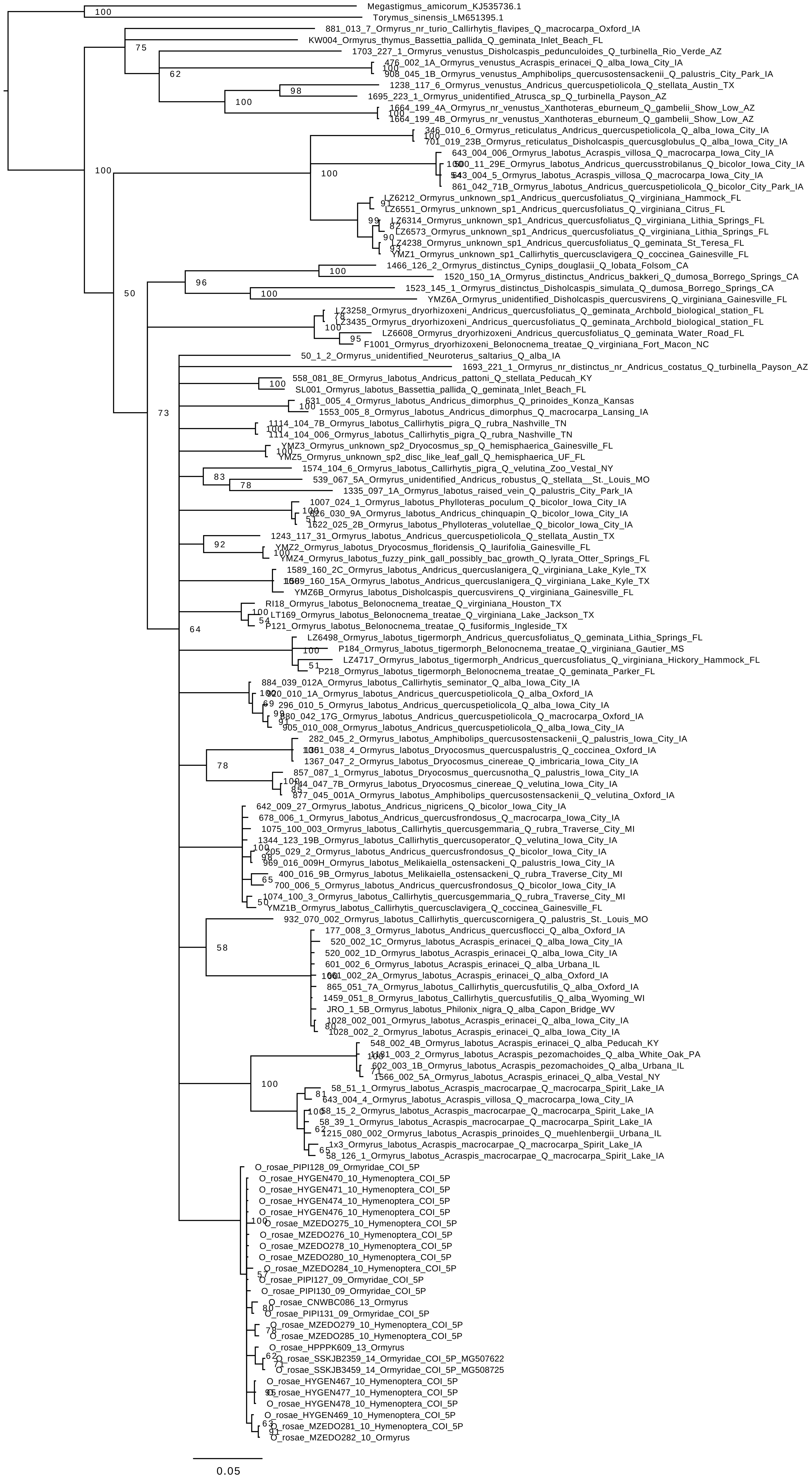

### Supplemental Figure 41

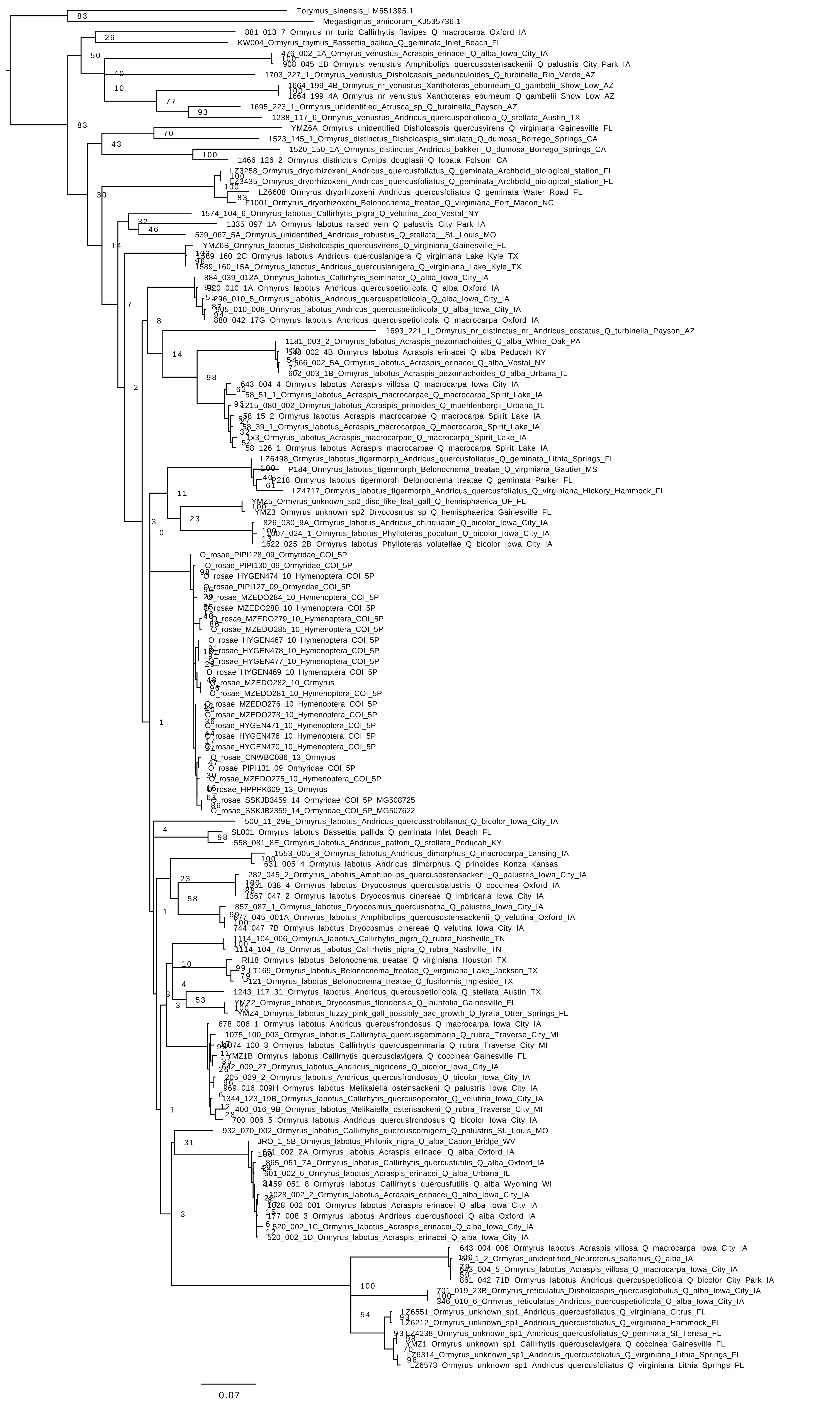
