## Supplemental Table 1 for "*Ormyrus labotus* Walker (Hymenoptera: Ormyridae): another generalist that should not be a generalist is not a generalist"

**Supplemental Table 1. *Ormyus labotus* host associations from Hanson 1992 and this study.**
An asterisk next to a name in the first column indicates a gall species from which we also reared an *Ormyrus labotus* included in this study.

| **Hanson 1992:** | **This study:** |
| --- | --- |
| *Acraspis alaria* | *Andricus chinquapin* |
| *Acraspis erinacei** | *Andricus dimorphus* |
| *Acraspis gemula* | *Andricus nigricens* |
| *Acraspis macrocarpae** | *Andricus quercusfrondosus* |
| *Acraspis pezomachoides** | *Andricus quercusstrobilanus* |
| *Acraspis prinoides** | *Belonocnema fossoria* |
| *Acraspis quercushirta* | *Belonocnema kinseyi* |
| *Acraspis villosa** | *Bassettia pallida* |
| *Amphibolips gainesi* | *Callirhytis pigra* |
| *Amphibolips nubilipennis* | *Callirhytis quercusclavigera* |
| *Amphibolips quercusoelebs* | *Disholcaspis quercusvirens* |
| *Amphibolips quercusspongifica* | *Dryocosmus cinerceae* |
| *Amphibolips tinctoriae* | *Phylloteras pocoulum* |
| *Andricus cinnamomeus* | *Phylloterus volutellae* |
| *Andricus coronus* | |
| *Andricus fullawayi* | |
| *Andricus ignotus** | |
| *Andricus pattoni** | |
| *Andricus quercusflocci** | |
| *Andricus quercusfoliatus** | |
| *Andricus quercuslanigera** | |
| *Andricus quercusostensackenii** | |
| *Andricus quercuspetiolicola** | |
| *Andricus quercussingularis* | |
| *Andricus tecturnarum* | |
| *Belonocnema treatae** | |
| *Callirhytis clavula* | |
| *Callirhytis cornigera** | |
| *Callirhytis elongata* | |
| *Callirhytis favosa* | |
| *Callirhytis flavipes* | |
| *Callirhytis gallaestriatae* | |
| *Callirhytis infuscata* | |
| *Callirhytis lanata* | |
| *Callirhytis pedunculata* | |
| *Callirhytis pulchra* | |
| *Callirhytis quercusfutilis** | |
| *Callirhytis quercusgemmaria** | |
| *Callirhytis quercusmedullae* | |
| *Melikaiella ostensackeni** | |
| *Callirhytis quercusoperator** | |

*Callirhytis quercuspunctata*
*Callirhytis quercusscitula*
*Callirhytis quercussimilis*
*Callirhytis seminator**
*Callirhytis tubicola*
*Callirhytis tumifica*
*Callirhytis vaccinii*
*Disholcaspis cinerosa*
*Disholcaspis spongiosa*
*Dryocosmus kuriphilus*
*Dryocosmus quercusnotha**
*Dryocosmus quercuspalustris**
*Loxaulus humilis*
*Loxaulus quercusmammula*
*Neuroterus exiquus*
*Neuroterus floccosus*
*Neuroterus quercusbatatus*
*Neuroterus quercusirregularis*
*Neuroterus quercusverrucarum*
*Philonix fulvicollis*
*Philonix nigra**
*Xanthoteras eburneum*
*Xanthoteras politum*
*Xanthoteras quercusforticorne*
