## Supplemental Table 2 for "*Ormyrus labotus* Walker (Hymenoptera: Ormyridae): another generalist that should not be a generalist is not a generalist"

**Supplemental Table 2.** ID numbers and collection information for Ormyrus included in this study

Unidentifed label means that we were unable to key out the individual; unknown_sp label means that representative individuals in the clade could not be keyed out to any existing species descriptions

Highlighted rows represent published oak-gall-associated *Ormyrus* sequences used in this study

| Genbank accession | Lab specific ID | Genus | Species | Clade on tree | Gall collected by | Gall Host | Tree Host | Collection Location | Date collected | Date emerged |
|---|---|---|---|---|---|---|---|---|---|---|
| OK624721 | 881_013_7 | *Ormyrus* | *nr turio* | 1 | A. Ward | *Callirhytis flavipes* | *Quercus macrocarpa* | Oxford, IA | 6/8/2017 | 7/6/2017 |
| MN935907.1 | KW004 | *Ormyrus* | *thymus* | 2 | | *Bassettia pallida* | *Quercus geminata* | Inlet Beach, FL | 8/1/2015 | |
| OK624757 | 1703_227_1 | *Ormyrus* | *venustus* | 3 | S. Sheikh | *Disholcaspis pedunculoides* | *Quercus turbinella* | Rio Verde, AZ | 9/17/2019 | 10/1/2019 |
| OK624696 | 476_002_1A | *Ormyrus* | *venustus* | 4 | A. Ward | *Acraspis erinacei* | *Quercus alba* | Iowa City, IA | 8/11/2016 | 8/25/2016 |
| OK624724 | 908_045_1B | *Ormyrus* | *venustus* | 4 | A. Ward | *Amphibolips quercusostensackenii* | *Quercus palustris* | City Park, IA | 6/16/2017 | 6/27/2017 |
| OK624737 | 1238_117_6 | *Ormyrus* | *venustus* | 5 | A. Forbes | *Andricus quercuspetiolicola* | *Quercus stellata* | Austin, TX | 4/12/2018 | 6/4/2018 |
| OK624756 | 1695_223_1 | *Ormyrus* | unidentified | 6 | S. Sheikh | Atrusca sp. | *Quercus turbinella* | Payson, AZ | 9/17/2019 | 11/20/2019 |
| OK624753 | 1664_199_4A | *Ormyrus* | *nr venustus* | 7 | S. Sheikh | *Xanthoteras eburneum* | *Quercus gambelii* | Show Low, AZ | 9/16/2019 | 11/25/2019 |
| OK624754 | 1664_199_4B | *Ormyrus* | *nr venustus* | 7 | S. Sheikh | *Xanthoteras eburneum* | *Quercus gambelii* | Show Low, AZ | 9/16/2019 | 11/25/2019 |
| OK624713 | 701_019_23B | *Ormyrus* | *reticulatus* | 8 | A. Ward | *Disholcaspis quercusglobulus* | *Quercus alba* | Iowa City, IA | 4/12/2017 | 5/25/2017 |
| OK624709 | 643_004_006 | *Ormyrus* | *labotus* | 9 | A. Ward | *Acraspis villosa* | *Quercus macrocarpa* | Iowa City, IA | 9/27/2016 | 7/23/2017 |
| OK624708 | 643_004_5 | *Ormyrus* | *labotus* | 9 | A. Ward | *Acraspis villosa* | *Quercus macrocarpa* | Iowa City, IA | 9/27/2016 | 5/20/2017 |
| OK624694 | 346_010_6 | *Ormyrus* | *reticulatus* | 8 | A. Ward | *Andricus quercuspetiolicola* | *Quercus alba* | Iowa City, IA | 6/24/2016 | 6/30/2016 |
| OK624717 | 861_042_71B | *Ormyrus* | *labotus* | 9 | A. Ward | *Andricus quercuspetiolicola* | *Quercus bicolor* | City Park, IA | 6/7/2017 | 7/5/2017 |
| OK624697 | 500_11_29E | *Ormyrus* | *labotus* | 9 | A. Ward | *Andricus quercusstrobilanus* | *Quercus bicolor* | Iowa City, IA | 8/18/2016 | 9/8/2016 |
| OK624764 | LZ6212 | *Ormyrus* | unknown_sp1 | 10 | L. Zhang | *Andricus quercusfoliatus* | *Quercus virginiana* | Hammock, FL | 12/15/2018 | 3/4/2019 |
| OK624765 | LZ6551 | *Ormyrus* | unknown_sp1 | 10 | L. Zhang | *Andricus quercusfoliatus* | *Quercus virginiana* | Citrus, FL | 12/17/2018 | 4/15/2019 |
| OK624759 | LZ6314 | *Ormyrus* | unknown_sp1 | 10 | L. Zhang | *Andricus quercusfoliatus* | *Quercus virginiana* | Lithia Springs, FL | 12/16/2018 | 3/15/2019 |
| OK624763 | LZ6573 | *Ormyrus* | unknown_sp1 | 10 | L. Zhang | *Andricus quercusfoliatus* | *Quercus virginiana* | Lithia Springs, FL | 12/16/2018 | 4/19/2019 |
| OK624767 | LZ4238 | *Ormyrus* | unknown_sp1 | 10 | L. Zhang | *Andricus quercusfoliatus* | *Quercus geminata* | St. Teresa, FL | 12/11/2017 | 9/26/2018 |
| OK624773 | YMZ1 | *Ormyrus* | unknown_sp1 | 10 | Y. M. Zhang | *Callirhytis quercusclavigera* | *Quercus coccinea* | Gainesville, FL | 11/3/2019 | |
| OK624744 | 1466_126_2 | *Ormyrus* | *distinctus* | 11 | A. Forbes | *Cynips douglasii* | *Quercus lobata* | Folsom, CA | 7/31/2018 | 8/19/2018 |
| OK624745 | 1520_150_1A | *Ormyrus* | *distinctus* | 12 | A. Forbes | *Andricus bakkeri* | *Quercus dumosa* | Borrego Springs, CA | 8/5/2018 | 9/2/2018 |
| OK624746 | 1523_145_1 | *Ormyrus* | *distinctus* | 13 | A. Forbes | *Disholcaspis simulata* | *Quercus dumosa* | Borrego Springs, CA | 8/5/2018 | 9/13/2018 |
| OK624779 | YMZ6A | *Ormyrus* | *unidentified* | 14 | Y. M. Zhang | *Disholcaspis quercusvirens* | *Quercus virginiana* | Gainesville, FL | | |
| OK624761 | LZ3258 | *Ormyrus* | *dryorhizoxeni* | 15 | L. Zhang | *Andricus quercusfoliatus* | *Quercus geminata* | Archbold biological station, FL | 12/14/2017 | 4/5/2018 |
| OK624762 | LZ3435 | *Ormyrus* | *dryorhizoxeni* | 15 | L. Zhang | *Andricus quercusfoliatus* | *Quercus geminata* | Archbold biological station, FL | 12/14/2017 | 4/12/2018 |
| OK624768 | LZ6608 | *Ormyrus* | *dryorhizoxeni* | 15 | L. Zhang | *Andricus quercusfoliatus* | *Quercus geminata* | Water Road, FL | 12/19/2018 | 4/24/2019 |
| OK624769 | F1001 | *Ormyrus* | *dryorhizoxeni* | 15 | | *Belonocnema treatae* | *Quercus virginiana* | Fort Macon, NC | | |
| OK624685 | 50_1_2 | *Ormyrus* | unidentified | 16 | E. Tvedte | *Neuroterus saltarius* | *Quercus alba* | IA | | |
| OK624755 | 1693_221_1 | *Ormyrus* | *nr distinctus* | 17 | S. Sheikh | *nr Andricus costatus* | *Quercus turbinella* | Payson, AZ | 9/17/2019 | 9/30/2019 |
| OK624702 | 558_081_8E | *Ormyrus* | *labotus* | 18 | A. Ward | *Andricus pattoni* | *Quercus stellata* | Peducah, KY | 9/3/2016 | 9/18/2016 |
| MN935904 | SL001 | *Ormyrus* | *labotus* | 18 | | *Bassettia pallida* | *Quercus geminata* | Inlet Beach, FL | 8/1/2015 | |
| OK624705 | 631_005_4 | *Ormyrus* | *labotus* | 19 | A. Hippee | *Andricus dimorphus* | *Quercus prinoides* | Konza, Kansas | 9/24/2016 | 11/6/2017 |
| OK624747 | 1553_005_8 | *Ormyrus* | *labotus* | 19 | A. Ward, S. Sheikh | *Andricus dimorphus* | *Quercus macrocarpa* | Lansing, IA | 10/6/2018 | 8/7/2019 |
| OK624733 | 1114_104_7B | *Ormyrus* | *labotus* | 20 | A. Ward | *Callirhytis pigra* | *Quercus rubra* | Nashville, TN | 8/25/2017 | 9/15/2017 |
| OK624734 | 1114_104_006 | *Ormyrus* | *labotus* | 20 | A. Ward | *Callirhytis pigra* | *Quercus rubra* | Nashville, TN | 8/25/2017 | 9/14/2017 |
| OK624778 | YMZ5 | *Ormyrus* | unknown_sp2 | 21 | Y. M. Zhang | disc like leaf gall | *Quercus hemisphaerica* | UF, FL | 3/20/2020 | |
| OK624776 | YMZ3 | *Ormyrus* | unknown_sp2 | 21 | Y. M. Zhang | *Dryocosmus sp* | *Quercus hemisphaerica* | Gainesville, FL | | |
| OK624749 | 1574_104_6 | *Ormyrus* | *labotus* | 22 | A. Forbes | *Callirhytis pigra* | *Quercus velutina* | Vestal, NY | 10/17/2018 | 10/29/2018 |
| OK624700 | 539_067_5A | *Ormyrus* | unidentified | 23 | A. Ward | *Andricus robustus* | *Quercus stellata* | St. Louis, MO | 9/2/2016 | 9/18/2016 |
| OK624739 | 1335_097_1A | *Ormyrus* | *labotus* | 24 | A. Ward | raised vein | *Quercus palustris* | City Park, IA | 5/16/2018 | 6/10/2018 |
| OK624715 | 826_030_9A | *Ormyrus* | *labotus* | 25 | A. Ward | *Andricus chinquapin* | *Quercus bicolor* | Iowa City, IA | 6/1/2017 | 6/17/2017 |
| OK624728 | 1007_024_1 | *Ormyrus* | *labotus* | 25 | A. Forbes | *Phylloteras poculum* | *Quercus bicolor* | Iowa City, IA | 7/14/2017 | 8/3/2017 |
| OK624752 | 1622_025_2B | *Ormyrus* | *labotus* | 25 | A. Ward, S. Sheikh | *Phylloteras volutellae* | *Quercus bicolor* | Iowa City, IA | 9/4/2019 | 9/17/2019 |
| OK624738 | 1243_117_31 | *Ormyrus* | *labotus* | 26a | A. Forbes | *Andricus quercuspetiolicola* | *Quercus stellata* | Austin, TX | 4/12/2018 | 9/9/2018 |
| OK624775 | YMZ2 | *Ormyrus* | *labotus* | 26b | Y. M. Zhang | *Dryocosmus floridensis* | *Quercus laurifolia* | Gainesville, FL | | |
| OK624777 | YMZ4 | *Ormyrus* | *labotus* | 26b | Y. M. Zhang | fuzzy pink gall possibly bac growth | *Quercus lyrata* | Otter Springs, FL | 1/23/2020 | |
| OK624750 | 1589_160_2C | *Ormyrus* | *labotus* | 27 | A. Ward | *Andricus quercuslanigera* | *Quercus fusiformis* | Lake Kyle, TX | 11/1/2018 | 11/12/2018 |
| OK624751 | 1589_160_15A | *Ormyrus* | *labotus* | 27 | A. Ward | *Andricus quercuslanigera* | *Quercus fusiformis* | Lake Kyle, TX | 11/1/2018 | 4/3/2019 |
| OK624780 | YMZ6B | *Ormyrus* | *labotus* | 27 | Y. M. Zhang | *Disholcaspis quercusvirens* | *Quercus virginiana* | Gainesville, FL | | |
| KR108716 | RI18 | *Ormyrus* | *labotus* | 28 | | *Belonocnema kinseyi* | *Quercus virginiana* | Houston, TX | | |
| KR108725.1 | LT169 | *Ormyrus* | *labotus* | 28 | | *Belonocnema kinseyi* | *Quercus virginiana* | Lake Jackson, TX | | |
| OK624770 | P121 | *Ormyrus* | *labotus* | 28 | R. Busbee | *Belonocnema kinseyi* | *Quercus virginiana* | Ingleside, TX | 11/1/2015 | Nov-15 |
| OK624758 | LZ6498 | *Ormyrus* | *labotus tigermorph* | 29 | L. Zhang | *Andricus quercusfoliatus* | *Quercus geminata* | Lithia Springs, FL | 12/16/2018 | 4/8/2019 |
| OK624760 | LZ4717 | *Ormyrus* | *labotus tigermorph* | 29 | L. Zhang | *Andricus quercusfoliatus* | *Quercus virginiana* | Hickory Hammock, FL | 12/15/2018 | 12/26/2018 |

| | | | | | | | | | | |
|---|---|---|---|---|---|---|---|---|---|---|
| OK624771 | P184 | *Ormyrus* | *labotus tigermorph* | 29 | R. Busbee | *Belonocnema treatae* | *Quercus virginiana* | Gautier, MS | 10/16/2015 | Oct-15 |
| OK624772 | P218 | *Ormyrus* | *labotus tigermorph* | 29 | R. Busbee | *Belonocnema fossoria* | *Quercus geminata* | Parker, FL | 10/15/2015 | Oct-15 |
| OK624725 | 920_010_1A | *Ormyrus* | *labotus* | 30 | A. Ward | *Andricus quercuspetiolicola* | *Quercus alba* | Oxford, IA | 6/20/2017 | 6/21/2017 |
| OK624693 | 296_010_5 | *Ormyrus* | *labotus* | 30 | A. Ward | *Andricus quercuspetiolicola* | *Quercus alba* | Iowa City, IA | 6/6/2016 | 6/20/2016 |
| OK624720 | 880_042_17G | *Ormyrus* | *labotus* | 30 | A. Ward | *Andricus quercuspetiolicola* | *Quercus macrocarpa* | Oxford, IA | 6/8/2017 | 6/27/2017 |
| OK624723 | 905_010_008 | *Ormyrus* | *labotus* | 30 | A. Ward | *Andricus quercuspetiolicola* | *Quercus alba* | Iowa City, IA | 6/15/2017 | |
| OK624722 | 884_039_012A | *Ormyrus* | *labotus* | 30 | A. Ward | *Callirhytis seminator* | *Quercus alba* | Iowa City, IA | 6/9/2017 | 6/29/2017 |
| OK624692 | 282_045_2 | *Ormyrus* | *labotus* | 31 | A. Ward | *Amphibolips quercusostensackenii* | *Quercus palustris* | Iowa City, IA | 5/27/2016 | 6/12/2016 |
| OK624719 | 877_045_001A | *Ormyrus* | *labotus* | 31 | A. Ward | *Amphibolips quercusostensackenii* | *Quercus velutina* | Oxford, IA | 6/8/2017 | 6/21/2017 |
| OK624742 | 1367_047_2 | *Ormyrus* | *labotus* | 31 | A. Ward | *Dryocosmus cinereae* | *Quercus imbricaria* | Iowa City, IA | 5/24/2018 | 6/9/2018 |
| OK624714 | 744_047_7B | *Ormyrus* | *labotus* | 31 | A. Ward | *Dryocosmus cinereae* | *Quercus velutina* | Iowa City, IA | 5/9/2017 | 6/5/2017 |
| OK624716 | 857_087_1 | *Ormyrus* | *labotus* | 31 | A. Ward | *Dryocosmus quercusnotha* | *Quercus palustris* | Iowa City, IA | 6/7/2017 | 6/12/2017 |
| OK624741 | 1351_038_4 | *Ormyrus* | *labotus* | 31 | A. Ward | *Dryocosmus quercuspalustris* | *Quercus coccinea* | Oxford, IA | 5/19/2018 | 6/9/2018 |
| OK624706 | 642_009_27 | *Ormyrus* | *labotus* | 32 | A. Ward | *Andricus nigricens* | *Quercus bicolor* | Iowa City, IA | 9/27/2016 | 8/18/2017 |
| OK624711 | 678_006_1 | *Ormyrus* | *labotus* | 32 | A. Ward | *Andricus quercusfrondosus* | *Quercus macrocarpa* | Iowa City, IA | 3/29/2017 | 4/28/2017 |
| OK624691 | 205_029_2 | *Ormyrus* | *labotus* | 32 | A. Ward | *Andricus quercusfrondosus* | *Quercus bicolor* | Iowa City, IA | 4/14/2016 | 5/21/2016 |
| OK624712 | 700_006_5 | *Ormyrus* | *labotus* | 32 | A. Ward | *Andricus quercusfrondosus* | *Quercus bicolor* | Iowa City, IA | 4/12/2017 | 5/4/2017 |
| OK624774 | YMZ1B | *Ormyrus* | *labotus* | 32 | Y. M. Zhang | *Callirhytis quercusclavigera* | *Quercus coccinea* | Gainesville, FL | 11/3/2019 | |
| OK624732 | 1075_100_003 | *Ormyrus* | *labotus* | 32 | A. Ward | *Callirhytis quercusgemmaria* | *Quercus rubra* | Traverse City, MI | 8/13/2017 | 10/25/2017 |
| OK624731 | 1074_100_3 | *Ormyrus* | *labotus* | 32 | A. Ward | *Callirhytis quercusgemmaria* | *Quercus rubra* | Traverse City, MI | 8/13/2017 | 9/4/2017 |
| OK624740 | 1344_123_19B | *Ormyrus* | *labotus* | 32 | A. Ward | *Callirhytis quercusoperator* | *Quercus velutina* | Iowa City, IA | 5/19/2018 | 6/9/2018 |
| OK624727 | 969_016_009H | *Ormyrus* | *labotus* | 32 | A. Ward | *Melikaiella ostensackeni* | *Quercus palustris* | Iowa City, IA | 7/6/2017 | 7/26/2017 |
| OK624695 | 400_016_9B | *Ormyrus* | *labotus* | 32 | A. Ward | *Melikaiella ostensackeni* | *Quercus rubra* | Traverse City, MI | 7/12/2016 | 7/12/2016 |
| OK624726 | 932_070_002 | *Ormyrus* | *labotus* | 33 | A. Ward | *Callirhytis quercuscornigera* | *Quercus palustris* | St. Louis, MO | 6/29/2017 | 7/31/2017 |
| OK624698 | 520_002_1C | *Ormyrus* | *labotus* | 34 | A. Ward | *Acraspis erinacei* | *Quercus alba* | Iowa City, IA | 8/19/2016 | 8/20/2016 |
| OK624699 | 520_002_1D | *Ormyrus* | *labotus* | 34 | A. Ward | *Acraspis erinacei* | *Quercus alba* | Iowa City, IA | 8/19/2016 | 8/20/2016 |
| OK624703 | 601_002_6 | *Ormyrus* | *labotus* | 34 | A. Ward | *Acraspis erinacei* | *Quercus alba* | Urbana, IL | 9/6/2016 | 9/29/2016 |
| OK624710 | 661_002_2A | *Ormyrus* | *labotus* | 34 | A. Ward | *Acraspis erinacei* | *Quercus alba* | Oxford, IA | 10/18/2016 | 5/13/2017 |
| OK624729 | 1028_002_001 | *Ormyrus* | *labotus* | 34 | A. Ward | *Acraspis erinacei* | *Quercus alba* | Iowa City, IA | 7/25/2017 | 7/26/2017 |
| OK624730 | 1028_002_2 | *Ormyrus* | *labotus* | 34 | A. Ward | *Acraspis erinacei* | *Quercus alba* | Iowa City, IA | 7/25/2017 | 8/4/2017 |
| OK624690 | 177_008_3 | *Ormyrus* | *labotus* | 34 | A. Ward | *Andricus quercusflocci* | *Quercus alba* | Oxford, IA | 4/2/2016 | 5/12/2016 |
| OK624718 | 865_051_7A | *Ormyrus* | *labotus* | 34 | A. Ward | *Callirhytis quercusfutilis* | *Quercus alba* | Oxford, IA | 6/8/2017 | 6/28/2017 |
| OK624743 | 1459_051_8 | *Ormyrus* | *labotus* | 34 | A. Ward | *Callirhytis quercusfutilis* | *Quercus alba* | Wyoming, WI | 7/28/2018 | 8/11/2018 |
| OK624766 | JRO_1_5B | *Ormyrus* | *labotus* | 34 | J. Ott | *Philonix nigra* | *Quercus alba* | Capon Bridge, WV | 8/14/2019 | 9/1/2019 |
| OK624701 | 548_002_4B | *Ormyrus* | *labotus* | 35 | A. Ward | *Acraspis erinacei* | *Quercus alba* | Peducah, KY | 9/3/2016 | 9/5/2016 |
| OK624748 | 1566_002_5A | *Ormyrus* | *labotus* | 35 | A. Forbes | *Acraspis erinacei* | *Quercus alba* | Vestal, NY | 10/17/2018 | 6/8/2019 |
| OK624735 | 1181_003_2 | *Ormyrus* | *labotus* | 35 | A. Ward | *Acraspis pezomachoides* | *Quercus alba* | White Oak, PA | 9/9/2017 | 11/12/2017 |
| OK624704 | 602_003_1B | *Ormyrus* | *labotus* | 35 | A. Ward | *Acraspis pezomachoides* | *Quercus alba* | Urbana, IL | 9/6/2016 | 9/10/2016 |
| OK624688 | 58_51_1 | *Ormyrus* | *labotus* | 36 | E. Tvedte | *Acraspis macrocarpae* | *Quercus macrocarpa* | Spirit Lake, IA | 8/1/2015 | 8/12/2015 |
| OK624686 | 58_15_2 | *Ormyrus* | *labotus* | 36 | E. Tvedte | *Acraspis macrocarpae* | *Quercus macrocarpa* | Spirit Lake, IA | 8/1/2015 | 8/8/2015 |
| OK624687 | 58_39_1 | *Ormyrus* | *labotus* | 36 | E. Tvedte | *Acraspis macrocarpae* | *Quercus macrocarpa* | Spirit Lake, IA | 8/1/2015 | 8/6/2015 |
| OK624684 | 1x3 | *Ormyrus* | *labotus* | 36 | A. Forbes | *Acraspis macrocarpae* | *Quercus macrocarpa* | Spirit Lake, IA | 7/13/2015 | 7/17/2015 |
| OK624689 | 58_126_1 | *Ormyrus* | *labotus* | 36 | E. Tvedte | *Acraspis macrocarpae* | *Quercus macrocarpa* | Spirit Lake, IA | 8/1/2015 | 8/6/2015 |
| OK624736 | 1215_080_002 | *Ormyrus* | *labotus* | 36 | A. Ward | *Acraspis prinoides* | *Quercus muehlenbergii* | Urbana, IL | 9/11/2017 | 10/12/2017 |
| OK624707 | 643_004_4 | *Ormyrus* | *labotus* | 36 | A. Ward | *Acraspis villosa* | *Quercus macrocarpa* | Iowa City, IA | 9/27/2016 | 6/21/2017 |
