## Supplemental Table 3 for "*Ormyrus labotus* Walker (Hymenoptera: Ormyridae): another generalist that should not be a generalist is not a generalist"

**Supplemental Table 3.** *Ormyrus* specimens collected in this study and deposited into the University of Iowa Musem of Natural History.

| SUI:INS# | Genus | Species | Order | Family | Determined by | Determined date | Collector | Collection date | Locality | County | State | Country | Region | Notes | Preparation | Sex | lifeStage | Specimen Count |
|---|---|---|---|---|---|---|---|---|---|---|---|---|---|---|---|---|---|---|
| 39302 | Ormyrus | nr. turio | Hymenoptera | Ormyridae | Forbes, Andrew | 5/20/2021 | Anna Ward and Joseph Verry | 2017 July 20 | FW Kent Park | Johnson | IA | United States | North America | 919-013-002; "Clade 1" from Sheikh et al. 2021 Emerged from gall of Callirhytis flavipes 7/1/17 | Pinned | Female | Adult | 1 |
| 39303 | Ormyrus | nr. turio | Hymenoptera | Ormyridae | Forbes, Andrew | 5/20/2021 | Anna Ward and Joseph Verry | 2017 June 8 | FW Kent Park | Johnson | IA | United States | North America | 881-013-006; "Clade 1" from Sheikh et al. 2021 Emerged from gall of Callirhytis flavipes 7/5/17 | Pinned | Female | Adult | 1 |
| 39304 | Ormyrus | venustus | Hymenoptera | Ormyridae | Forbes, Andrew | 5/20/2021 | Anna Ward and Joseph Verry | 2017 June 16 | City Park, Iowa City | Johnson | IA | United States | North America | 908-045-1A; "Clade 4" from Sheikh et al. 2021 Emerged from gall of Amphibolips quercusostensackenii | Pinned | Female | Adult | 1 |
| 39305 | Ormyrus | venustus | Hymenoptera | Ormyridae | Forbes, Andrew | 5/20/2021 | Sofia Sheikh | 2019 September 16 | Show Low | Navajo | AZ | United States | North America | 1664-199-4D; "Clade 7" from from Sheikh et al. 2021 Emerged from gall of Xanthoterus eburnum 11/25/19 | Pinned | Female | Adult | 1 |
| 39306 | Ormyrus | reticulatus | Hymenoptera | Ormyridae | Forbes, Andrew | 5/20/2021 | Anna Ward and William Carr | 2017 April 12 | City Park, Iowa City | Johnson | IA | United States | North America | 701-019-026; "Clade 8" from Sheikh et al. 2021 Emerged from gall of Disholcaspis quercusglobulous (found dead May 2017) | Pinned | Female | Adult | 1 |
| 39307 | Ormyrus | labotus | Hymenoptera | Ormyridae | Forbes, Andrew | 5/20/2021 | Anna Ward and William Carr | 2016 August 18 | City Park, Iowa City | Johnson | IA | United States | North America | 500-011-3C; "Clade 9" from Sheikh et al. 2021 Emerged from gall of Andricus quercusstrobilanus 8/18/16 | Pinned | Female | Adult | 1 |
| 39308 | Ormyrus | unknown | Hymenoptera | Ormyridae | Forbes, Andrew | 5/20/2021 | Miles Zhang | 2019 | Gainesville | Alachua | FL | United States | North America | "Miles1"; "Clade 10" from Sheikh et al. 2021 Emerged from gall of Callirhytis quercusclavigera 11/3/19 | Pinned | Female | Adult | 1 |
| 39309 | Ormyrus | dryorhizoxeni | Hymenoptera | Ormyridae | Davis, Charles | 5/20/2021 | | 2017 December 10 | Inlet Beach | Walton | FL | United States | North America | LZ4112; "Clade 15" from Sheikh et al. 2021 Emerged from gall of Andricus quercusfoliatus 8/27/18 | Pinned | Female | Adult | 1 |
| 39310 | Ormyrus | labotus | Hymenoptera | Ormyridae | Forbes, Andrew | 5/20/2021 | Anna Ward | 2016 September 3 | Bob Noble Park, Peducah | McCraken | KY | United States | North America | 558-081-8A; "Clade 18" from Sheikh et al. 2021 Emerged from gall of Andricus pattoni 9/18/16 | Pinned | Female | Adult | 1 |
| 39311 | Ormyrus | labotus | Hymenoptera | Ormyridae | Forbes, Andrew | 5/20/2021 | Anna Ward | 2016 September 3 | Bob Noble Park, Peducah | McCraken | KY | United States | North America | 558-081-8B; "Clade 18" from Sheikh et al. 2021 Emerged from gall of Andricus pattoni 9/18/16 | Pinned | Male | Adult | 1 |
| 39312 | Ormyrus | labotus | Hymenoptera | Ormyridae | Forbes, Andrew | 5/20/2021 | Alaine Hippee | 2016 September 24 | Konza Prairie | Geary | KS | United States | North America | 631-005-5; "Clade 19" from Sheikh et al. 2021 Emerged from gall of Andricus dimorphus 11/6/2017 | Pinned | Female | Adult | 1 |
| 39313 | Ormyrus | labotus | Hymenoptera | Ormyridae | Forbes, Andrew | 5/20/2021 | Anna Ward | 2017 August 25 | Beaman Park, Nashville | Davidson | TN | United States | North America | 1114-104-007A; "Clade 20" from Sheikh et al. 2021 Emerged from gall of Callirhytis pigra 11/15/17 | Pinned | Male | Adult | 1 |
| 39314 | Ormyrus | labotus | Hymenoptera | Ormyridae | Forbes, Andrew | 5/20/2021 | Andrew Forbes | 2018 October 17 | Vestal | Broome | NY | United States | North America | 1574-104-9A; "Clade 22" from Sheikh et al. 2021 Emerged from gall of Callirhytis pigra 5/24/19 | Pinned | Female | Adult | 1 |
| 39315 | Ormyrus | labotus | Hymenoptera | Ormyridae | Forbes, Andrew | 5/20/2021 | Anna Ward and Leo Gastel | 2018 May 16 | City Park, Iowa City | Johnson | IA | United States | North America | 1335-097-2; "Clade 24" from Sheikh et al. 2021 Emerged from an unidentified raised vein gall on pin oak 6/19/18 | Pinned | Male | Adult | 1 |
| 39316 | Ormyrus | labotus | Hymenoptera | Ormyridae | Forbes, Andrew | 5/20/2021 | Anna Ward and Daniel McGarry | 2017 August 8 | Mormon Handcart Trail, Coralville | Johnson | IA | United States | North America | 1059-024-3; "Clade 25" from Sheikh et al. 2021 Emerged from gall of Phylloterus pocoulum 9/4/17 | Pinned | Female | Adult | 1 |
| 39317 | Ormyrus | labotus | Hymenoptera | Ormyridae | Forbes, Andrew | 5/20/2021 | Anna Ward | 2018 June 5 | Coralville | Johnson | IA | United States | North America | 1384-30-1B; "Clade 25" from Sheikh et al. 2021 Emerged from gall of Andricus chinquapin 10/29/18 | Pinned | Female | Adult | 1 |
| 39318 | Ormyrus | labotus | Hymenoptera | Ormyridae | Forbes, Andrew | 5/20/2021 | Andrew Forbes | 2018 April 12 | McKinney Roughs Nature Park | Travis | TX | United States | North America | 1238-117-12; "Clade 26" from Sheikh et al. 2021 Emerged from gall of Andricus quercuspetiolicola 9/21/18 | Pinned | Male | Adult | 1 |
| 39319 | Ormyrus | labotus | Hymenoptera | Ormyridae | Forbes, Andrew | 5/20/2021 | Miles Zhang | 2019 | Gainesville | Alachua | FL | United States | North America | "Miles-3"; "Clade 26" from Sheikh et al. 2021 Emerged from gall of Dryocosmus sp. 3/20/20 | Pinned | Female | Adult | 1 |
| 39320 | Ormyrus | labotus | Hymenoptera | Ormyridae | Forbes, Andrew | 5/20/2021 | Anna Ward | 2018 November 1 | Kyle | Hays | TX | United States | North America | 1589-160-2A; "Clade 27" from Sheikh et al. 2021 Emerged from gall of Andricus quercuslanigera 11/12/18 | Pinned | Female | Adult | 1 |
| 39321 | Ormyrus | labotus | Hymenoptera | Ormyridae | Forbes, Andrew | 5/20/2021 | Anna Ward | 2018 November 1 | Kyle | Hays | TX | United States | North America | 1589-160-4; "Clade 27" from Sheikh et al. 2021 Emerged from gall of Andricus quercuslanigera (found dead 3/22/18) | Pinned | Female | Adult | 1 |
| 39322 | Ormyrus | labotus | Hymenoptera | Ormyridae | Forbes, Andrew | 5/20/2021 | Scott Egan | 2014 October 27 | Lake Jackson | Brazoria | TX | United States | North America | LT_169; "Clade 28" from Sheikh et al. 2021 Emerged from gall of Belonocnema kinseyi 11/12/14 | Pinned | Female | Adult | 1 |
| 39323 | Ormyrus | labotus | Hymenoptera | Ormyridae | Forbes, Andrew | 5/20/2021 | Scott Egan | 2015 | Sapelo Island | McIntosh | GA | United States | North America | P172; "Clade 29" from Sheikh et al. 2021 Emerged from gall of Belonocnema treatae 10/2015 | Pinned | Female | Adult | 1 |
| 39324 | Ormyrus | labotus | Hymenoptera | Ormyridae | Forbes, Andrew | 5/20/2021 | Anna Ward and Joseph Verry | 2017 June 8 | FW Kent Park | Johnson | IA | United States | North America | 880-042-17F; "Clade 30" from Sheikh et al. 2021 Emerged from gall of Neuroterus quercusbatatus 6/27/17 | Pinned | Female | Adult | 1 |
| 39325 | Ormyrus | labotus | Hymenoptera | Ormyridae | Forbes, Andrew | 5/20/2021 | Anna Ward and Joseph Verry | 2017 June 9 | U.Iowa Campus | Johnson | IA | United States | North America | 884-039-10A; "Clade 30" from Sheikh et al. 2021 Emerged from gall of Callirhytis seminator 7/4/17 | Pinned | Female | Adult | 1 |
| 39326 | Ormyrus | labotus | Hymenoptera | Ormyridae | Forbes, Andrew | 5/20/2021 | Anna Ward, Daniel McGarry, and Joseph Verry | 2017 June 7 | City Park, Iowa City | Johnson | IA | United States | North America | 857-087-002; "Clade 31" from Sheikh et al. 2021 Emerged from gall of Dryocosmus quercusnotha 6/16/17 | Pinned | Female | Adult | 1 |
| 39327 | Ormyrus | labotus | Hymenoptera | Ormyridae | Forbes, Andrew | 5/20/2021 | Anna Ward and Kyle McElroy | 2018 May 19 | FW Kent Park | Johnson | IA | United States | North America | 1344-123-26A; "Clade 32" from Sheikh et al. 2021 Emerged from gall of Callirhytis quercusoperator 6/13/18 | Pinned | Female | Adult | 1 |
| 39328 | Ormyrus | labotus | Hymenoptera | Ormyridae | Forbes, Andrew | 5/20/2021 | Anna Ward and Kyle McElroy | 2018 May 19 | FW Kent Park | Johnson | IA | United States | North America | 1344-123-25B; "Clade 32" from Sheikh et al. 2021 Emerged from gall of Callirhytis quercusoperator 6/13/18 | Pinned | Female | Adult | 1 |
| 39329 | Ormyrus | labotus | Hymenoptera | Ormyridae | Forbes, Andrew | 5/20/2021 | Anna Ward | 2016 July 12 | Traverse City | Grand Traverse | MI | United States | North America | 400-016-10; "Clade 32" from Sheikh et al. 2021 Emerged from gall of Melikaiella ostensackeni 8/7/16 | Pinned | Female | Adult | 1 |
| 39330 | Ormyrus | labotus | Hymenoptera | Ormyridae | Forbes, Andrew | 5/20/2021 | Anna Ward and Daniel McGarry | 2017 July 6 | City Park, Iowa City | Johnson | IA | United States | North America | 969-016-7C; "Clade 32" from Sheikh et al. 2021 Emerged from gall of Melikaiella ostensackeni 7/21/17 | Pinned | Female | Adult | 1 |
| 39331 | Ormyrus | labotus | Hymenoptera | Ormyridae | Forbes, Andrew | 5/20/2021 | Anna Ward | 2017 June 3 | Creve Coeur Lake | St. Louis | MO | United States | North America | 837-070-29A; "Clade 33" from Sheikh et al. 2021 Emerged from gall of Callirhytis quercuscornigera 7/2/17 | Pinned | Female | Adult | 1 |
| 39332 | Ormyrus | labotus | Hymenoptera | Ormyridae | Forbes, Andrew | 5/20/2021 | Anna Ward, Robin Bagley, and Sarah DeLong-Duhon | 2018 July 28 | Dodgeville | Iowa | WI | United States | North America | 1450-051-5; "Clade 34" from Sheikh et al. 2021 Emerged from gall of Callirhytis quercusfutilis 8/19/18 | Pinned | Female | Adult | 1 |
| 39333 | Ormyrus | labotus | Hymenoptera | Ormyridae | Forbes, Andrew | 5/20/2021 | Anna Ward, Robin Bagley, and Sarah DeLong-Duhon | 2018 July 28 | Spring Green | Sauk | WI | United States | North America | 1459-051-6; "Clade 34" from Sheikh et al. 2021 Emerged from gall of Callirhytis quercusfutilis 7/28/18 | Pinned | Female | Adult | 1 |
| 39334 | Ormyrus | labotus | Hymenoptera | Ormyridae | Forbes, Andrew | 5/20/2021 | Anna Ward, Daniel McGarry, and Joseph Verry | 2017 June 15 | Hickory Hills Park | Johnson | IA | United States | North America | 899-051-008; "Clade 34" from Sheikh et al. 2021 Emerged from gall of Callirhytis quercusfutilis 6/27/17 | Pinned | Female | Adult | 1 |
| 39335 | Ormyrus | labotus | Hymenoptera | Ormyridae | Forbes, Andrew | 5/20/2021 | Anna Ward and Joseph Verry | 2017 June 8 | FW Kent Park | Johnson | IA | United States | North America | 865-051-009; "Clade 34" from Sheikh et al. 2021 Emerged from gall of Callirhytis quercusfutilis 6/30/17 | Pinned | Female | Adult | 1 |
| 39336 | Ormyrus | labotus | Hymenoptera | Ormyridae | Forbes, Andrew | 5/20/2021 | Jim Ott | 2019 August | Capon Bridge | Hampshire | WV | United States | North America | JRO-1-JA; "Clade 34" from Sheikh et al. 2021 Emerged from gall of Philonix nigra 9/1/19 | Pinned | Female | Adult | 1 |
| 39337 | Ormyrus | labotus | Hymenoptera | Ormyridae | Forbes, Andrew | 5/20/2021 | Anna Ward, Robin Bagley, and Sofia Sheikh | 2018 October 6 | McGregor | Clayton | IA | United States | North America | 1561-002-1A; "Clade 35" from Sheikh et al. 2021 Emerged from gall of Acraspis erinacei 5/23/19 | Pinned | Female | Adult | 1 |
| 39338 | Ormyrus | labotus | Hymenoptera | Ormyridae | Forbes, Andrew | 5/20/2021 | Andrew Forbes | 2018 October 17 | Vestal | Broome | NY | United States | North America | 1566-002-5B; "Clade 35" from Sheikh et al. 2021 Emerged from gall of Acraspis erinaccei 6/8/19 | Pinned | Female | Adult | 1 |
| 39339 | Ormyrus | labotus | Hymenoptera | Ormyridae | Forbes, Andrew | 5/20/2021 | Anna Ward and Caleb Wilson | 2016 July 27 | Coralville | Johnson | IA | United States | North America | 428-001-003; "Clade 36" from Sheikh et al. 2021 Emerged from gall of Acraspis macrocarpae 8/11/16 | Pinned | Female | Adult | 1 |
| 39340 | Ormyrus | labotus | Hymenoptera | Ormyridae | Forbes, Andrew | 5/20/2021 | Andrew Forbes | 2015 July 13 | Spirit Lake | Dickinson | IA | United States | North America | 1-X-4; "Clade 36" from Sheikh et al. 2021 Emerged from gall of Acraspis macrocarpae 7/18/15 | Pinned | Female | Adult | 1 |
| 39341 | Ormyrus | labotus | Hymenoptera | Ormyridae | Forbes, Andrew | 5/20/2021 | Kelly Weinersmith | 2019 March | | | FL | United States | North America | 174; "Clade 2" from Sheikh et al. 2021 Emerged from gall of Bassetia pallida 4/1/19 | Pinned | Female | Adult | 1 |
